# Carbohydrate active enzymes of giant viruses: Molecular and biochemical characterization of glycosyl hydrolases from algae-infecting chloroviruses

**DOI:** 10.64898/2026.07.15.738767

**Authors:** Ellen G. Oliveira, Pavla Fajtová, Mateus S. M. Serafim, Sara A. C. Souza, Clécio A. C. Filho, João Victor R. P. Carvalho, Álan N. P. Gomes, Daniel A. Santos, Matheus A. Motta, Lucas Bleicher, Ronaldo A. P. Nagem, Anthony J. O’Donoghue, Rodrigo A. L. Rodrigues

## Abstract

Microbial hydrolases are considered to be promising enzymes for pathogen control. Bacterial and viral chitinases of the glycosyl hydrolase (GH) 18 family are important biological macromolecules with antifungal and anti-insect activity. Chloroviruses, nucleocytoplasmic large DNA viruses (NCLDVs) that infect unicellular green algae have a considerable number of genes involved in carbohydrate metabolism, including the chitinase GH18 family. In this study, we investigated the abundance and diversity of chitinases in chlorovirus genomes using a combination of *silico* and *in vitro* strategies, and characterized these enzymes at a molecular and biochemical level. Different enzymatic profiles were observed in *Chlorovirus* subgenera revealing the different viral machinery related to host species. We performed a comprehensive biochemical characterization of three heterologous expressed GH18 domains, which revealed their endo and exochitinase activity and thermostability. Crystallographic analysis of the GH18 domain by X-ray diffraction yielded a structure at 1.0 Å resolution, representing the highest-resolution structure reported to date for a giant viral protein and showing lower predominancy of residue coevolution compared GH18 chitinases from other organisms. Additionally, our binding site characterization predicted high conservation in betachloroviruses and gammachloroviruses, and less so in alphachloroviruses. Lastly, these enzymes did not inhibit fungal growth of medical and agricultural importance species *in vitro* but exhibited high inhibitory activity against different algae at nanogram/mL range. Together, our experimental and computational data show that evolutionary events may contribute to maintaining viral chitinases’ enzymatic activity and specificity. These findings highlight the potential of virus-derived enzymes as promising new biotechnological tools for microbial control against different algal strains.

## INTRODUCTION

Global demand for industrial enzymes is continuously growing and is estimated to reach US$7 billion in the coming years ^1^. In this context, microbial enzymes offer several advantages such as high yield, use in low-cost platforms, ease of acquisition, and optimization. Approximately 60% of industrial enzymes are obtained from fungi, ∼24% from bacteria, ∼4% from yeasts, and the remainder from plants and animals ^2,3^. While viral enzymes represent small percentage of the total industrial enzyme market, biotechnology relies heavily on them for genetic engineering, DNA and RNA regulation mechanisms, such as endolysins and hydrolases with potential antimicrobial action ^4,5^.

Nucleocytoplasmic large DNA viruses (NCLDV), commonly referred to as giant viruses, carry large genomes (>200 kbp), with a diversity of genes capable of encoding different proteins ^6^. Chloroviruses belong to the family *Phycodnaviridae* and infect the unicellular green algae of the family Chlorellaceae. They are currently classified in three subgenera based on genomic and phylogenomic features, and also related to the specificity to the host species ^7^. *Alphachlorovirus* classifies the viruses that infect *Chlorella variabilis*, including viruses capable of infecting the NC64A strain and those capable of infecting only the Syngen 2-3 strain, called Osy-viruses; *Betachlorovirus* comprises the viruses that infect *Micractinium conductrix* Pbi; and *Gammachlorovirus* includes those that infect *Chlorella heliozoae* SAG3.83 ^8–10^. There viruses have a diverse set of genes involved in nucleotide metabolism, DNA transcription, RNA translation and carbohydrate metabolism, such as chitinase glycosyl hydrolase (GH) 18 family (EC 3.2.1.14). These GH18 are hydrolytic enzymes that break chitin chains of *N*-acetyl d-glucosamine in the (1→4) β-glycoside region, acting as a catalysis-assisted substrate ^11^.

Viral and bacterial GH18 chitinases have been shown as important biological macromolecules with antifungal activity and pest insect control, thus promising for microbial control ^12–16^. In this context, such biomolecules have greater advantages compared to synthetic molecules, often associated with mechanisms with high specificity, presenting less environmental impact and reduced risks to human health ^17^. Commonly present in baculoviruses and phages, GH18 chitinases are an important and unexplored source of biomolecules with biotechnological potential in chloroviruses ^15,18^. In this study we demonstrate the complexity of GH18 chitinases in 128 complete genomes of chloroviruses spanning all subgenera, revealing evolutionary factors that lead to their structural and biochemical diversity. We performed a comprehensive biochemical characterization of three heterologous expressed GH18 domains, which revealed their endo and exochitinase activity and thermostability. Crystallographic analysis of the GH18 domain by X-ray diffraction yielded a structure at 1.04 Å resolution, showing lower predominancy of residue coevolution compared GH18 chitinases from other organisms. Additionally, our binding site characterization predicted high conservation in betachloroviruses and gammachloroviruses, and less so in alphachloroviruses. Moreover, we demonstrated the biological activity of these enzymes against different algal strains at nanogram range, supporting its potential as biotechnological biomolecules.

## RESULTS AND DISCUSSION

### Genome analyses reveal the abundance and diversity of carbohydrate active enzymes in chloroviruses

Genes involved in sugar metabolism, glycosylation, and cell wall modulation are present in chloroviruses genomes, being a notable source of carbohydrate active enzymes (CAZymes). We evaluated the diversity and abundance of these genes in 89 chloroviruses genomes. A high variability was identified in chloroviruses that infect the same host species (Figure 1A). In chloroviruses that infect *Chlorella variabilis* NC64A, the maximum number of genes was n=25 and in those infecting *Chlorella variabilis* Syngen 2-3, n=23. In *Betachlorovirus* and *Gammachlorovirus*, we identified 18 and 27 genes, respectively. Comparing between groups, the average number of carbohydrate-modulating genes is higher in the OSy-viruses (*Alphachlorovirus*) and in *Gammachlorovirus* (Figure 1A). Chloroviruses have a considerable number of genes involved in sugar and carbohydrates modulation mechanisms, such as those involved in glycosylation and in the biosynthesis and degradation of polysaccharides ^19–21^,some of them are also found in other giant viruses ^22,23^. We identified genes involved in the biosynthesis and modification of sugars, such as GDP-L-fucose synthase-2, GDP-D-mannose dehydratase, glucosamine-6-phosphate aminotransferase, acetyltransferases, and D-lactate dehydrogenase, as well as enzymes involved in the modification of polysaccharide chains, such as polysaccharide deacetylases and glycosyltransferases. In addition, we observed a diversity of genes in the analyzed genomes involved in the biosynthesis, modification, and degradation of different structural and extracellular polysaccharides, including cellulose, chitin, β-glucans, alginate, and hyaluronan (Figure 1B).

**Figure 1.**
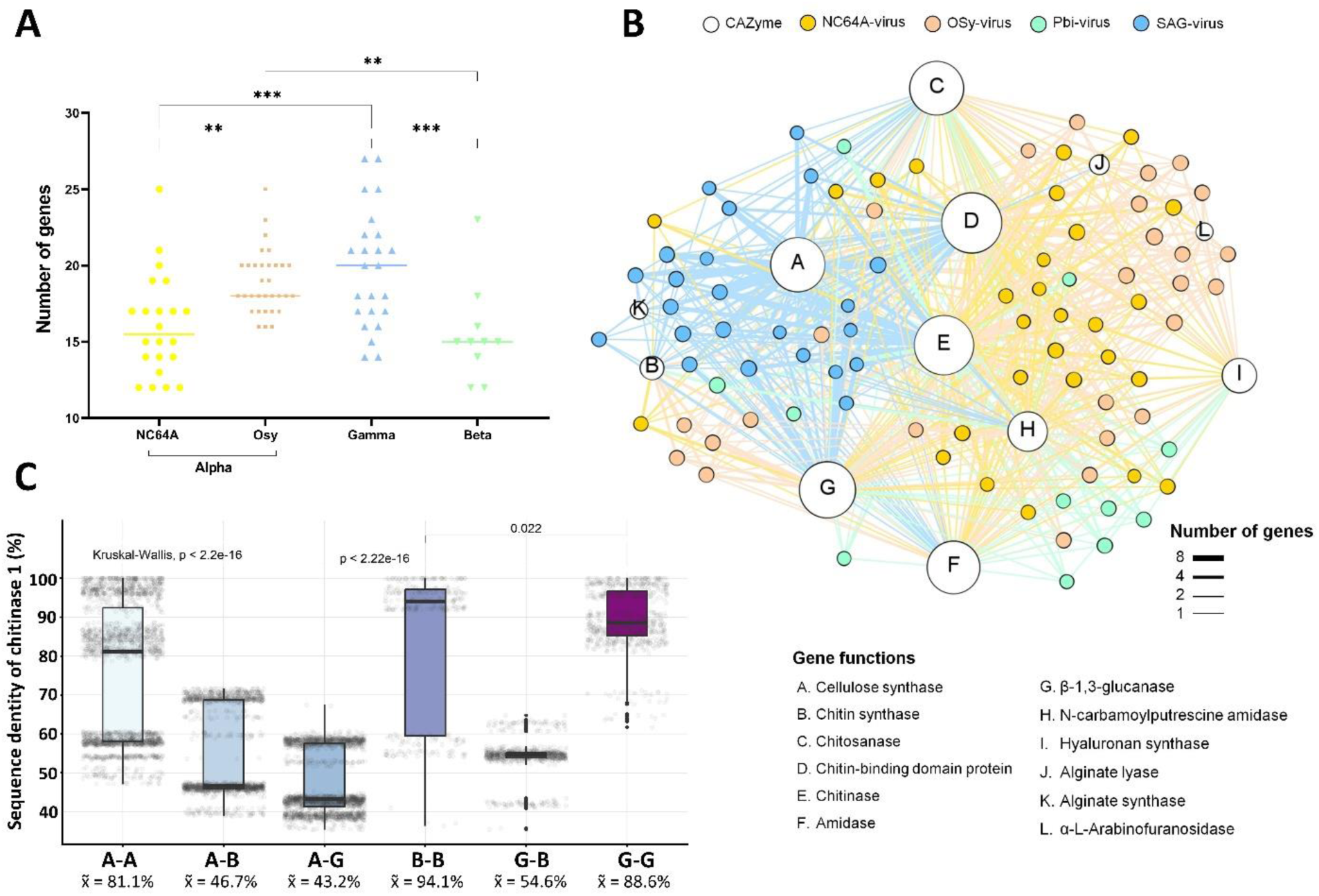
Abundance and diversity of CAZymes in chloroviruses genomes. **A:** Absolute number of genes involved in carbohydrate and sugar metabolism in *Alpha-* (NC64A-and Osy-viruses), *Gamma-* and *Betachloroviruses* that infect different algae hosts, based on the gene annotation of 89 viral genomes. One-way ANOVA statistical test was performed. *p*-value < 0.0001. **B:** Network representation of CAZymes present in the 89 analyzed chlorovirus genomes. Line thickness represents the number of genes (one, two, four, or eight) connecting two or more chloroviruses by different gene functions (A to L, *e.g.*, cellulose synthase). Colored circles represent NC64A-viruses (bright orange), OSy-viruses (salmon), Pbi-viruses (aquamarine), and SAG-viruses (sky blue), or CAZymes (white). **C:** Pairwise comparison of amino acid sequence identity of chitinase 1 gene found in 128 genomes of chlorovirus isolates from all three subgenera. Each gray dot represents a percentage value of nucleotide identity obtained from the alignment of two different sequences. A-A: Alpha-Alpha, A-B: Alpha-Beta, A-G: Alpha-Gamma, B-B: Beta-Beta, G-G: Gamma-Gamma. x̃: median, Kruskal-Wallis non-normal distribution statistical analysis was performed, with the highest *p*-value < 2.2e^-16^ (*p*-values < 0.05 indicate differences between the medians of the groups). The p-values shown in the bars were obtained from the Wilcoxon analysis (*p*-values < 0.05 and the absence of numbers indicate that there are differences between the medians of the groups).

Isolates 41-NE-6 and O-NE-20 of *Alphachlorovirus* subgenus have higher number of genes encoding glycosyl transferases and chitinase-binding domains. In contrast, cellulose synthase and (1-3)-beta-glucanase are more abundant in gammachloroviruses (e.g.: isolate CL-S-1-m). Some genes such as hyaluronan synthase are less abundant among the subgenus and are absent in gammachlorovirus. The chitin synthase genes are highly abundant in OSy-viruses and gammachloroviruses, just as the cellulose synthase in gammachloroviruses. The alginate synthase gene was more frequently found in gammachloroviruses, while alginate lyase, on the other hand, is more prevalent in alphachloroviruses, being almost absent in the other chloroviruses’ groups. Curiously, the alpha-L-arabinofuranosidase gene was found exclusively in the OSy-viruses (Figure 1B and Table S1). Although the factors of specificity of these viruses with their hosts are not elucidated, it is interesting to note that the predominance of some carbohydrate-modulating genes is different in the viral clades and might be related to host specificity. We identified a higher number of cellulose synthase copies in gammachloroviruses, the presence of the alpha-L-arabinofuranosidase gene only in alphachloroviruses, the chitin synthase gene in gammachloroviruses and betachloroviruses, and the major presence of alginate lyase in alphachloroviruses (Figure 1B). Altogether, these findings may indicate the need of different enzymatic systems for the full entry and replication of the virus in the host.

The genes involved in polysaccharide degradation, like chitosanase, chitinase, beta-(1,3)-glucanase, and amidase, are the most abundant across all clades. Enzymes such as amidase and alginate lyase are present in the viral particle and may be involved in the early phase of viral infection, assisting viral particles in digesting the rigid cell wall of *Chlorella* ^24^. Although alginate lyase is not present in all chlorovirus genomes, this protein is known to be present in the viral particle of PBCV-1, the most studied chlorovirus to date ^25^. This enzyme is also related to the degradation of alginate, a polymer composed of α-L-guluronate and C5 epimer β-D-mannuronate, also present in algal cell walls ^26^. Further, exposure to the enzyme alginate lyase, encoded by the a*561l* gene, allowed for greater specificity and increased of infection of viral particle, demonstrating an important function in the early phase ^24,25,27^.

Among the CAZymes, chitinases belonging to the GH18 family (EC 3.2.1.14) are highly abundant in chloroviruses genomes. Thus, given their implications in algae biology and potential for biotechnology application, we focused on further analysis of these genes. GH18 chitinases are named in this work according to the motif found in their active site. Genes having two aspartates and one glutamate (DxDxE) as the catalytic residues are hereafter referred to as chitinase 1, which were detected in all chloroviruses to date, and those that have only one aspartate and one glutamate (DxE) are referred to as chitinase 2.

Chitinase 1 exhibits substantial variation in amino acid sequence identity among viruses of the same subgenus, ranging from 50 to 100% (median 81.1%) in alphachlorovirus, 60 to 100% (median 88.6%) in gammachlorovirus, and 55 to 100% (median 94.1%) in betachlorovirus (Figure 1C). When sequences between different subgenera are compared, this similarity drops to between 65.8 to 71.6%. This pattern is also observed for chitinase 2 (Figure S1A). In contrast, comparing the identity of a conserved topoisomerase gene, we observed high similarity and lower variability, with median values greater than 91% (Figure S1B). These findings indicate higher variability of chitinase 1 comparing to topoisomerase and chitinase 2 in the genomes.

Chitinase 1 has a type II carbohydrate-binding module (CBM2) linked to the first GH18 domain that consists of a DFDIE motif. The presence of phenylalanine between the two aspartates in the catalytic site is found in all chloroviruses genomes. The second GH18 domain (termed GH18 domain 2) has the DMDIE motif. Methionine, located between the two aspartates, is only present in alphachloroviruses, particularly *C. alphanebraskense*, *C. syngense*, and *C. primosyngense*. In betachloroviruses, this second GH18 domain consists of DLDIE motif instead, and is present in only two isolates, FR483 and NW665.2 (Figure S1C). Taking the PBCV-1 (reference virus) for comparison with chitinase 1 sequences, the overall identity ranged from 82 to 100% in alphachloroviruses, 58 to 71% in betachloroviruses, and 57 to 71% in gammachloroviruses. The first GH18 domain shows between 72.63 and 80.79% identity among viral groups, and the second GH18 domain is less conserved, showing an overall lower sequence identity compared to the first domain, being 44.54% and 35.68% for alphachloroviruses and betachloroviruses, respectively (Figure S1C).

Other giant viruses that infect amoebas instead of algae, such as Marseillevirus LCMAC201, Tupanvirus soda lake, Tupanvirus deep ocean, Homavirus sp., Satyrvirus sp., and Fadolivirus, have genes that encode chitinases GH18. However enzymes involved in cell wall modulation are more frequently found in chloroviruses ^28^. Most chitinase genes seem to have been transferred to chloroviruses from their hosts prior the divergence of different viral clades (Figure S2), contrary to previous hypothesis of gene acquisition through bacteria ^28^. However, it is interesting that some gammachlorovirus chitinases might have a different origin, possibly coming from prokaryotes (Figure S2). Our findings suggest that multiple origin scenario might be the case for chitinases in chloroviruses, and a more in-depth evolutionary analysis is required. Interestingly, the GH18 domain of chitinase 1 shows approximately 31% identity to an enzyme encoded by the archaea *Pyrococcus furiosus*, while chitinase 2 shows 35-38% similarity to a chitinase of the fungus *Clonostachys rosea*. Similar enzymes are also found in bacteria that infect different *Chlorella*, such as *Vampirovibrio chlorellavorus*, a cyanobacterium that infect *Chlorella vulgaris* ^29^. *V. chlorellavorus* requires hydrolytic enzymes such as endoglucanase, chitinase GH19, alginate lyase, and beta-galactosidase to inject itself into the cytoplasm and digest the cellular contents, thereby promoting cell death ^29^. These mechanisms demonstrate that the acquisition of these enzymes into microbial genomes highlights their biological interaction with Chlorella-like hosts. To maintain successful infection, they need to keep these genes in their genomes. This mechanism in chloroviruses is not yet well elucidated and, therefore, is necessary not only for understanding the biology of viral infection but also for a better understanding of the function of these enzymes.

Transcript levels and hydrolytic activity of chitinases have been previously described for the chlorovirus CVK-2, which infects *Chlorella variabilis* NC64A, providing evidence that they are expressed during viral cycle and can have a comprehensive role in cell wall degradation ^30^. However, transcriptome and proteome analyses of PBCV-1 revealed that these proteins are absent in the viral particle, suggesting expression in the late phase of infection ^25,31^. Chitinases have a major role in the viral cycle of baculoviruses, being responsible for terminal liquefaction of host insect corpses and release of occlusion bodies with viral particles, an essential mechanism in the release of these viruses ^32^. Interestingly, this enzyme is also found in the endoplasmic reticulum of insects, where the final assembly of viral particles normally occurs ^33^. Thus, similarly to baculoviruses, chloroviruses could also use their chitinases as important enzymes for cell wall degradation in the final phase of infection, when viral particles are released from the host cells.

### BINDING SITE CHARACTERIZATION OF CHLOROVIRUS GH18 CHITINASES REVEAL PARTICULARITIES IN ENZYME DOMAINS

Given the universal presence of chitinase 1 in chloroviruses, we focused our efforts on characterizing this enzyme in terms of biochemical and physiological properties, considering the two distinct GH18 domains. First, our binding site characterizations using PrankWeb 3 predicted two potential sites in the first domain (Figure 2). The first is composed of only six amino acid residues in alphachlorovirus, four of them being polar residues. Of note, this same site is composed of 23 residues in gammachlorovirus, and 25 in betachlorovirus (Figure 2). Although rich in aspartate, they showed a greater diversity of nonpolar residues compared to alphachlorovirus, e.g., high frequency of alanine in betachlorovirus. The second predicted binding site is composed of 10 residues in alphachlorovirus, eight of which are polar. Conversely, seven residues were identified in gammachlorovirus, only two of which are polar (Figure 2). In gamma-and betachlorovirus, we observed a higher conservation of the predicted sites, showing no residue substitutions among the genomes of the different subgenera analyzed. Importantly, different substitutions were found in three different viral species of alphachlorovirus.

**Figure 2.**
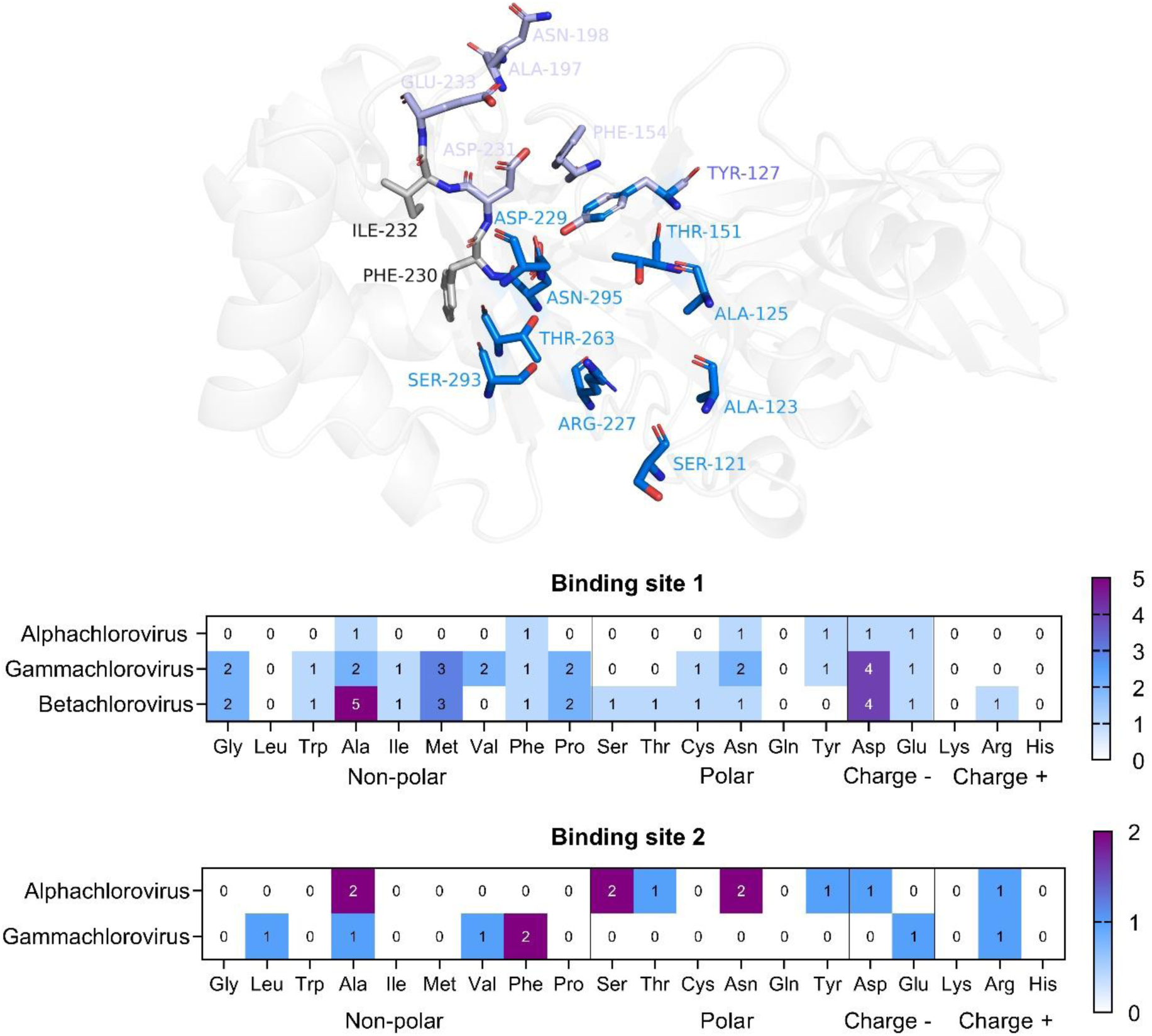
Frequency of residues predicted in the potential binding sites of GH18 domain 1 in all subgenera of *Chlorovirus.* Three-dimensional structure of PBCV-1 chitinase domain 1 (gray) modeled by AlphaFold2 is shown as transparent cartoon with the potential binding sites 1 (purple) and 2 (blue), represented by surface predicted by PrankWeb 3. Ile and Phe resideus from catalytic motif DFDIE are shown in gray. Absolute numbers (reference bars, 0 to 5, and 0 to 2) of each amino acid residue predicted in each binding site of each *Chlorovirus* subgenera (*Alpha*-, *Beta*-, and *Gamma*-) are compiled in the heatmap. Residues are separated as polar, nonpolar, charged negatively (-) or positively (+).

In *Alphachlorovirus*, substitutions in the predicted binding sites are present in viruses that comprise species *Chlorovirus americanus*, *C. alphanebraskense*, and *C. primosyngense*. Interestingly, all viruses that have substitutions in domain 1 also have substitutions it in domain 2. For instance, all viruses of species *C. americanus* (chlorovirus isolates WNE-10B-S1, MA-1D, NYs1, NY-2B, IL-5-2s1, AR158, and NY2A)^7^, have an Ala123Val and Asn259Glu in domain 1, and Gly19Cys, Gly161Glu in domain 2 (Figure 3A, B). In five of the 11 viruses of species *C. alphanebraskense* (N-NE-5, NE-40-2S, 41-NE-4, NE-41-3-S, and 40-NE-5), an Asn259Thr occur in domain 1, and Gln19Gly and Ser159Thr in domain 2 (Figure 3C, D). As for species *C. syngense*, composed of eight OSy-viruses, Ser121Asn is observed in domain 1 for six viruses (O-NE-19, O-NE-22, O-NE-29, O-NE-27, O-NE-23, and NE-O-7-S). Last, in domain 2, there is an Ala152Serin the same seven viruses (O-NE-19, O-NE-22, O-NE-29, O-NE-27, O-NE-25, O-NE-7-S, and O-NE-23), except for O-NE-18 (Figure 3E, F). These data indicate that the amino acid composition may be related to the degree of conservation of sites in this domain, where the predominance of polar residues is associated with increased variability and the predominance of nonpolar residues with the conservation of the binding site. Finally, only one binding site was predicted in domain 2 for alphachloroviruses, with 25 residues, and in two isolates of betachloroviruses, with 21 residues. Both domains have a high prevalence of aspartate, a polar amino acid, with similar distributions of polar and nonpolar amino acids (Figure S3). In *Betachlorovirus*, the site is also conserved, and in *Alphachlorovirus*, substitutions associated with species *C. americanus*, *C. alphanebraskense*, and *C. syngense* were also found. For this domain, there was no correlation between amino acid predominance and conserved amino acid positions.

**Figure 3.**
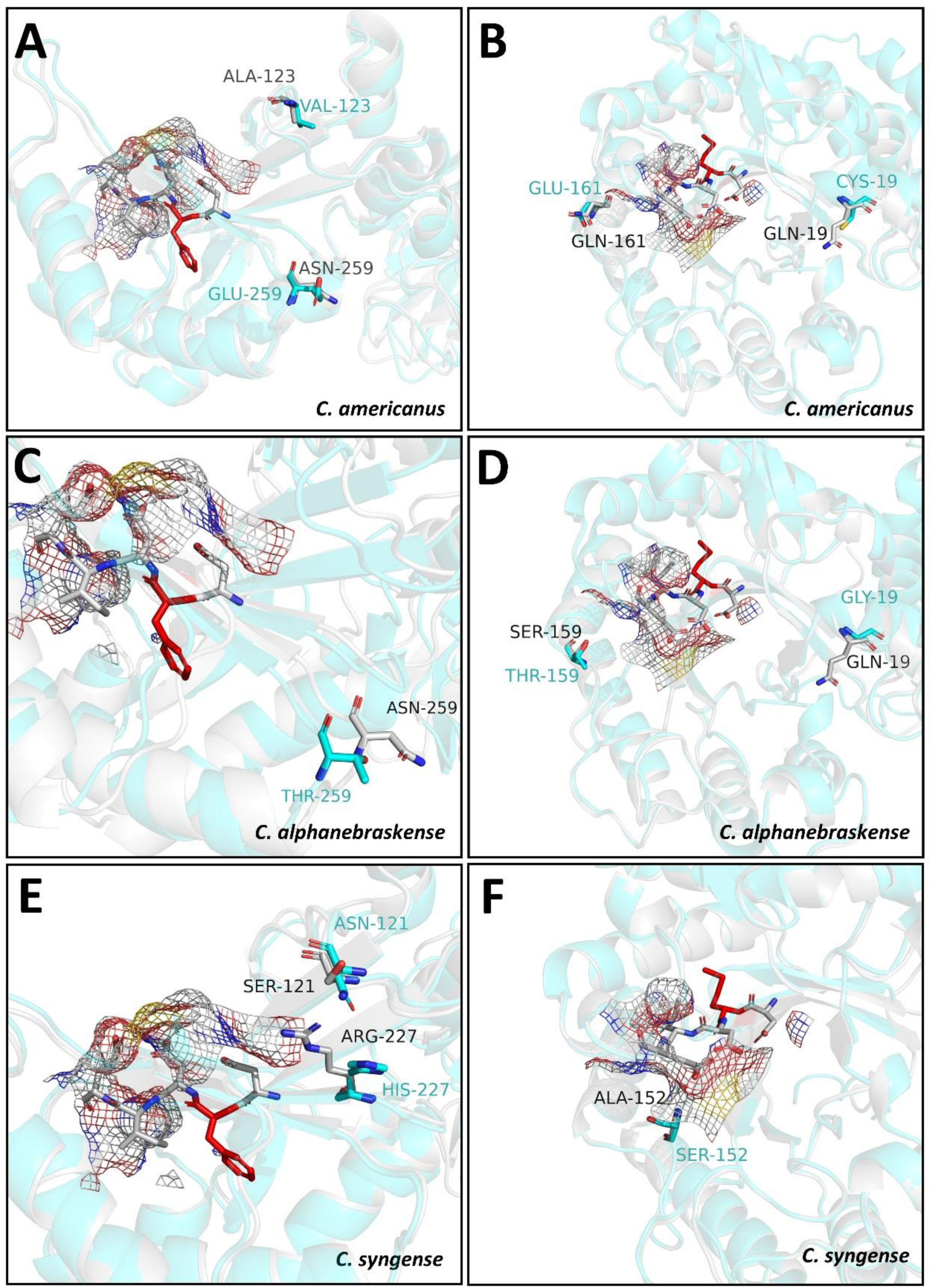
Residue substitutions within predicted binding sites of GH18 domains in *Alphachlorovirus*. Residue substitutions in GH18 domains 1 and 2 of of species *C. americanus*, *C. alphanebraskense*, and *C. syngense* modeled by AlphaFold2 (cyan). A, C, and E: substitutions of domain 1 present in chitinases of species *C. americanus*, *C. alphanebraskense*, and *C. syngense*, respectively. B, D, and F: substitutions of domain 2 present in species *C. americanus*, *C. alphanebraskense*, and *C. syngense*. Structures are superimposed to the three-dimensional structure of the modeled PBCV-1 domains (gray). Conserved or substituted residues are highlighted as sticks and labeled. Colored mesh corresponds to the predicted binding sites by PrankWeb 3.

The predominance of nonpolar residues in the binding sites predicted for domain 1 of *Betachlorovirus* and *Gammachlorovirus* can be associated with the conservation of these sites in viruses of these subgenera. Such nonpolar residues may be related to the stabilization of the enzyme tertiary structures ^34^. On the other hand, in the binding sites of domain 1 in *Alphachlorovirus* there is a prevalence of polar residues, having important substitutions that characterize species *C. americanus*, *C. alphanebraskense*, and *C. syngense*. This may be associated with the protein adaptation to possible interactions in the external environment, such as solubility and pH ^35,36^. Local environments or pockets formed by residues in the tertiary structure of the protein influence the probability of substitutions in evolutionary mechanisms, which may support their occurrence in the two GH18 domains of the three species in *Alphachlorovirus* ^37^. Thus, it is possible that these viruses need to maintain the activity of these proteins in cellular compartments with distinct biochemical characteristics, also correlating to their host specificity.

Common genes to all chloroviruses, such as chitinases 1, also exhibit important structural and biochemical variations that can modulate this specificity and indicate viral resistance to biophysical factors. The evolutionary selective pressure on domain 1 appears to be greater in all chloroviruses, due to the conservation of its residues, ultimately interfering with the maintenance of its enzymatic activity ^38^. The presence of the second GH18 domain, predominantly found in *Alphachlorovirus* and only two betachloroviruses so far, can be the result of a genomic duplication, which differentiated from the first domain, as a way for these viruses’ proteins to adapt to low pH environments. In support of this notion, repetitions of the GH18 domain are commonly found in eukaryotic cells, such as Ecdysozoa and fungi ^39^. Also, these protein architectures can vary in different organisms and have more than five domains. These repetitive structures can be explained by exon rearrangement events, and their existence may be related to increased efficiency of synergistic action in chitin degradation ^39–41^.

### BIOCHEMICAL CHARACTERIZATION OF CHLOROVIRUS GH18 CHITINASES DETERMINATE THEIR ENZYMATIC PROFILE

To experimentally characterize the chitinases from chloroviruses, we choose three GH18 domains with lower identity: (i) PBCV-1 GH18 domain 1 (herein termed GH18 PBCV-1 d1), common to all chloroviruses genomes; (ii) the second domain of PBCV-1 (GH18 PBCV-1 d2), representing the domain only present in alphachloroviruses; and (iii) the second domain of FR483 (GH18 FR483 d2), only present in betachloroviruses. The genes were cloned into a synthetic plasmid and expressed in *E. coli* with a N-terminal His-tag and pSUMO tag that can be cleaved off using the SUMO protease (Figure S4). The three enzymes were confirmed to be catalytically active by their ability to cleave the synthetic fluorogenic substrate, 4-Methylumbelliferyl N-acetyl-β-D-glucosaminide (4-MUF-NAG) (Figure 4).

**Figure 4.**
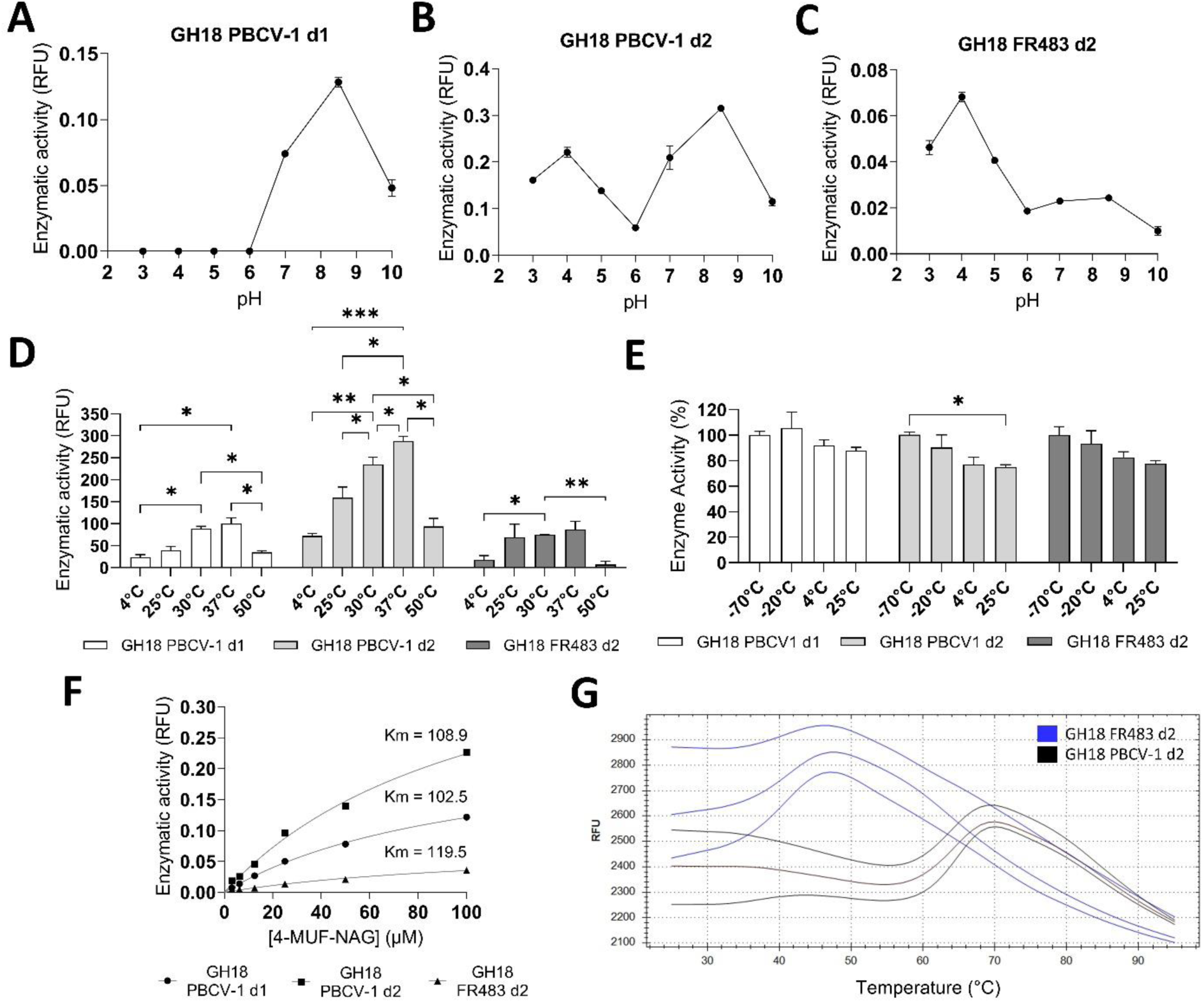
Enzymatic characterization of GH18 domains. **A, B, C:** Enzymatic activity PBCV-1 chitinase domain 1 and 2, and FR483 domain 2, respectively, in McIlvaine buffer at different pH. **D:** Enzymatic activity of the three GH18 domains at temperatures ranging from 4 to 50 °C. **E:** Stability of GH18 domains after 48h incubation at temperatures ranging from -70 to 25 °C. Two-Way ANOVA test was used to compare statistical differences at different temperatures. **F:** Michaelis-Menten curves and respective Michaelis constant (K_M_) obtained for the substrate 4-MUF-NAG. **G:** Temperature denaturation (T_m_) curves obtained from thermal shift assays of the GH18 PBCV-1 d2 (black) and FR483 d2 (blue) proteins. Each curve represents an independent replicate (*n* = 3).

Domain 1 of PBCV-1 GH18 could not cleave 4-MUF-NAG at pH 3 to 6 but had strong enzymatic activity at pH 7 to 10, with highest activity at pH 8.5. Domain 2 cleaved the substrate at all pH conditions tested, with highest activity with two peaks at pH 4 and 8.5. The second domain of FR483 cleaved the substrate mor efficiently at acidic pH with maximum activity at pH 4. (Figure 4A, B, C). The K_M_ for each enzyme was calculated to be 102.5 μM for PBCV-1 d1 (pH 8.5), 108.9 μM for PBCV-1 d2 (pH 8.5), and 119.5 μM for FR483 d2 (pH 3) (Figure 4F). GH18 family chitinases usually exhibit their highest activity at pH ranging from 3 to 7, and some organisms may have two domains acting as endochitinase (lower pH) and exochitinase (neutral to high pH) ^42^. Endochitinase cleaves the polymer at internal structural bonds, acting as N-acetyl-β-glucosaminidase, while exochitinase acts at terminal positions. This evolutionary adaptation confers an advantage to some microorganisms, as the same protein can digest different chitin polymer complexes in nature ^43–46^. GH18 PBCV-1 d1 was previously characterized as a unique enzyme, with maximum activity at pH ranging from 4 to 8 ^47^. Here, we show that each GH18 domain of PBCV-1 also has specific pH range for optimal activity. As previously demonstrated for CVK-2 virus, the PBCV-1 first domain likely acts as an exochitinase, and the second as an endochitinase, which is consistent with our characterization results (Figure 4B, C) ^48^.

The three domains are shown to be mesophilic enzymes, similar to those in soil and plant-associated bacteria, such as *Serratia marcescens* and *Bacillus thuringiensis* ^49,50^. The activity was evaluated between 4 and 50°C, with 37°C considered the optimal temperature for the three recombinant GH18 domains (Figure 4D). The activity of the three recombinant enzymes remained at nearly 90% after 48h incubation at 4 °C and 25 °C, while they maintained ∼100% activity after being incubated at negative temperatures (−70 and -20 °C), showing no or minimal statistical differences at different temperatures and overall high stability (Figure 4E). Finally, GH18 PBCV-1 d2 showed greater thermal stability compared to GH18 FR483 d2, with *T_m_* values of 64.3 and 41.0 °C, respectively (Figure 4G), having aggregation and dissociation states of the fluorophore beginning at 70 °C for PBCV-1 d2 and 45 °C for FR483 d2.

### STRUCTURAL COEVOLUTION AND PREDICTED LIGAND FITNESS IN GH18 CHITINASES SUPPORT THEIR ENZYMATIC PROFILE

To better understand the evolutionary divergence of chlorovirus chitinases, we first performed a coevolution analysis using 4,045 domain sequences from UniProt. In this analysis, we included experimental data obtained with the GH18 FR483 d2 that we purified, crystallized, and submitted to X-ray diffraction for data collection, resulting in a protein structure of 1.04 Å resolution (Table S2). Chloroviruses chitinases, including FR483 d2, lack several conserved residues with strong coevolutionary history in other GH18 family enzymes, such as human Chitotriosidase-1 (Figure 5A). Four distinct sets of coevolving residues within the GH18 family were identified (Figure 5B and Tables S3-S7). The first set consists of 10 residues surrounding the binding site, which are present in 54% to 70% of the sequences in our curated alignment (data not shown). Within chloroviruses sequences, two glycine and one aspartate were found in this region (Figure 5D, orange). The second set comprises six highly prevalent residues located primarily inside the TIM barrel, appearing in 79% to 93% of the aligned sequences (data not shown). Curiously, in chloroviruses, this group maintains a conserved motif consisting of the two catalytic aspartates, the catalytic glutamate, and a methionine (Figure 5C and D, purple). Conversely, the third and fourth sets highlight absent features in viral proteins: the third set represents coevolving cysteines involved in disulfide bridges (Figure 5B, yellow), while the fourth indicates coevolution between a glycine and a tryptophan near the active site (Figure 5B, cyan).

**Figure 5.**
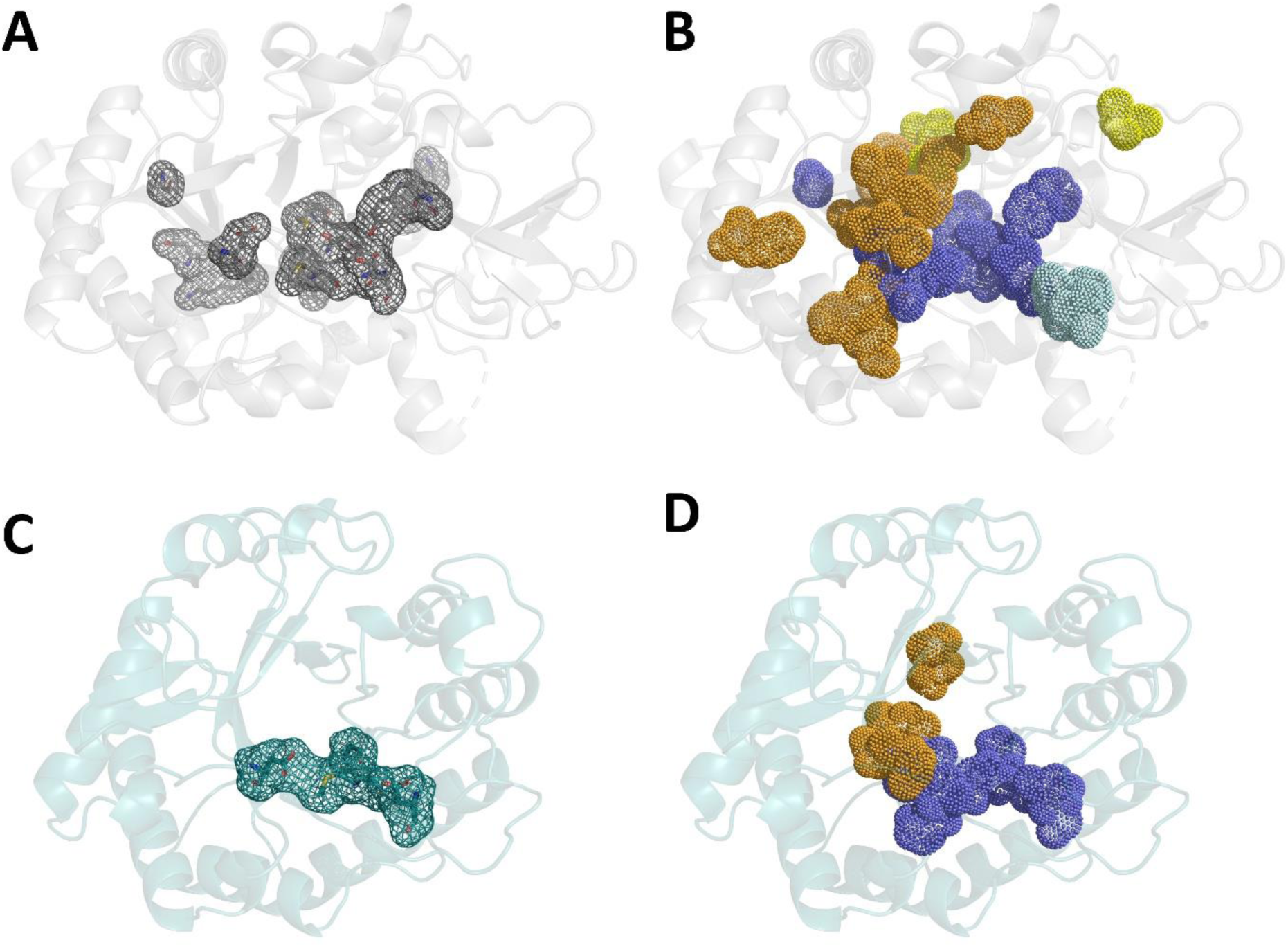
Residue conservation and coevolution in chitinases domains. **A:** Conserved residues (>80%) in chitinase domains, shown on the structure of human chitotriosidase (silver). **B:** Coevolving residue sets in chitinase domains. Set 1 (Y27, T52, H53, F58, G97, G98, F119, D136, W139, P142, D363, PDB 1GUV numbering) is shown in orange, set 2 (G134, D138, E140, M210, Y212, D213, G263, Y267, M358) in purple, set 3 (C26, C51, C307) in yellow, and set 4 (G216, W218) in cyan. **C and D:** conserved and coevolving residues (set 1 and 2) present in FR483 d2 domain (orange), which was crystallized with 1.04 Å resolution. The complete dataset is included in tables S3 to S7.

To further understand the impact of such conserved and non-conserved residues in the binding site of the three domains, we performed docking analysis with chitin tetramer and 4-MUF-NAG ligands. (Figure S5) Our predictions using GOLD v2025.1.0 resulted in an average CHEMPLP fitness scores (*n* = 200 runs) of 42.4 for PBCV-1 d1, 48.21 for PBCV-1 d2, and 42.76 for FR483 d2. Although small variations in average scores, our visual inspection of the docking poses revealed longer distances between the catalytic glutamate in PBCV-1 d1 and the chitin tetramer cleavage scissile bond (5.0 Å) (Figure S5E), when compared to PBCV-1 d2 (4.1 Å) (Figure S5F) and FR483 d2 (3.7 Å) (Figure S5G). Both poses predicted five hydrogen bonds for GH18 PBCV-1 d1 and four for both GH18 d2. Similar results were found for 4-MUF-NAG ligand, where both GH18 d2 showed a higher propensity for interaction, due to the shorter distances found between the catalytic residue and the fluorogenic substrate (Figure S5A-C). These results are consistent with our biochemical data (Figure 4F), in which GH18 d2 had slightly higher K*_m_* values. This shows that the presence of the second GH18 domains may be related to an increased affinity of these proteins for chitin polymers, impacting in the efficiency of its activity. Therefore, the second domain in chitinase 1 of chloroviruses suggests an important biological evolutionary adaptation of these viruses.

### CHARACTERIZED CHLOROVIRUS GH18 CHITINASES EXHIBIT SELECTIVE ANTIMICROBIAL ACTIVITY

Considering the biotechnology potential of chitinases in microbial control, we evaluated the biological activity of the GH18 domain 2 enzymes against fungi, algae, and cyanobacteria. First, the GH18 domains were tested against five yeasts and three filamentous fungi of medical clinical and agricultural importance, but no antifungal activity was observed up to 1024 µg/mL (Table S8). Next, and most importantly, the GH18 domain 2 of PBCV-1 and FR483 showed potent activity against the host algae of different chloroviruses, with inhibitory activities in the nanogram/mL range, except for *C. heliozoae* (Figure 6 and Figure S6). The two GH18 domains showed similar IC_50_ values for *Chlorella variabilis* NC64A and *Micractinium conductrix* Pbi, equal to 3 and 2 ng/mL for PBCV1 d2 and 63 and 59 ng/mL for FR483 d2, respectively. In *Chlorella variabilis* Syngen 2-3, the inhibitory activity was ∼10 to 15-fold lower compared to the first two species tested, having IC_50_ values of 32 ng/mL and 920 ng/mL for PBCV1 d2 and FR483 d2, respectively. Although some inhibitory activity was observed in *Chlorella heliozoae* SAG 3.83, the IC_50_ values could not be determined at tested concentrations, highlighting a lower susceptibility compared to the other algal species.

**Figure 6.**
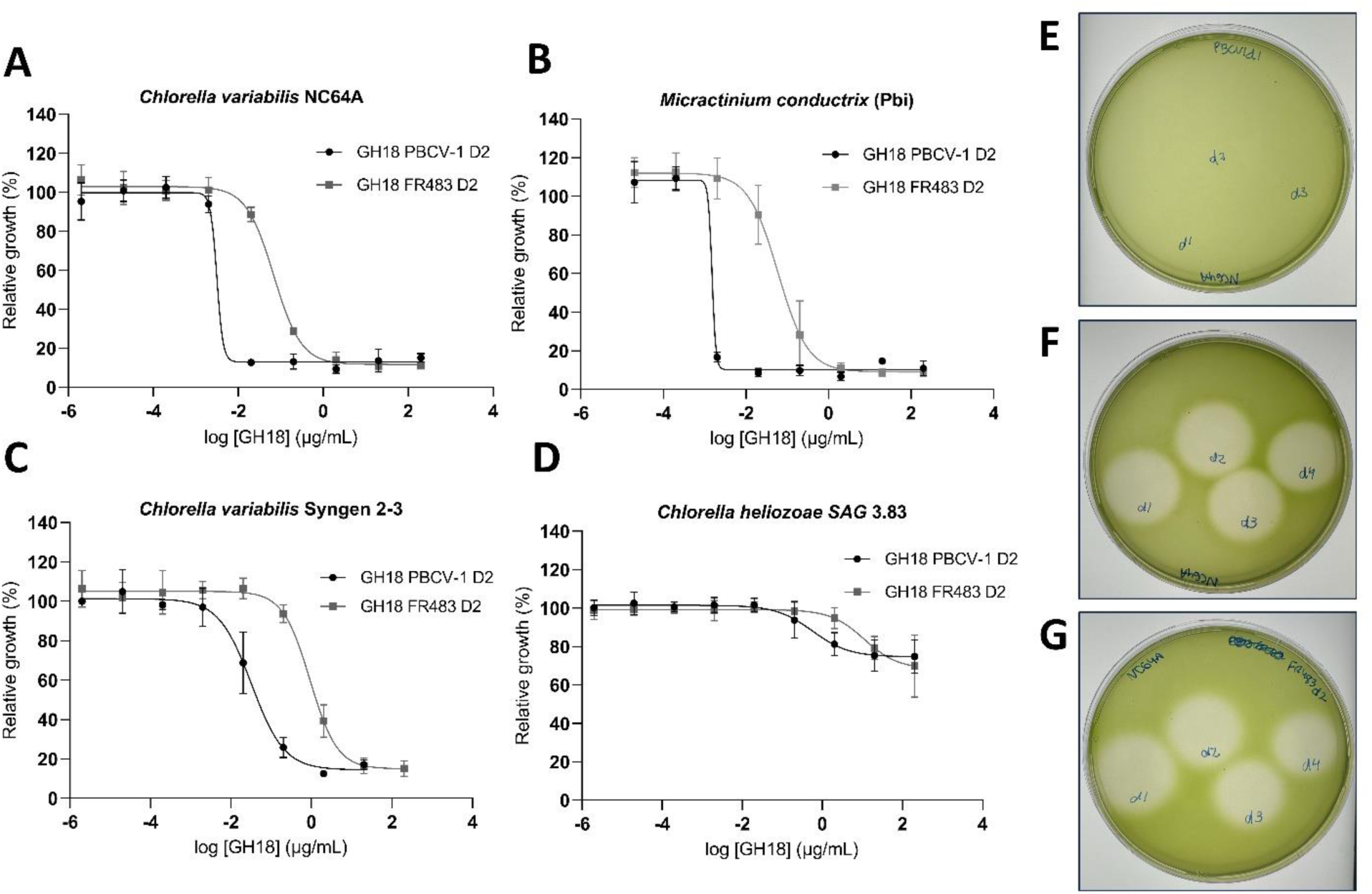
Antialgal activity of chlorovirus GH18 chitinases domains. IC_50_ curves for PBCV-1 d2 and FR483 d2 in the algal hosts of chloroviruses (*zoochlorella*). **A:** IC_50_ curves in *Chlorella variabilis* NC64A. **B:** IC_50_ curves in *Chlorella variabilis* Syngen 2-3. **C:** IC_50_ curves in *Micractinium conductrix*. **D:** IC_50_ curves in *Chlorella heliozoae* SAG 3.83. Zone of inhibition caused by chitinase inhibitory activity in an agar diffusion plate *vs.* **E:** *C. variabilis*, **F:** *M. conductrix*, and **G:** *C. heliozoae*.

Both enzymes were not active against *Chlorella vulgaris* and *Chlorella minutissima*, free-living species that are not hosts of chloroviruses, as well as against cyanobacteria (Table 1). Additionally, GH18 PBCV-1 d2 showed ∼30-fold increase in activity compared to GH18 FR483 d2 *vs. C. variabilis* NC64A and *M. conductrix* Pbi, and ∼28-fold *vs. C. variabilis* Syngen 2-3. These data suggest that both enzymes are highly specific to algae that are susceptible to chloroviruses infection, and their activity is even higher in the algae host of viruses that have the tested domains (*C. variabilis* NC64A and *M. conductrix* Pbi).

**Table 1.**
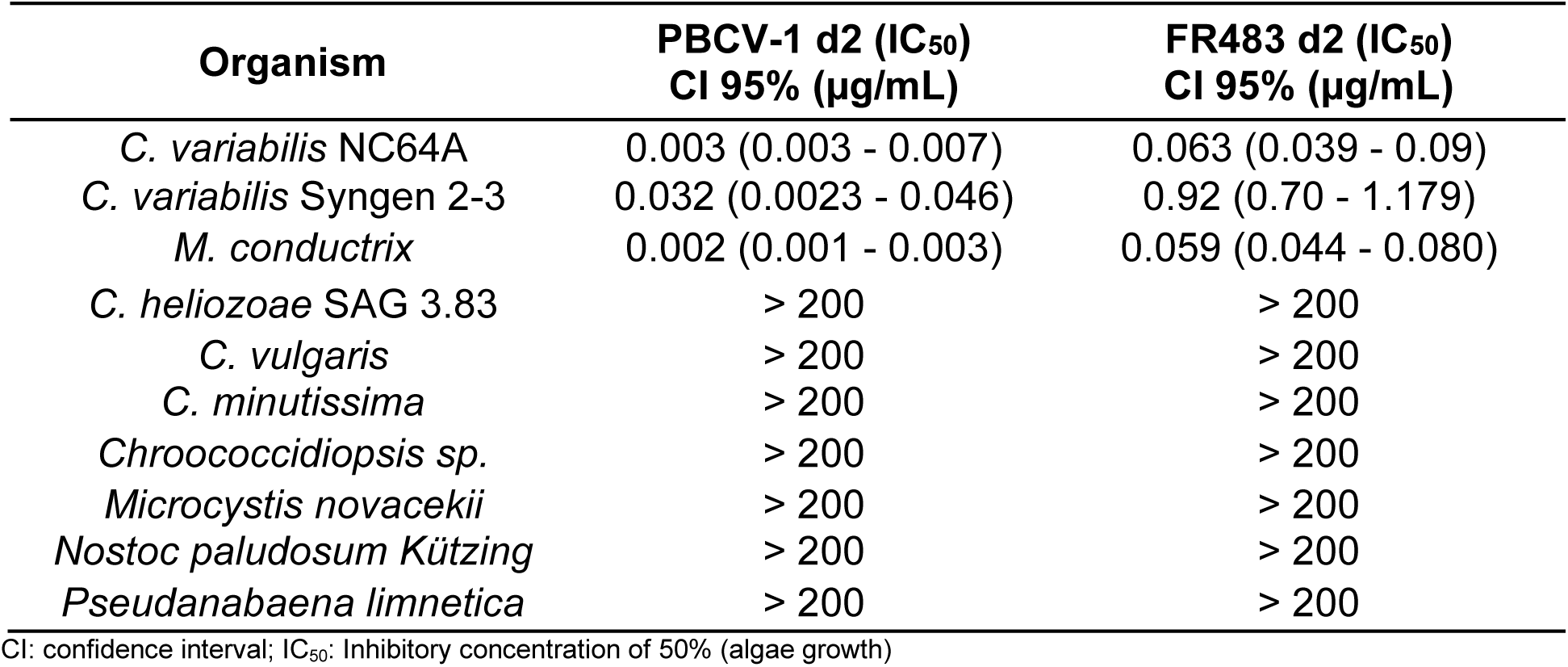
Anti-algal activity of chitinases GH18 domain 2 of PBCV-1 and FR483 against zoochlorella, free living chlorella and cyanobacteria, represented as IC50:

The lack of antifungal activity of viral chitinases, as well as inhibition in free-living algae and cyanobacteria, while having potent activity in endosymbiotic algae, evidence evolutionary mechanisms that favored the specificity of these proteins related to the host algae of chloroviruses. This may be associated with different cell wall composition, as previously described for fungi ^51^. The cell wall of *Chlorella* sp. is mainly composed of sugars, lipids, proteins, and a variety of carbohydrate polymers called glycosaminoglycans. The proportion and definition of the types of polymers present in each species are not yet fully understood ^52,53^. *Chlorella vulgaris*, the most described alga so far, presents other sugars as major components and cellulose as the most abundant carbohydrate, followed by chitin ^52,54^. Furthermore, studies have demonstrated that cellulases degrade of the cell wall in different species ^52^. On the other hand, in symbiotic algae, such as *Chlorella variabilis*, cellulose does not appear to be the major carbohydrate, but rather chitin^55^. In *Micractinium conductrix*, there are no documented studies, and our results suggest that the composition of its cell wall may be similar to *Chlorella variabilis*. Additionally, differences in glycosaminoglycans in the cell walls of free-living algae and endosymbionts have also been previously documented as a possible determining factor for the endosymbiotic relationship with the protozoan *Paramecium bursaria*, showing that the difference in cell wall composition affects the biology of algae between these two groups ^56^. Furthermore, the cell wall composition of *Chlorella heliozoae* appears to be different from *C. variabilis* and *M. conductrix*, as it is less susceptible to chitinase activity. This alga lives in endosymbiosis with another protozoan, *Acanthocystis turfacea*, and therefore, biological and structural factors that determine this relationship may be different when compared to algae associated with *Paramecium bursaria*.

Alternative methods are being studied for algae control, including biotechnological approaches. Chlorella organisms are not inherently toxic or harmful substances; however, they are associated with algal bloom events, in which physical and chemical changes foster their excessive growth, triggering environmental imbalances and resulting in biological and economic damage ^57,58^. Molecules used in water remediation, such as Fe_2_O_3_–TiO_2_ nanoparticles, have demonstrated activity against *Chlorella vulgaris*, by releasing reactive oxygen species ^59^. Surfactants have been considered promising compounds with low environmental impact, such as the gemini surfactant 12-5-12, which showed inhibitory activity of 19.6 µmol/L *vs*. *C. vulgaris*^60^. Synthetic compounds such as N,N’-diarylalkanediamides have also demonstrated activity against *C. vulgaris*, by inhibiting chlorophyll biosynthesis ^61^. Plant extracts also comprise a diverse source of potential compounds for antialgal control, such as those from *Salvia miltiorrhiza, Acorus tatarinowii,* and *Polygonum cuspidatum*, which demonstrated effective concentration of 50% (EC_50_) values between 98 and 130 mg/mL against *Microcystis aeruginosa* ^62^. Allelochemicals are also promising, such as compounds isolated from the green alga *Ulva pertusa*, which have demonstrated activity against red tide microalgae, with EC_50_ values up to 120 µg/mL ^63^. Overall, our results suggest chloroviruses chitinases as being highly specific to host algae. Future modifications, such as the addition of different CBM domains to the glycosyl hydrolase domain, may improve its specificity. Furthermore, addition of other GH18 domains from different organisms, as a hybrid protein, could also enhance broader antialgal activity to other different algal species ^64^.

## CONCLUSIONS

Chloroviruses are rich in genes involved in carbohydrate metabolism, and the presence and absence of these genes in different subgenera appear to be related to virus-host interaction. These genes may have been acquired through horizontal genome transfer from their algae hosts, undergoing differentiation and functional adaptation during evolution, becoming highly specific. Differences in optimal pH activities, thermal stability, and functional stability highlight the adaptation of these proteins and importance of the diversity of these genes in these viruses. They confer distinct structural domains, differences in amino acid residues present in binding sites and near the catalytic site, which may be responsible for providing different affinities with chitin, and adaptations to different cellular compartments. The GH18 FR483 d2 is the first non-structural protein domain coded by chloroviruses that was solved at high resolution (1.04 Å), and we showed that despite the lack of many of the coevolving residues compared to other organisms, this viral chitinase maintained its activity. We also indicated that evolutionary events are related to the variability and specificity of these bioactive enzymes, which have a promising application for algal control. We have determined potent and remarkable activity against some *Chlorella* strains, with IC_50_ values at the nanogram range. Furthermore, their thermostability and specificity are important characteristics for determining their applicability as a biotechnological alternative, deepening the understanding of these enzymes and emerging as a promising alternative in the control of algal blooms.

## METHODS

The Supporting Information (SI) includes all the information about the methods used in this study. No unexpected or unusually high safety hazards were encountered. All genome sequences used in this study are publicly available in GenBank. The chlorovirus GH18 protein structure obtained by X-ray crystallography is available at the Protein Data Bank (PDB ID 37AC).

### Supporting Information

Experimental methods and materials; Figure S1. Identity of chloroviruses’ chitinase sequences and their distribution among subgenera; Figure S2. Phylogenetic reconstruction of chitinase 1, including viral enzymes and cellular counterparts; Figure S3. Frequency of predicted amino acid residues in the binding site of domain 2 GH18 of alphachloroviruses and betachloroviruses; Figure S4. Cloning plasmid and enzymatic characterization of GH18 domains; Figure S5. Interaction of GH18 domains with 4-MUF-NAG and chitin tetramer; Figure S6. Antialgal activity against Chlorella using agar diffusion test. Table S1. Distribution of CAZymes found in 89 chloroviruses genomes; Table S2. X-ray diffraction data collection, data processing, and crystal structure refinement statistics for the catalytic domain 2 of FR483 betachlorovirus; Table S3. Prevalence of coevolving residues on chlorovirus chitinases in set 1. Colored in green represent coevolved residues; Table S4. Prevalence of coevolving residues on chlorovirus chitinases in set 2. Colored in green represents coevolved residues; Table S5. Prevalence of coevolving residues on chlorovirus chitinases in set 3; Table S6. Prevalence of coevolving residues on chlorovirus chitinases in set 4; Table S7: Prevalence of conserved residues on chlorovirus chitinases in set 4. Colored in green represents coevolved residues; Table S8. Minimum inhibitory concentration (MIC) of GH18 domains in yeast-like, dimorphic, and filamentous fungi at different pH ranges.

## Supporting information

Supporting information with methods, figures and tables

## ACKNOWLEDGEMENTS

We thank all colleagues and staff from Laboratório de Vírus of UFMG and Centro Nacional de Pesquisa em Energia e Materiais (CNPEM) for all the technical support during the development of this work. We thank Professors James Van Etten (UNL-USA) and Francisco Barbosa (UFMG-Brazil) for providing algal and cyanobacteria cultures used in this work for biological assays. We also thank Liza Felicori (UFMG-Brazil) for the useful discussions during the development of this work and for the critical reading of this manuscript. This work was partially funded by the Fundação de Amparo à Pesquisa do Estado de Minas Gerais (FAPEMIG), Grant n° APQ-01057-23 & BDT-00022-25, and Coordenação de Aperfeiçoamento de Pessoal de Nível Superior (CAPES), financial code 001. E.G.O acknowledges previous CAPES-PrInt scholarship (88887.911426/2023−00). M.S.M.S. acknowledges his FAPEMIG fellowship (APD-01212-25) and his previous PDE scholarship by CNPq (200069/2024-1). L.B. and R.A.L.R. are CNPq researchers.

