## Supporting information with methods, figures and tables for "Carbohydrate active enzymes of giant viruses: Molecular and biochemical characterization of glycosyl hydrolases from algae-infecting chloroviruses"

#### EXPERIMENTAL METHODS

##### Annotation of genes involved in sugar and carbohydrate metabolism.

Open reading frame regions (ORFs) of 89 chlorovirus genomes (table S1) were predicted *de novo* using the online software GeneMarkS [1], with the "Prokaryotes" option. The predicted ORFs were annotated for identification of corresponding coding sequences (CDSs) using Blastp (*e* value $< 10^{-5}$) against the NCBI non-redundant (nr) proteins database, followed by protein domain search using HHpred [2]. In case of disagreement in the results, the Interproscan software was used to obtain the final annotation [3–6]. The number of genes involved in sugar and carbohydrate metabolism was manually annotated among the subgenera *Alphachlorovirus* (NC64A and OSy-viruses), *Betachlorovirus* (Pbi-viruses), and *Gammachlorovirus* (SAG-viruses), based on genomic annotation obtained from the 89 genomes. One-way ANOVA statistical analysis was performed in GraphPad Prism 9.0 for comparative purposes.

##### Identification, comparison, and *in silico* characterization of chitinase sequences in chlorovirus genomes.

To assess the similarity between the amino acid sequences of chitinase 1 and chitinase 2 among the viruses used in this study, a comparative analysis was performed employing 128 chlorovirus genomes. The sequences were obtained from GenBank, separated, and used in an internal script based on global pair alignment. For comparison purposes, the amino acid sequences of the hallmark gene, topoisomerase II, was used. The identity values were employed for comparison purposes, which required the reclassification of the pairs previously designated as Species X *vs.* Species Y from the *Chlorovirus* subgenus that each virus belongs [7]. Statistical analyses and figure construction were performed using R software version 4.4.3 "Trophy Case" (R Core Team, 2025), with the car (Fox and Weiseber, 2019) and tidyverse (Wickham et al., 2019) packages, respectively. More specifically, regarding statistics, the normality and homoscedasticity of the samples were evaluated using the Kolmogorov-Smirnov and Levene tests. Given the non-normal and variable nature of all samples, the Kruskal-Wallis test was employed for a comparative analysis of identities, followed by pairwise comparisons using the Wilcoxon rank-sum test to identify which groups were divergent. For both tests, $p$-values < 0.05 were considered to identify statistical differences between groups.

For comparative structural analysis 128 chitinase 1 sequences (DxDxE as the catalytic site) and 95 chitinase 2 sequences (DxE as the catalytic site) were identified. All sequences were submitted to the PROSITE [8] Expasy platform (https://prosite.expasy.org/) for prediction of domains, motifs, and catalytic sites. Domains not found by the platform were check based on HHPred for data complementarity [2]. Multiple sequence alignment using the MEGA 11 software [9] of all chitinase 1 sequences was performed to obtain the percentage of identity and identification of amino acid residues conservation between the sequences. Chitinase sequences were submitted to AlphaFold2 [10] via ColabFold [11] (v1.5.5; https://github.com/sokrypton/ColabFold) using default parameters (no template mode and no relaxed structures). Each model was considered based on a predicted Template Modeling score (pTM) > 0.80[10]. Modeled structures were submitted to PrankWeb [12] 3 (https://prankweb.cz/) to perform their binding site characterization using the AlphaFold strucure mode with default parameters (using conservation). Predictions were performed considering binding sites with a probability > 0.5 [13]. Comparative characterization between binding sites was performed using sequence alignments previously obtained by MEGA [9], and comparing the predominance of residues present in each binding site among isolates of all three subgenera.

**Phylogenetic analysis of chitinase gene**

Sequences of chitinase 1 from 89 chloroviruses and different organisms were obtained from the NCBI database. Sequence alignment was performed using MUSCLE available in MEGA 11 with default parameters [9,14]. A maximum likelihood reconstruction was performed using IQ-TREE v2.3.5 with 1,000 bootstrap replicates [15]. The best substitution model, VT+F+R8, was obtained with ModelFinder [16]. The resulting phylogenetic tree was visualized and edited using iTOL v6 [17].

**Docking analysis of ligand interactions to modeled chitinase structures.**

The modeled structures were aligned and analyzed with PyMOL software (v2.5.7) (PyMOL Molecular Graphics System, DeLano Scientific LLC, 2006). The 3D structures of the molecules selected as ligands (4-MUF-NAG and chitin tetramer) were built using Discovery Studio (BIOVIA, USA, 2017), as described[18]. Docking analysis were performed using GOLD 5.1 (v2025.1.0)[19], as described[18]. Modeled structures were prepared by adding hydrogen atoms only. The options and parameters in GOLD were as follows: (i) all rotating protein bonds were fixed; (ii) binding site defined with the centered residue (catalytic glutamate) and all atoms within 20 Å; (iii) "chemscore_kinase" as a template; (iv) 200 genetic algorithm (GA) runs; (v) CHEMPLP as a scoring function; (vi) no early termination allowed; (vii) and GA search options set to slow. The overall average docking scores (CHEMPLP fitness) were used for comparison purposes[20]. Poses were selected based on scores and by visual inspection[21]. The PyMOL software (v2.5.7) was used to analyze results and generate images.

**Cloning of GH18 domains into the synthetic pSUMO plasmid.**

Three predicted GH18-domain sequences of chitinase 1, selected as the three sharing the lowest pairwise sequence identity, were cloned into a synthetic pSUMO plasmid (5,763 bp) encoding the Small Ubiquitin-like Modifier (SUMO) protein. On this basis, GH18 domains 1 and 2 of PBCV-1 and domain 2 of virus FR483 were cloned into pSUMO. *Escherichia coli* BL21 was transformed by heat shock. For protein expression, BL21 clones were grown in LB medium containing 50 µg/mL ampicillin with shaking at 150 rpm. When the $OD_{600}$ reached 0.4–0.8, expression was induced with IPTG at a final concentration of 0.5 mM, and the cultures were incubated overnight at 20 °C and 150 rpm. Cells were harvested by centrifugation at 3,000 × *g* for 15 min, the supernatant was discarded and the pellet was resuspended in lysis buffer (20 mM Tris HCl, 300 mM NaCl, 10% glycerol, 1 mM DTT). Cells were lysed using a high-pressure microfluidizer, and the lysate was clarified by centrifugation at 20,000 × *g* for 20 min at 4 °C. The supernatant was applied in lysis buffer to an ÄKTA pure™ system fitted with a nickel (Ni-NTA) column for initial purification of the GH18–SUMO fusion. The fraction eluted with 500 mM imidazole was collected, then concentrated and buffer-exchanged (lysis buffer) using Amicon Ultra 10 kDa devices (Ultracel-10k; Millipore, Merck) until the imidazole concentration fell below 10 mM. The fusion protein was subsequently incubated overnight at 4 °C with His-tagged SUMO protease under gentle agitation. A second Ni-NTA step was then performed, in which the cleaved GH18 protein was recovered in the imidazole-free flow-through while the His-tagged SUMO moiety and the protease were retained on the column. This flow-through fraction, containing the purified GH18 protein, was collected, concentrated, and again buffer-exchanged (lysis buffer) with Amicon Ultra 10 kDa devices (Ultracel-10k; Millipore, Merck). Protein concentration was determined with the Pierce™ BCA Protein Assay kit, and the purified protein was stored at −70 °C. Selected fractions were further purified by size-exclusion chromatography with the same lysis buffer in preparation for quantification and subsequent crystallization trials.

**Characterization of the enzymatic activity of GH18 chitinase domains**

The substrate used to test the activity for each purified GH18 chitinases was 4-methylumbelliferyl N-acetyl-β-D-glucosaminide (4-MUF-NAG; Sigma-Aldrich), which releases the fluorophore 4-methylumbelliferone (excitation 360 nm, emission 460 nm) upon hydrolysis [22,23]. To determine the optimal pH range, a total of 6 ng of GH18 PBCV-1 d1, 80 ng of GH18 PBCV-1 d2, and 60 ng of FR483 d2 were incubated with 100 µM of the 4-MUF-NAG substrate. The substrate was diluted in McIlvaine (citrate phosphate) buffer at pHs 3.0, 4.0, 5.0, 6.0, 7.0, 8.5, and 10.0. Enzymatic activity was measured with a Synergy fluorescent reader (360/460 nm) for 2 h at 37 °C. The Vmax value was used to compare activities at different pHs. For temperature optimization, a total of 6 ng of GH18 PBCV-1 d1, 80 ng of GH18 PBCV-1 d2, and 60 ng of FR483 d2 were incubated with 100 µM of the 4-MUF-NAG substrate in McIlvaine buffer at pH 8.5 or pH 3.0. The tubes, in triplicate, were incubated at different temperatures (4, 25, 30, 37, 50, and 60 °C) for 2 h. The reaction was stopped with 3 M urea, and the Vmax value was obtained by a single reading. For temperature stability assay, a total of 20 µL of each purified enzyme were incubated at 4 °C, 25 °C, -70 °C, and -20 °C for 48 h in triplicate. After this period, a total of 6 ng of GH18 PBCV-1 d1, 80 ng of GH18 PBCV-1 d2, and 60 ng of FR483 d2 were incubated with 100 µM of the 4-MUF-NAG substrate, and the enzymatic activity was measured with the Synergy reader (360/460 nm) for 2 h at 37 °C and Vmax value was used to obtain the relative activities. For Michaelis-Menten curve, a total of 6 ng of GH18 PBCV-1 d1, 80 ng of GH18 PBCV-1 d2 and 60 ng of FR483 d2 were incubated with 200, 100, 50, 25 and 12.5 μM of the substrate 4-MUF-NAG, diluted first in DMSO and then in McIlvaine buffer at pH 8.5. The highest concentration was chosen to achieve a maximum of 1% DMSO at all substrate concentrations. Enzymatic activity was measured with the Synergy reader (360/460 nm) for 2 h at 37 °C. Product formation curves for each concentration of 4-MUF-NAG were obtained in triplicate, with their respective $V_0$ values. The Km value (substrate concentration at which velocity = ½ Vmax) was calculated from the $V_0$ values, and two-Way ANOVA statistical analysis was performed using Prism 9.0 software

**Thermal Shift Assay**

A total of 20 mg of the GH18 PBCV-1 d2 and GH18 FR483 d2 proteins were incubated with Invitrogen™ SYPRO™ dye diluted 1:1000 in Tris buffer. The buffer pH was adjusted to 8.0 for GH18 PBCV-1 d2 and 3.0 for GH18 FR483 d2, corresponding to the optimal pH for the activity of each enzyme. Thermal denaturation assays were performed using a Bio-Rad CFX96 real-time PCR system, monitoring fluorescence with excitation and emission wavelengths of 533 and 559 nm, respectively (HEX fluorophore). The temperature was gradually increased at 2-second intervals up to 90°C, and melting curves were generated from the resulting fluorescence data.

**Experimental Determination for Structural Characterization: Crystallization**

Crystallization trials were performed using an automated high-throughput screening approach at the Brazilian Biosciences National Laboratory (LNBio), part of the Brazilian Center for Research in Energy and Materials (CNPEM, Campinas, SP, Brazil). A total of 1,728 crystallization conditions were screened at 18°C for 30 days using the following commercial crystallization kits: Crystal Screen and Crystal Screen 2 (288 conditions; Hampton Research), Wizard I and Wizard II (288 conditions; Emerald BioSystems), PACT Suite (288 conditions; Qiagen), JCSG Suite (288 conditions; Qiagen), SaltRx (288 conditions; Hampton Research), and Precipitant Synergy (288 conditions; Emerald

BioSystems). The most promising condition was subsequently optimized using the hanging-drop vapor-diffusion method. Crystallization drops consisted of a mixture of protein and reservoir solution in a 1:2 ratio (2 µL protein solution and 4 µL reservoir solution). The optimized reservoir solution consisted of 0.1 M imidazole buffer (pH 8.0), 20% (w/v) PEG 8000, and 0.2 M NaCl. Crystal growth was observed after approximately 30 days. Crystals were harvested using nylon loops (Hampton Research) and transferred to a cryoprotectant solution consisting of the reservoir solution supplemented with 20% (v/v) ethylene glycol (2 µL ethylene glycol added to 8 µL reservoir solution). The crystals were then flash-cooled in a nitrogen gas stream at −173°C prior to X-ray diffraction data collection.

**Experimental Determination: X-ray diffraction of protein crystals**

The GH18 FR483 d2 crystals were used for X-ray diffraction data collection and three-dimensional structure determination. Diffraction experiments were carried out at the Manacá beamline of the Brazilian Synchrotron Light Laboratory (LNLS), part of the Sirius synchrotron facility (Campinas, SP, Brazil). Diffraction images were recorded using a CCD detector. The collected data were indexed, integrated, and scaled using the XDS software package. The structure was solved by molecular replacement as implemented in the PHENIX suite [24]. using an AlphaFold-predicted model as the search model. Initial model building was performed automatically with AutoBuild and subsequently improved through manual model adjustment in Coot [24,25]. Iterative rounds of refinement were carried out using *phenix.refine* [24] until convergence of the crystallographic statistics and optimization of the structural model.

**Evolutionary structural analysis of chitinases 1**

A total of 4045 chitinase sequences were scanned for Interpro [1] domains, and on most of them at least one of three domains were present: CBM2 Carbohydrate Binding Module 2, represented by Pfam entry PF00553, Glycosyl Hydrolase 18 Chitinase, represented by Panther entry PTHR42976, and Glycosyl Hydrolases family 18 (GH18), represented by Pfam entry PF00704. The latter corresponds to the crystallized segment in the FR483 d2 protein. Sequences having these domains were obtained from representative proteomes using a 15% cutoff in Uniprot [27]. These sequences were aligned with the viral sequences used in this study, using HMMER. This alignment was used for conservation and coevolution analysis using PFstats [28]. The alignment was filtered to remove possible fragments and redundancy. Residues were considered conserved if present in at least 80% of the final curated alignment. Coevolving residues were indicated if the presence of a specific residue in a given position increased the frequency of another residue in a different position to have a value higher than 80%, while the associated *p*-value is less than $10^{-10}$, and the two residues existed in at least 10% of the sequences, as determined by a Dima-Thirumalai test. The PyMOL software (v2.5.7) was used to analyze results and generate images.

**Antifungal Activity Assay**

Five yeast species that cause diseases in humans were used: *Candida albicans* (ATCC 18804), *Candida parapsilosis* (ATCC 22019), *Candida krusei* (ATCC 6258), *Candida auris* (CBS 10913), *Cryptococcus neoformans* (H99). In addition, three filamentous fungi were also considered: *Sporothrix brasiliensis* (ATCC 5110), *Aspergillus fumigatus* (ATCC 16913), and *Fusarium spp.* The inoculum for each species was prepared using the cell density method. Briefly, cultured colonies in XYZ agar were collected and adjusted to a McFarland standard solution of 0.5 at 530 nm, that is, by adding sufficient saline in a glass tube to obtain the equivalent transmittance at 530 nm, corresponding to $1 \times 10^6$ to $5 \times 10^6$ cells/mL (OD 0.09 to 0.13 for Sporothrix, 83% T for Aspergillus, OD 0.15 to 0.17 for Fusarium). The working solution added in 96-well microplates is produced by making a 1:50 dilution of each fungi suspension, followed by a 1:20 dilution for yeasts and a 1:50 dilution for filamentous fungi, using RPMI 1640 medium. Final concentrations were in between $5.0 \times 10^2$ to $2.5 \times 10^3$ cells/mL. The assay was performed in RPMI 1640 medium at pH 7-8 and at pH 3-4, considering the optimal activity range of the two enzymes. The minimum inhibitory concentrations (MIC) for fluconazole (FCZ) (Sigma-Aldrich), amphotericin B (AMB) (Sigma-Aldrich), and the chitinases PBCV1 d1 and FR483 d2 were determined by broth microdilution, following the M27-S4 method [29] of the Clinical and Laboratory Standards Institute (CLSI). The final concentrations ranged from 0.125 to 64.0 µg/mL for FCZ, 0.03 to 16.0 µg/mL for AMB, and up to 1024 μg/mL for the viral chitinases. The microplates were incubated at 37 °C for 72 h and a visual inspection was performed to assess antifungal activity, considering wells without turbidity, *i.e.* no visual growth, using the control wells as a reference control. One independent assay in triplicate was performed (*n* = 3 data points).

**Antialgal 50% Growth Inhibition ($IC_{50}$) Assay**

The test was performed assessing the chloroviruses' host species *Chlorella variabilis* NC64A, *Micractinium conductrix* Pbi, *Chlorella heliozoae* SAG 3.83, and *Chlorella variabilis* Syngen 2-3. These cells were kindly provided by Professor James Van Etten (Department of Plant Pathology, University of Nebraska-Lincoln, USA). The free-living algal species *Chlorella variabilis* and *Chlorella minutissima,* and the cyanobacteria *Chroococcidiopsis* sp, *Microcystis novacekii*, *Nostoc paludosum Kützing*, *Pseudanabaena limnetica* were also included for testing. These cells were kindly provided by Professor Francisco Barbosa (Department of Genetics, Ecology, and Evolution, UFMG, Brazil). The evaluation of the antialgal activity of chitinases was performed by determining the $IC_{50}$ values, which corresponds to the concentration capable of inhibiting 50% of cell growth. The experiment was performed in 96-well microplates with a total of $2x10^4$ cells/well in MBBM (Modified Bold's Basal Medium. Serial dilutions of the chitinases PBCV1 d2 and FR483 d2 were performed in MBBM medium in a 1:10 serial dilution to achieve a final concentration of 200, 20, and 2 μg/mL, and 200, 20, and 2 pg/mL. The microplates were incubated for 72 h under continuous light at 25 °C. The reading was performed in a spectrophotometer (VersaMax, San Jose, CA, USA) at 550 nm. Three independent assays in triplicate were performed. The $IC_{50}$ values were obtained from the growth relative percentage/log concentration curve using GraphPad Prism 9.0.

**Double-Layer Agar Diffusion Assay**

The double layer agar diffusion test was performed assessing chloroviruses' host species *Chlorella variabilis* (NC64A), *Micractinium conductrix* (Pbi), and *Chlorella heliozoae* (SAG 3.83). A total of $1x10^8$ cells were plated on MBBM agar or FES (Forssbohm's Emerson Synthetic medium) agar 0.75% (Pbi only) in Petri dishes above agar 1.5%. The three chitinases obtained were filtered using a 0.22 μm filter to remove contaminants. The concentrations used for PBCV-1 d1 were 7.3, 3.15, and 1.5 μg, due to the low yield obtained in its purification. As for the other two chitinases, concentrations were 75, 37.5, 18.7, and 9 μg. The drop diffusion method was used on 0.75% agar until completely dry, and the plates were incubated under continuous light at 25 °C for 72 h.

### SUPPLEMENTARY FIGURES AND TABLES

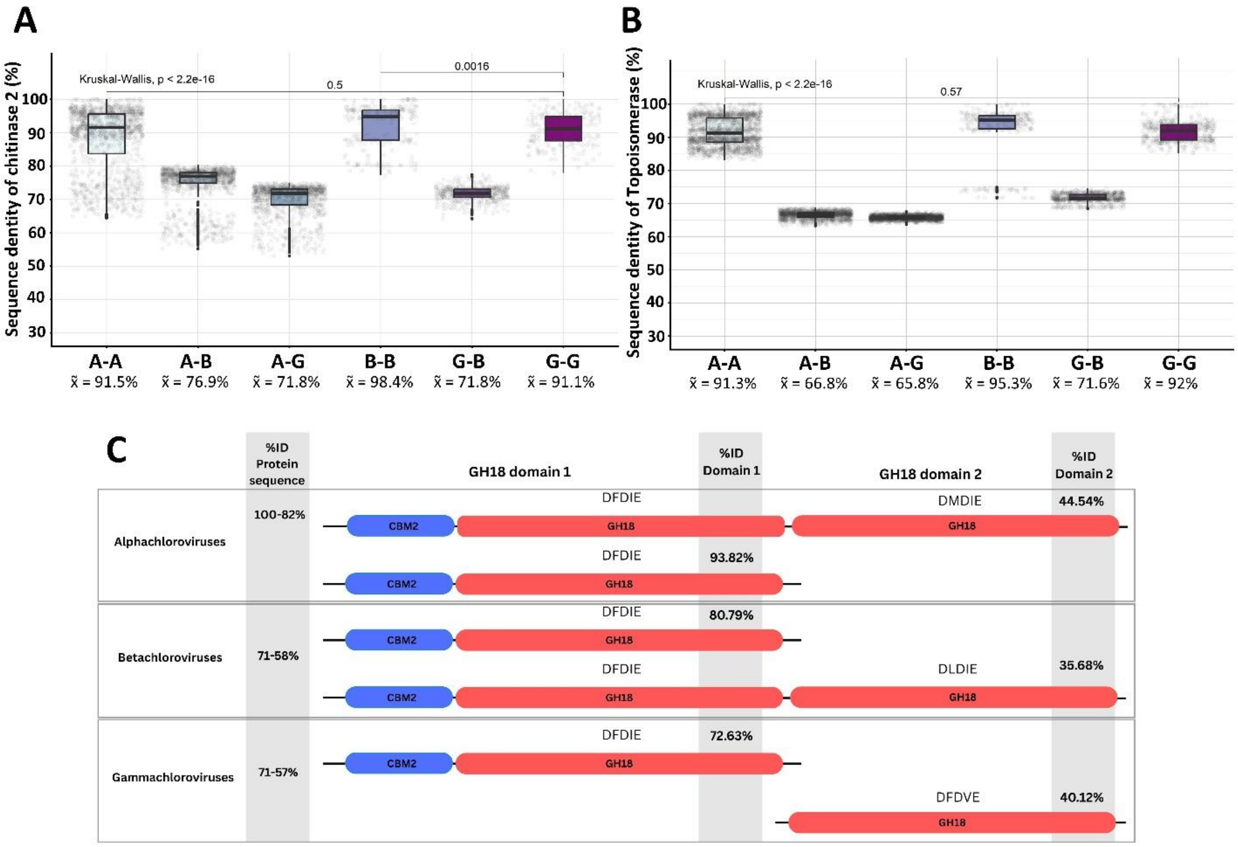

**Figure S1. Identity of chloroviruses' chitinase sequences and their distribution among subgenera.** Pairwise comparison of amino acid residues sequence identity of chitinase 2 **(A)** and topoisomerase **(B)** gene found in 128 genomes of chlorovirus isolates from all three subgenera (*Alphachlorovirus*, *Betachlorovirus*, and *Gammachlorovirus*). Each gray dot represents a percentage value of nucleotide identity obtained from the alignment of two different sequences. A-A: Alpha-Alpha, A-B: Alpha-Beta, A-G: Alpha-Gamma, B-B: Beta-Beta, G-G: Gamma-Gamma. Avg: mean. Med: median. Kruskal-Wallis non-normal distribution statistical analysis was performed, with the highest p-value $<2.2e^{-16}$ ($p$-values $<0.05$ indicate differences between the medians of the groups). The $p$-values shown in the bars were obtained from the Wilcoxon analysis ($p$-values $< 0.05$ and the absence of numbers indicate that there are differences between the medians of the groups). **C:** Schematic representation of domains present in the sequences of chitinases 1 predicted in 128 chloroviruses, with their identity values varying between viral subgenera. CBM2: Chitin binding module 2. GH18: glycosyl hydrolase 18. Letters above the GH18 domain representations indicate the motif of residues found in the catalytic site.

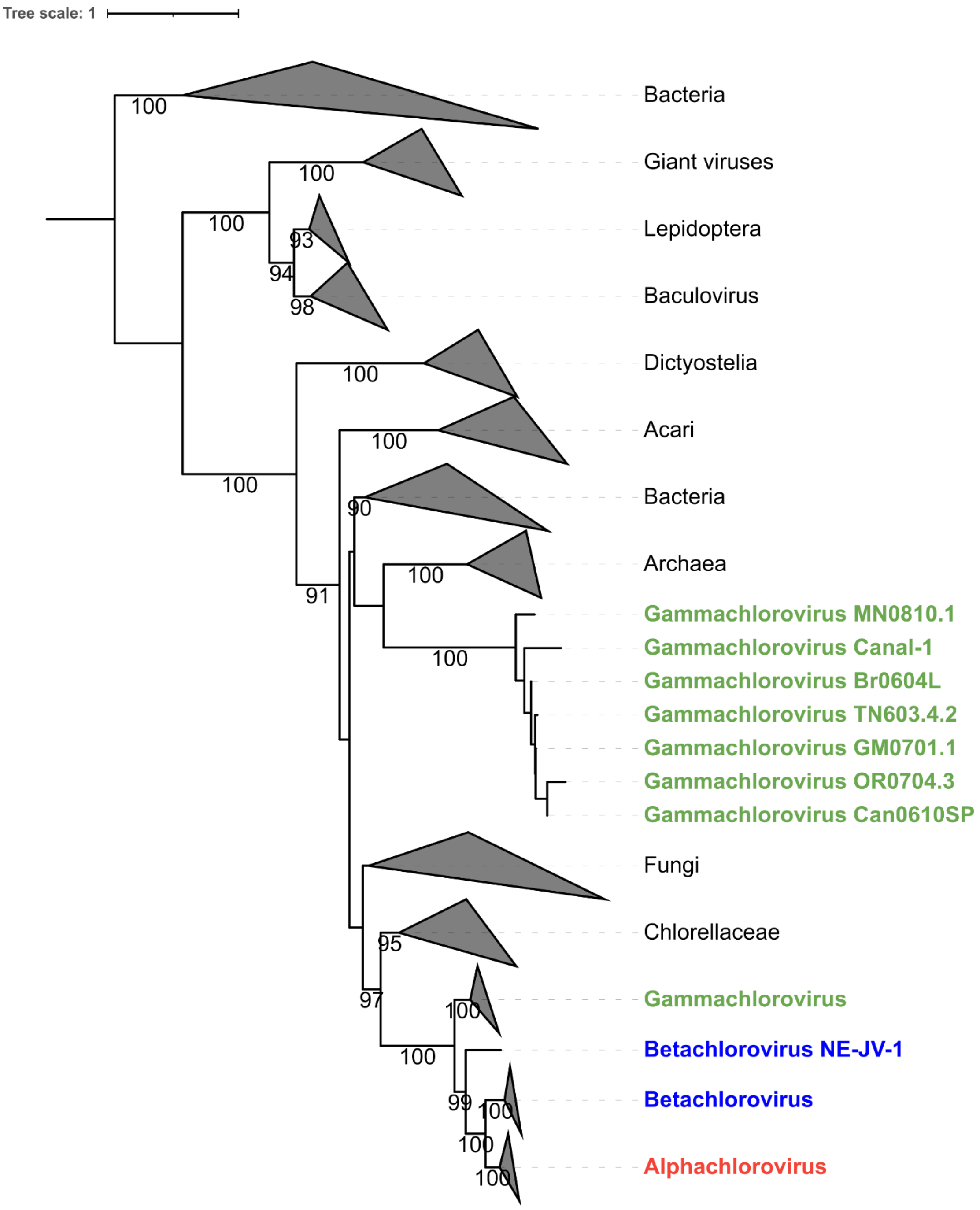

**Figure S2. Phylogenetic reconstruction of chitinase 1, including viral enzymes and cellular counterparts.** Maximum likelihood tree constructed with 1,000 bootstrap replicas based on residues sequence alignment of GH18 chitinases found in different viruses and cellular organisms. Phylogenetic tree was built using IQ-tree 2.2.0 and visualized using iToL. Scale bars represent the rate of residue substitutions.

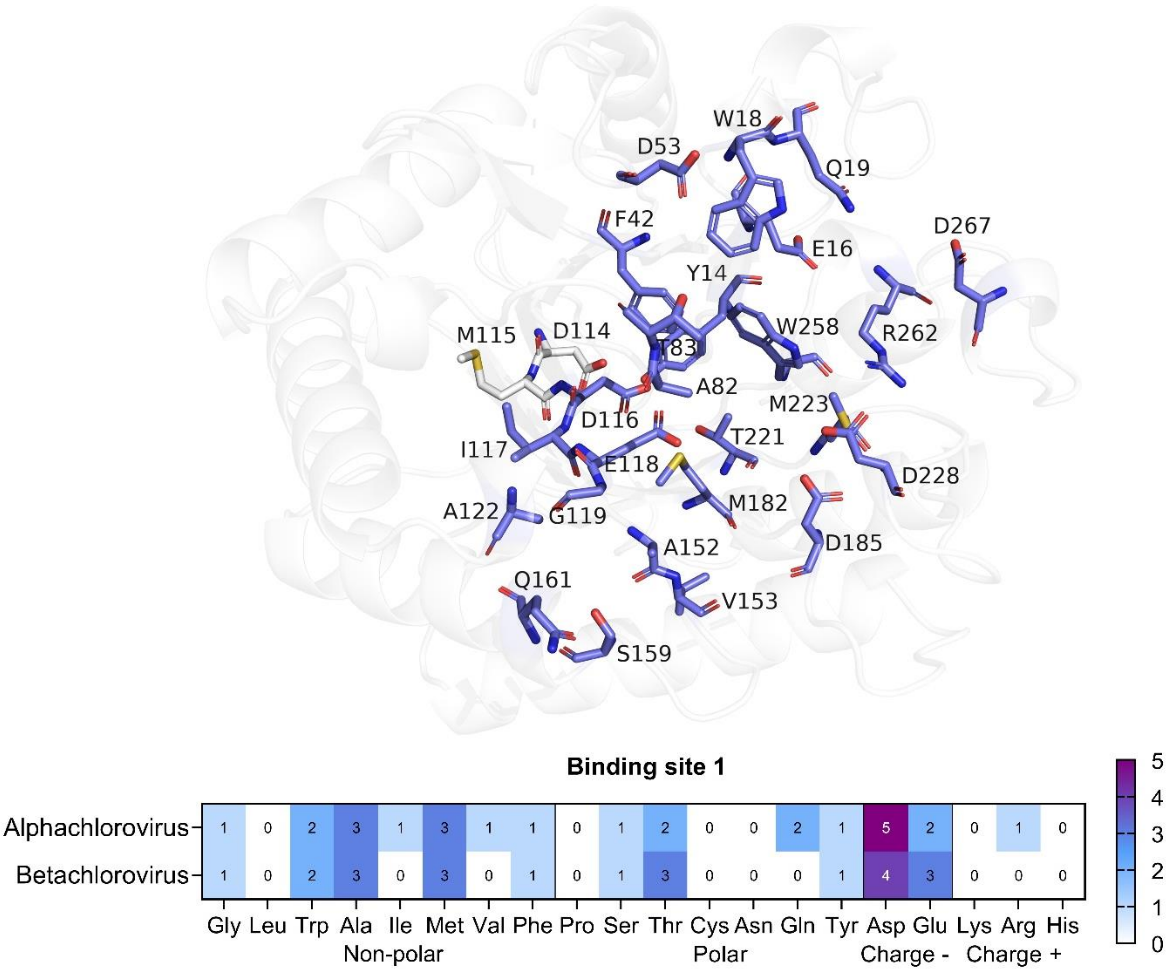

**Figure S3. Frequency of predicted amino acid residues in the binding site of domain 2 GH18 of alphachloroviruses and betachloroviruses.** The three-dimensional structure corresponds to the prediction of domain 2 of the PBCV-1 chlorovirus chitinase by AlphaFold2. The pocket predicted by PrankWeb is represented in pink. Stick amino acids correspond to those that make up the catalytic site motif (DMDIE). Absolute numbers of each amino acid are described in the heatmap.

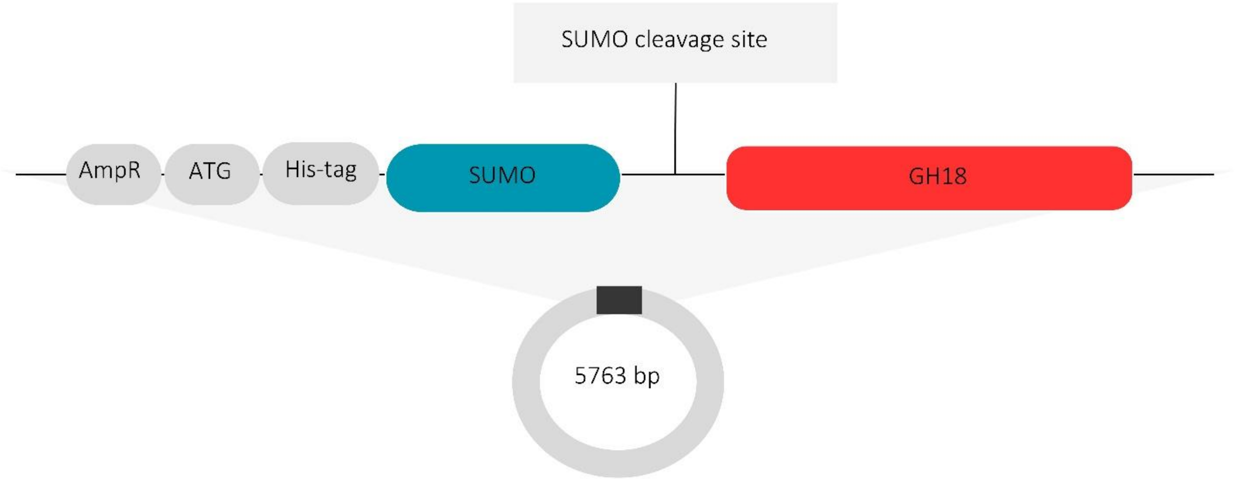

**Figure S4. Cloning plasmid and enzymatic characterization of GH18 domains. A:** Construction of the cloning synthetic pSUMO plasmid (5763 base pairs). With ampicillin resistance gene, 6-histidine tail linked to the pSUMO protease, the coding sequence, and the pSUMO protein cleavage site (two glycines).

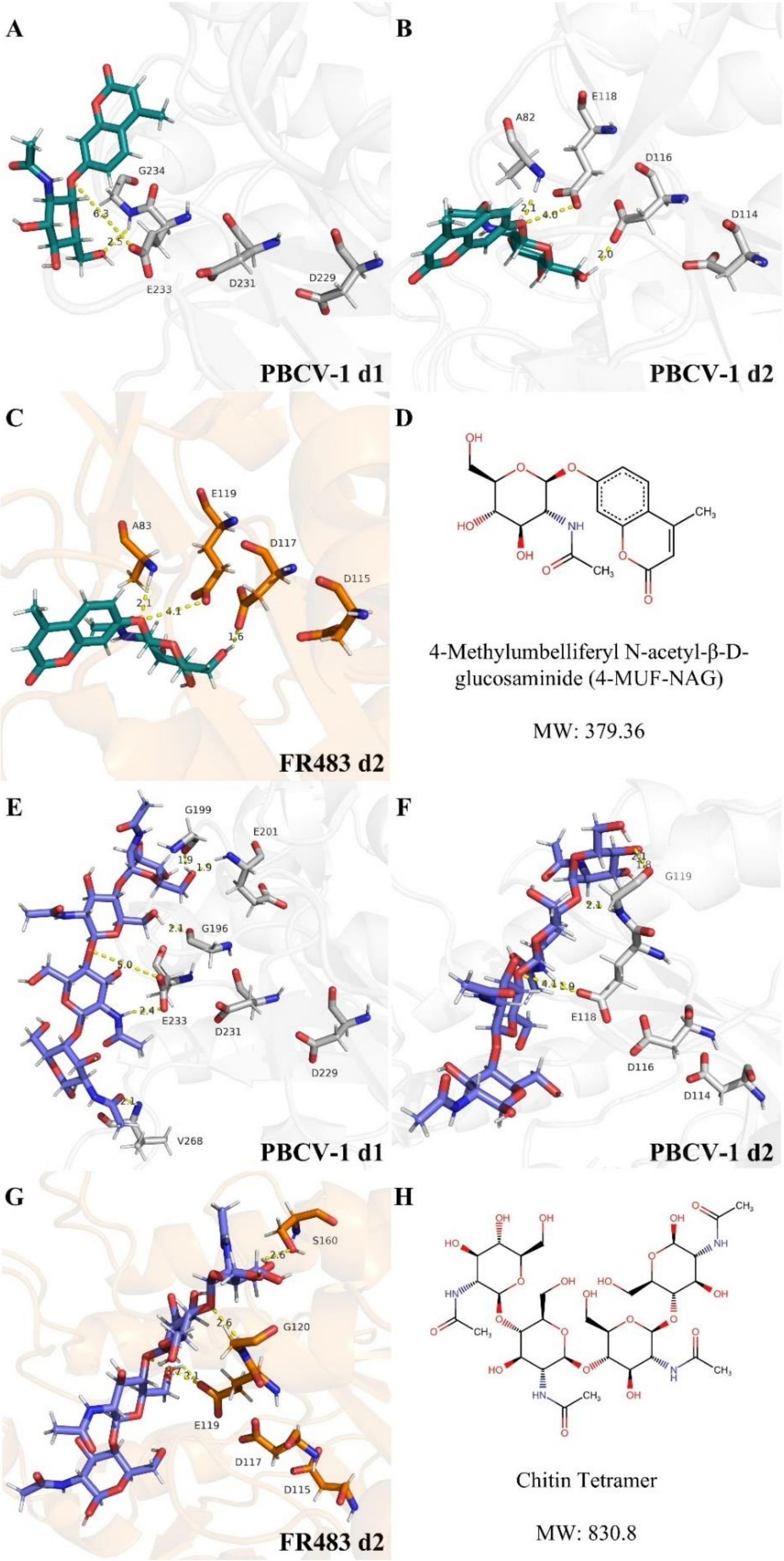

**Figure S5. Interaction of GH18 domains with 4-MUF-NAG and chitin tetramer. A:** Coupling interaction of the catalytic site (aspartate, aspartate, and glutamate) of domain 1 of PBCV-1 with 4-MUF-

NAG, domain 2 of PBCV-1 **(B)** and domain 2 of FR483 **(C)**. **D:** Chemical structure of 4-MUF-NAG. **E:** Coupling interaction of the catalytic site (aspartate, aspartate, and glutamate) of domain 1 of PBCV-1 with chitin tetramer, domain 2 of PBCV-1 **(F)** and domain 2 of FR483 **(G)**. **H:** Chemical structure of chitin tetramer.

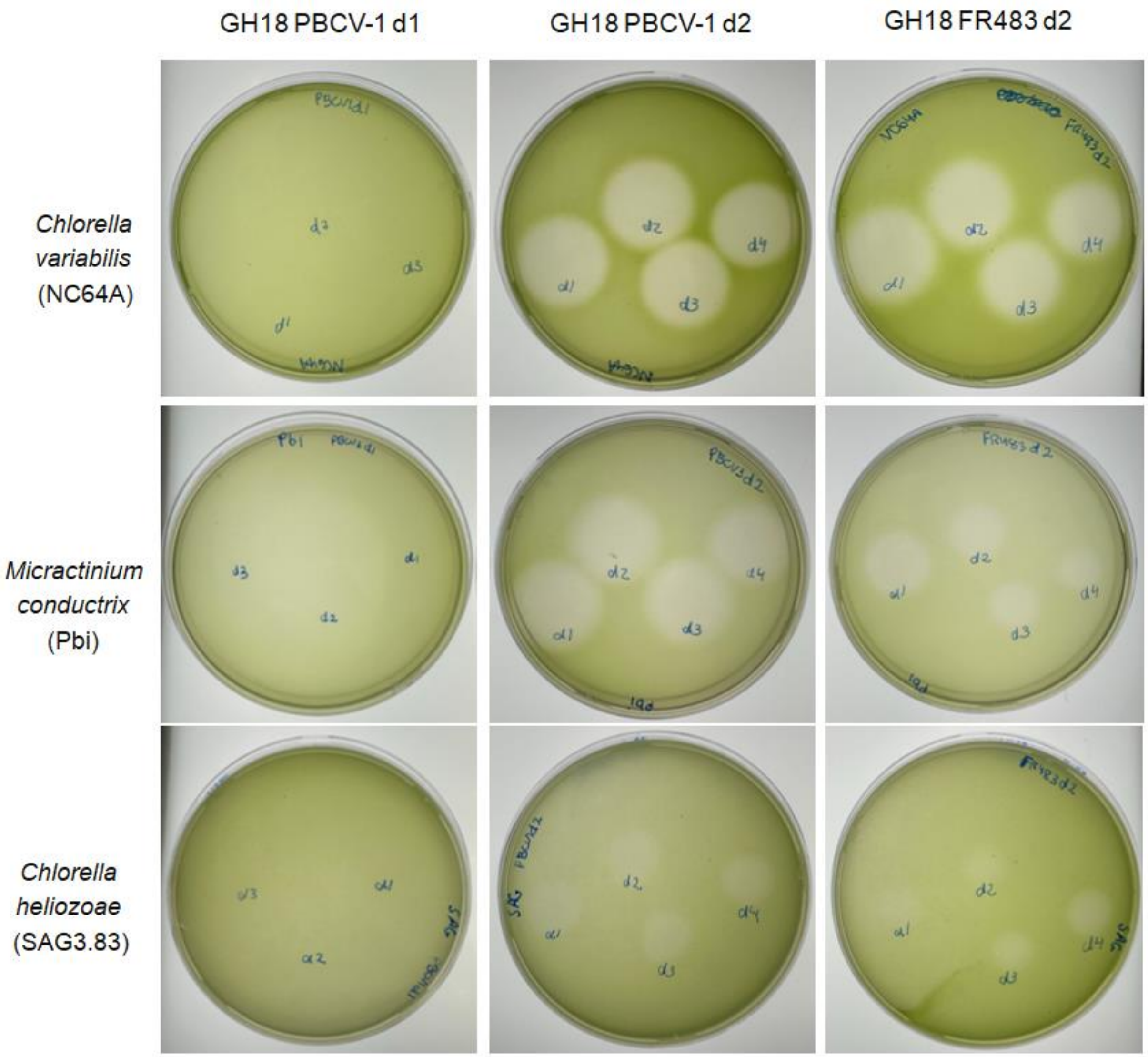

**Figure S6: Antialgal activity against *Chlorella* using agar diffusion test.** Zone of inhibition corresponds to the following concentration used, being: d1 = 7.3, d2 = 3.15, d3 = 1.5 µg for PBCV-1 d1; and d1 = 75, d2 = 37.5, d3 = 18.7, and d4 = 9 µg for PBCV-1 d2 and FR483 d2. Cells were plated in MBBM (NC64A and SAG) or FES (Pbi) agar.

**Table S1:** Distribution of CAZymes genes found in 89 chloroviruses genomes.

| Chlorovirus isolate | Cellulose synthase | Chitin synthase | Chitosanase | Chitin-binding domain | Chitinase | Aminidase | Beta-(1,3)-glucanase | N-carbamoylputrescine amidase | Hyaluronan synthase | Alginate lyase | Alginate synthase | Alpha-L-arabinofuranosidase |
|---|---|---|---|---|---|---|---|---|---|---|---|---|
| *Alphachlorovirus* (NC64A-viruses) | | | | | | | | | | | | |
| NE-40-2-s | | | 1 | 2 | 1 | 1 | 1 | 1 | | | | |
| NE-41-3-s | | | 1 | 3 | 2 | 1 | | 1 | | | | |
| CA-4A | 2 | | 1 | 2 | 1 | 1 | 1 | | 1 | | | |
| CA-4B | 2 | | 1 | 3 | 3 | 1 | 1 | | 1 | | | |
| WNE-11A-L2 | 2 | | 1 | 2 | 3 | 1 | 1 | | | | | |
| NC-1A | 1 | | 1 | 2 | 2 | 1 | 1 | | 1 | 1 | | |
| NY-2C | 2 | | 1 | 2 | 1 | 1 | 1 | 1 | 1 | | | |
| WNE-10B-S1 | 1 | 1 | 1 | | 2 | | 1 | | | | | |
| PBCV1 | | | 1 | 1 | 2 | 1 | 1 | | 1 | 1 | | |
| SH-6A | | | 1 | 2 | 3 | 1 | 1 | | 1 | | | |
| XZ-3A | | | 1 | 1 | 2 | 1 | 1 | 1 | 1 | 1 | | |
| XZ-4C | 2 | | | 2 | 3 | 1 | 1 | | 1 | | | |
| XZ-5C | | | | 2 | 2 | 1 | 1 | | 1 | | | |
| XZ-6E | 1 | | | 2 | 3 | 1 | 1 | | 1 | | | |
| 40-NE-3 | 2 | | 1 | 2 | 2 | 1 | 1 | 1 | 1 | 1 | | |
| 40-NE-4 | 2 | | 1 | 2 | 2 | 1 | 1 | 1 | 1 | | | |
| 40-NE-5 | | | 1 | 3 | 2 | 1 | | 1 | | | | |
| 41-NE-4 | | | 1 | 3 | 3 | 1 | | 1 | | | | |
| 41-NE-5 | 2 | | 1 | 2 | 2 | 1 | 1 | | 1 | 1 | | |
| 41-NE-6 | 1 | | 1 | 3 | 2 | 1 | 1 | 2 | 1 | | | |
| N-NE-4 | 1 | | 1 | 3 | 2 | 1 | 1 | 1 | | | | |
| N-NE-5 | | | 1 | 2 | 2 | 1 | | 1 | | | | |
| *Alphachlorovirus* (OSy-viruses) | | | | | | | | | | | | |
| NE-40-1-m | 1 | | 1 | 1 | 2 | 1 | 1 | 1 | 1 | | | |
| NE-41-1-L | 1 | | 1 | 1 | 2 | 1 | 1 | | 1 | | | |
| NE-41-2-m | 1 | | 1 | 2 | 2 | 1 | 1 | 1 | 1 | | | |
| NE-O-7-s | 1 | 2 | 1 | 1 | 1 | 1 | 2 | | | | | |
| NE-O-8- | 1 | | 1 | 2 | 2 | 1 | 1 | | 1 | | | |
| NE-O-9-L | 1 | | 1 | 2 | 2 | 1 | 1 | | 1 | 1 | | 2 |
| OSyNE-4B-L2 | 1 | | 1 | 1 | 2 | 1 | 1 | | 1 | 1 | | 1 |
| OSyNE-4B-M2 | 1 | | 1 | 1 | 2 | 1 | 1 | | 1 | | | 2 |
| OSyNE-4B-S1 | 1 | | 1 | 1 | 2 | | 1 | 1 | 1 | 1 | | 1 |

| | | | | | | | | | | | | |
|---|---|---|---|---|---|---|---|---|---|---|---|---|
| OSyNE-5A-L1 | 1 | | 1 | 2 | 2 | 1 | 1 | 1 | 1 | | | |
| OSyNE-5B-M2 | 1 | | 1 | 1 | 2 | 1 | 1 | 1 | 1 | | | 2 |
| OSyNE-5B-S1 | 1 | | 1 | 2 | 2 | 1 | 1 | | 1 | | | 2 |
| OSyNE-ZA | 1 | | 1 | 2 | 2 | 1 | 1 | | 1 | 1 | | |
| O-NE-10 | 1 | | 1 | 2 | 2 | 1 | 1 | 1 | 1 | | | 1 |
| O-NE-11 | 1 | | 1 | 1 | 2 | 1 | 1 | 1 | 1 | | | |
| O-NE-12 | 1 | | 1 | 2 | 2 | 1 | 1 | 1 | 1 | | | |
| O-NE-13 | 1 | | 1 | 2 | 4 | 1 | 1 | 2 | | | | |
| O-NE-14 | 1 | | 1 | 1 | 2 | 1 | 1 | 2 | 1 | | | |
| O-NE-15 | 1 | 2 | 1 | 2 | 2 | 1 | 3 | 1 | | | | |
| O-NE-16 | 1 | | 1 | 2 | 2 | | 1 | 1 | 1 | 1 | | 1 |
| O-NE-17 | 1 | | 1 | 2 | 2 | 1 | 1 | 1 | 1 | | | 2 |
| O-NE-18 | 1 | | 1 | 2 | | | 1 | 1 | 1 | 1 | | 2 |
| O-NE-19 | 1 | 2 | 1 | 1 | | 1 | 2 | 1 | | | | |
| O-NE-20 | 1 | | 1 | 2 | 2 | 1 | 1 | 1 | 1 | 1 | | 2 |
| O-NE-22 | 1 | 2 | 1 | 1 | 1 | 1 | 2 | 1 | | | | |
| O-NE-23 | 1 | 2 | 1 | 1 | 1 | 1 | 2 | 1 | | 1 | | |
| O-NE-24 | 2 | | 1 | 1 | 3 | 1 | 2 | 1 | 1 | | | 2 |
| O-NE-25 | 1 | 2 | 1 | 1 | 1 | 1 | 2 | 1 | | | | |
| O-NE-26 | 1 | | 1 | 2 | 2 | 1 | 1 | 1 | 1 | | | |
| O-NE-27 | 1 | 1 | 1 | 1 | 1 | 1 | 1 | 1 | | 1 | | |
| O-NE-28 | 1 | | 1 | 2 | 3 | 1 | | 1 | | 1 | | |
| O-NE-29 | 1 | 2 | 1 | 1 | 1 | 1 | 2 | 1 | | | | |
| *Gammachlorovirus* | | | | | | | | | | | | |
| CL-S-1-m | 8 | 1 | 1 | 2 | 2 | 1 | 2 | | | | 1 | |
| GNLD-22 | 4 | 1 | 1 | 1 | 2 | 1 | 1 | | | | 1 | |
| LP-F3a-4a | 3 | | 1 | 2 | 1 | 1 | | | | | | |
| NES-5A-L1 | 4 | 1 | 1 | 1 | 2 | 1 | 1 | | | | 1 | |
| NES-5A-M1 | 3 | 1 | 1 | 1 | 2 | 1 | 1 | | | | 1 | |
| NES-5A-S1 | 7 | | | 2 | 2 | 1 | 2 | | | | | |
| NES-4B-L1 | 2 | | | 2 | 2 | 1 | | | | | 1 | |
| NES-4A-M1 | | 1 | 1 | 1 | 1 | 1 | 1 | | | | 1 | |
| NES-4A-S1 | 2 | 1 | 1 | 1 | 3 | 1 | 1 | | | | 1 | |
| S-NE-7 | 3 | 1 | 1 | 2 | 3 | 1 | 4 | 1 | | | 1 | |
| S-NE-9 | 3 | 1 | 1 | 2 | 2 | | 1 | | | | | |
| S-NE-10 | 4 | | | 2 | 1 | 1 | 1 | 1 | | | | |
| S-NE-11 | 3 | 1 | 1 | 1 | 2 | | | | | | | |
| S-NE-12 | 6 | | | 2 | 3 | | 4 | 1 | | | | |
| S-NE-13 | 2 | 1 | 1 | 2 | 1 | 1 | 3 | | | | 1 | |
| S-NE-15 | 5 | | | 2 | 3 | | 2 | | | | | |
| S-NE-16 | 4 | | | 2 | 1 | | 2 | | | | | |
| S-NE-17 | 6 | | | 2 | 2 | | 2 | 1 | | | | |
| S-NE-18 | 6 | 1 | 1 | 1 | 2 | 1 | 2 | 1 | | | 1 | |
| S-NE-19 | 5 | | | 2 | 3 | | 3 | 1 | | | | |
| S-NE-20 | 6 | 1 | 1 | 3 | 3 | 1 | 2 | 1 | | 1 | | |
| S-NE-22 | 2 | 1 | 1 | 1 | 2 | 1 | 1 | 1 | | | 1 | |

| | | | | | | | | | | | |
|---|---|---|---|---|---|---|---|---|---|---|---|
| S-NE-23 | 4 | | 1 | 2 | 2 | | 1 | 1 | | | |
| *Betachlorovirus* | | | | | | | | | | | |
| NE-P-1-L | 1 | | | 1 | 2 | 1 | 1 | | 1 | | |
| NE-P-2-m | 1 | | 1 | 1 | 2 | 1 | 1 | | | | |
| NE-P-4-fs | 1 | | 1 | 1 | 2 | 1 | 1 | | | | |
| NE-P-3-s | 1 | | 1 | 1 | 2 | 1 | 1 | | 1 | | |
| P-NE-9 | 1 | | 1 | 1 | 3 | | 1 | 1 | | | |
| P-NE-10 | | | 1 | 1 | 2 | | 1 | 1 | | | |
| P-NE-11 | 1 | | 1 | 1 | 2 | 1 | 1 | 1 | 1 | | |
| P-NE-12 | 1 | | 1 | 1 | 2 | 1 | 1 | 1 | 1 | | |
| P-NE-13 | 1 | | 1 | 1 | 2 | 1 | 1 | 1 | 1 | | |
| NE-P-6-s | 1 | 2 | 1 | 1 | 3 | 1 | 1 | 2 | | | 1 |
| NE-P-8-f | 1 | | 1 | 1 | 2 | 1 | 1 | 1 | 1 | | |

**Table S2:** X-ray diffraction data collection, data processing, and crystal structure refinement statistics for the catalytic domain 2 of FR483 betachlorovirus (PDB ID:37CA).

| | |
|---|---|
| | ***Data collection*** |
| Location | The Brazilian Synchrotron Light Laboratory - Sirius |
| Beamline | Macromolecular Micro and Nano Crystallography (MANACÁ) |
| Detector | Pilatus 2M |
| Wavelength (Å) | 0.77490 |
| Detector distance (mm) | 112 |
| Rotation per image (deg) | 0.1 |
| Total rotation range (deg) | 360 |
| ***Data processing*** | |
| Space group | $P2_12_12_1$ |
| a, b & c (Å) | 42.33, 80.28, 80.75 |
| Mosaicity (deg) | 0.169 |
| Resolution range (Å) | 80.75 – 1.04 (1.11 – 1.04)[a] |
| Number of accepted reflections | 1,663,849 (264,222) |
| Number of accepted unique reflections | 130,496 (20,226) |
| Redundancy | 12.8 (13.1) |
| Completeness (%) | 99.4 (96.2) |
| <I/σ(I)> | 16.2 (1.1) |
| R-meas (%)[b] | 10.6 (250.2) |
| CC(1/2)[c] | 99.9 (45.7) |
| Estimated Matthews Coeff. (Å$^3$.Da$^{-1}$)[d] | 2.2 |
| Estimated solvent content (%)[d] | 44 |
| ***Refinement statistics*** | |
| R work (%) | 15.6 |
| R free (%)[e] | 17.5 |
| Number of protein atoms[f] | 2,190 |
| Number of water molecules | 380 |
| Protein atoms average B factor (Å$^2$) | 12.8 |
| Water molecules average B factor (Å$^2$) | 24.2 |
| Bond angles RMSD from ideal (deg) | 1.827 |
| Bond lengths RMSD from ideal (Å) | 0.025 |
| Ramachandran's favored regions (%) | 97.8 |
| Ramachandran's allowed regions (%) | 1.8 |
| Ramachandran's outliers (%) | 0.4 |

[a] Values in parentheses correspond to the highest resolution shell

[b] Redundancy-independent R factor based on intensities, according to Diederichs and Karplus (1997)

[c] Percentage correlation between intensities from randomly selected half-datasets, according to Karplus and Diederichs (2012)

[d] Estimated values based on the presence of 1 molecule of the catalytic domain 2 of FR483 betachlorovirus: in the asymmetric unit of the crystal

[e] R free is the *R*-factor calculated from 5% of reflections (6,521 reflections) excluded from structure refinement

[f] Hydrogen atoms were not included in the calculation, whereas atoms with alternative conformations were counted

**Table S3:** Prevalence of coevolving residues on chlorovirus chitinases in set 1. Colored in green represent coevolved residues.

| Sequence | D1755 | D543 | F477 | G383 | F83 | G387 | W562 | H73 | T72 | Y5 |
|---|---|---|---|---|---|---|---|---|---|---|
| WZO81342.1/1-840bifunctionalchitinase/lysozyme\[CA-4B]/142-304 | - | D88 | L71 | G54 | F13 | G55 | I91 | F8 | K7 | - |
| WZO81342.1/1-840bifunctionalchitinase/lysozyme\[CA-4B]/573-747 | - | D97 | L80 | G63 | Y28 | G64 | I100 | L23 | T22 | Y3 |
| WZO81006.1/1-840putativebifunctionalchitinase/lysozyme\[CA-4A]/142-304 | - | D88 | L71 | G54 | F13 | G55 | I91 | F8 | K7 | - |
| WZO81006.1/1-840putativebifunctionalchitinase/lysozyme\[CA-4A]/573-747 | - | D97 | L80 | G63 | Y28 | G64 | I100 | L23 | T22 | Y3 |
| WZO82266.1/1-840putativebifunctionalchitinase/lysozyme\[NY-2C]/142-304 | - | D88 | L71 | G54 | F13 | G55 | I91 | F8 | K7 | - |
| WZO82266.1/1-840putativebifunctionalchitinase/lysozyme\[NY-2C]/573-747 | - | D97 | L80 | G63 | Y28 | G64 | I100 | L23 | T22 | Y3 |
| WZO80816.1/1-830putativebifunctionalchitinase/lysozyme\[WNE-11A-L2]/142-304 | - | D88 | L71 | G54 | F13 | G55 | I91 | F8 | K7 | - |
| WZO80816.1/1-830putativebifunctionalchitinase/lysozyme\[WNE-11A-L2]/563-740 | - | D97 | L80 | G63 | Y28 | G64 | I100 | L23 | T22 | Y3 |
| AGE57241.1/1-815putativebifunctionalchitinase/lysozyme\[ParameciumbursariaChlorellavirusNE-JV-4]/142-304 | - | D88 | L71 | G54 | F13 | G55 | I91 | F8 | K7 | - |
| AGE57241.1/1-815putativebifunctionalchitinase/lysozyme\[ParameciumbursariaChlorellavirusNE-JV-4]/548-725 | - | D97 | L80 | G63 | Y28 | G64 | I100 | L23 | T22 | Y3 |
| WZO75107.1/1-840chitinase\[40-NE-3]/142-304 | - | D88 | L71 | G54 | F13 | G55 | I91 | F8 | K7 | - |
| WZO75107.1/1-840chitinase\[40-NE-3]/573-747 | - | D97 | L80 | G63 | Y28 | G64 | I100 | L23 | T22 | Y3 |
| WZO75726.1/1-840putativebifunctionalchitinase/lysozyme\[40-NE-4]/142-304 | - | D88 | L71 | G54 | F13 | G55 | I91 | F8 | K7 | - |
| WZO75726.1/1-840putativebifunctionalchitinase/lysozyme\[40-NE-4]/573-747 | - | D97 | L80 | G63 | Y28 | G64 | I100 | L23 | T22 | Y3 |
| WZO76741.1/1-840chitinase\[41-NE-5]/142-304 | - | D88 | L71 | G54 | F13 | G55 | I91 | F8 | K7 | - |
| WZO76741.1/1-840chitinase\[41-NE-5]/573-747 | - | D97 | L80 | G63 | Y28 | G64 | I100 | L23 | T22 | Y3 |
| WZO83158.1/1-830chitinase\[XZ-4C]/142-304 | - | D88 | L71 | G54 | F13 | G55 | I91 | F8 | K7 | - |
| WZO83158.1/1-830chitinase\[XZ-4C]/563-737 | - | D97 | L80 | G63 | Y28 | G64 | I100 | L23 | T22 | Y3 |
| AGE53811.1/1-835putativebifunctionalchitinase/lysozyme\[ParameciumbursariaChlorellavirusIL3A]/142-304 | - | D88 | L71 | G54 | F13 | G55 | I91 | F8 | K7 | - |
| AGE53811.1/1-835putativebifunctionalchitinase/lysozyme\[ParameciumbursariaChlorellavirusIL3A]/567-745 | - | D98 | L81 | G64 | Y29 | G65 | I101 | L24 | T23 | Y4 |
| AGE54506.1/1-817putativebifunctionalchitinase/lysozyme\[ParameciumbursariaChlorellavirusKS1B]/142-304 | - | D88 | L71 | G54 | F13 | G55 | I91 | F8 | K7 | - |
| AGE54506.1/1-817putativebifunctionalchitinase/lysozyme\[ParameciumbursariaChlorellavirusKS1B]/550-727 | - | D97 | L80 | G63 | Y28 | G64 | I100 | L23 | T22 | Y3 |
| AGE48399.1/1-835putativebifunctionalchitinase/lysozyme\[ParameciumbursariaChlorellavirusAN69C]/142-304 | - | D88 | L71 | G54 | F13 | G55 | I91 | F8 | K7 | - |
| AGE48399.1/1-835putativebifunctionalchitinase/lysozyme\[ParameciumbursariaChlorellavirusAN69C]/568-742 | - | D97 | L80 | G63 | Y28 | G64 | I100 | L23 | T22 | Y3 |
| WZO91574.1/1-809putativebifunctionalchitinase/lysozyme\[O-NE-29]/139-304 | - | D91 | L74 | G57 | F16 | G58 | I94 | F11 | K10 | - |
| WZO91574.1/1-809putativebifunctionalchitinase/lysozyme\[O-NE-29]/542-716 | - | D97 | L80 | G63 | Y28 | G64 | I100 | L23 | T22 | F3 |
| WZO81686.1/1-840chitinase\[NC-1A]/142-304 | - | D88 | L71 | G54 | F13 | G55 | I91 | F8 | K7 | - |
| WZO81686.1/1-840chitinase\[NC-1A]/573-747 | - | D97 | L80 | G63 | Y28 | G64 | I100 | L23 | T22 | Y3 |
| NP_048529.2/1-830Chitinase\[ParameciumbursariaChlorellavirus1]/142-304 | - | D88 | L71 | G54 | F13 | G55 | I91 | F8 | K7 | - |
| NP_048529.2/1-830Chitinase\[ParameciumbursariaChlorellavirus1]/563-737 | - | D97 | L80 | G63 | Y28 | G64 | I100 | L23 | T22 | Y3 |
| WZO82806.1/1-830chitinase\[XZ-3A]/142-304 | - | D88 | L71 | G54 | F13 | G55 | I91 | F8 | K7 | - |
| WZO82806.1/1-830chitinase\[XZ-3A]/563-737 | - | D97 | L80 | G63 | Y28 | G64 | I100 | L23 | T22 | Y3 |
| WZO90762.1/1-809putativebifunctionalchitinase/lysozyme\[O-NE-27]/139-304 | - | D91 | L74 | G57 | F16 | G58 | I94 | F11 | K10 | - |
| WZO90762.1/1-809putativebifunctionalchitinase/lysozyme\[O-NE-27]/542-716 | - | D97 | L80 | G63 | Y28 | G64 | I100 | L23 | T22 | F3 |
| WZO83510.1/1-820putativebifunctionalchitinase/lysozyme\[XZ-5C]/142-304 | - | D88 | L71 | G54 | F13 | G55 | I91 | F8 | K7 | - |
| WZO83510.1/1-820putativebifunctionalchitinase/lysozyme\[XZ-5C]/553-731 | - | D97 | L80 | G63 | Y28 | G64 | I100 | L23 | S22 | Y3 |
| WZO89952.1/1-822putativebifunctionalchitinase/lysozyme\[O-NE-25]/142-304 | - | D88 | L71 | G54 | F13 | G55 | I91 | F8 | K7 | - |
| WZO89952.1/1-822putativebifunctionalchitinase/lysozyme\[O-NE-25]/555-729 | - | D97 | L80 | G63 | S28 | G64 | I100 | L23 | T22 | F3 |
| AGE51440.1/1-836putativebifunctionalchitinase/lysozyme\[ParameciumbursariaChlorellavirusCviKI]/142-304 | - | D88 | L71 | G54 | F13 | G55 | I91 | F8 | K7 | - |
| AGE51440.1/1-836putativebifunctionalchitinase/lysozyme\[ParameciumbursariaChlorellavirusCviKI]/569-748 | - | D97 | L80 | G63 | Y28 | G64 | I100 | L23 | T22 | F3 |
| AGE52455.1/1-826putativebifunctionalchitinase/lysozyme\[ParameciumbursariaChlorellavirusCvsA1]/142-304 | - | D88 | L71 | G54 | F13 | G55 | I91 | F8 | K7 | - |
| AGE52455.1/1-826putativebifunctionalchitinase/lysozyme\[ParameciumbursariaChlorellavirusCvsA1]/559-739 | - | D97 | L80 | G63 | Y28 | G64 | I100 | L23 | T22 | F3 |
| WZO87987.1/1-821putativebifunctionalchitinase/lysozyme\[O-NE-19]/142-304 | - | D88 | L71 | G54 | F13 | G55 | I91 | F8 | K7 | - |
| WZO87987.1/1-821putativebifunctionalchitinase/lysozyme\[O-NE-19]/554-728 | - | D97 | L80 | G63 | S28 | G64 | I100 | L23 | T22 | F3 |
| WZO82436.1/1-820putativebifunctionalchitinase/lysozyme\[SH-6A]/142-304 | - | D88 | L71 | G54 | F13 | G55 | I91 | F8 | K7 | - |
| WZO82436.1/1-820putativebifunctionalchitinase/lysozyme\[SH-6A]/553-731 | - | D97 | L80 | G63 | Y28 | G64 | I100 | L23 | S22 | Y3 |
| WZO92132.1/1-827putativebifunctionalchitinase/lysozyme\[NE-O-7-s]/139-304 | - | D91 | L74 | G57 | F16 | G58 | I94 | F11 | K10 | - |
| WZO92132.1/1-827putativebifunctionalchitinase/lysozyme\[NE-O-7-s]/560-734 | - | D97 | L80 | G63 | Y28 | G64 | I100 | L23 | T22 | F3 |
| WZO89127.1/1-809putativebifunctionalchitinase/lysozyme\[O-NE-23]/139-304 | - | D91 | L74 | G57 | F16 | G58 | I94 | F11 | K10 | - |
| WZO89127.1/1-809putativebifunctionalchitinase/lysozyme\[O-NE-23]/542-716 | - | D97 | L80 | G63 | S28 | G64 | I100 | L23 | T22 | F3 |

| | | | | | | | | | | |
|---|---|---|---|---|---|---|---|---|---|---|
| WZO85876.1/1-819putativebifunctionalchitinase/lysozyme\[O-NE-14]/142-304 | - | D88 | L71 | G54 | F13 | G55 | I91 | F8 | K7 | - |
| WZO85876.1/1-819putativebifunctionalchitinase/lysozyme\[O-NE-14]/552-726 | - | D97 | L80 | G63 | Y28 | G64 | I100 | L23 | T22 | Y3 |
| WZO88848.1/1-822putativebifunctionalchitinase/lysozyme\[O-NE-22]/142-304 | - | D88 | L71 | G54 | F13 | G55 | I91 | F8 | K7 | - |
| WZO88848.1/1-822putativebifunctionalchitinase/lysozyme\[O-NE-22]/555-729 | - | D97 | L80 | G63 | S28 | G64 | I100 | L23 | T22 | F3 |
| WZO83897.1/1-820putativebifunctionalchitinase/lysozyme\[XZ-6E]/142-304 | - | D88 | L71 | G54 | F13 | G55 | I91 | F8 | K7 | - |
| WZO83897.1/1-820putativebifunctionalchitinase/lysozyme\[XZ-6E]/553-732 | - | D97 | L80 | G63 | Y28 | G64 | I100 | L23 | S22 | Y3 |
| YP_009665302.1/1-806putativebifunctionalchitinase/lysozyme\[ParameciumbursariaChlorellavirusNYs1]/142-304 | - | D88 | L71 | G54 | F13 | G55 | I91 | F8 | K7 | - |
| YP_009665302.1/1-806putativebifunctionalchitinase/lysozyme\[ParameciumbursariaChlorellavirusNYs1]/539-715 | - | D97 | L80 | G63 | S28 | G64 | I100 | L23 | T22 | F3 |
| WZO87550.1/1-808putativebifunctionalchitinase/lysozyme\[O-NE-18]/143-303 | - | D86 | L69 | G52 | F11 | G53 | I89 | F6 | K5 | - |
| WZO87550.1/1-808putativebifunctionalchitinase/lysozyme\[O-NE-18]/541-715 | - | D97 | L80 | G63 | S28 | G64 | I100 | L23 | T22 | F3 |
| AGE54806.1/1-806putativebifunctionalchitinase/lysozyme\[ParameciumbursariaChlorellavirusMA1D]/142-304 | - | D88 | L71 | G54 | F13 | G55 | I91 | F8 | K7 | - |
| AGE54806.1/1-806putativebifunctionalchitinase/lysozyme\[ParameciumbursariaChlorellavirusMA1D]/539-715 | - | D97 | L80 | G63 | S28 | G64 | I100 | L23 | T22 | F3 |
| WZO84706.1/1-831putativebifunctionalchitinase/lysozyme\[O-NE-11]/142-304 | - | D88 | L71 | G54 | F13 | G55 | I91 | F8 | K7 | - |
| WZO84706.1/1-831putativebifunctionalchitinase/lysozyme\[O-NE-11]/564-738 | - | D97 | L80 | G63 | Y28 | G64 | I100 | L23 | T22 | Y3 |
| WZO80449.1/1-806putativebifunctionalchitinase/lysozyme\[WNE-10B-S1]/142-304 | - | D88 | L71 | G54 | F13 | G55 | I91 | F8 | K7 | - |
| WZO80449.1/1-806putativebifunctionalchitinase/lysozyme\[WNE-10B-S1]/539-714 | - | D97 | L80 | G63 | S28 | G64 | I100 | L23 | T22 | F3 |
| WZO85478.1/1-830putativebifunctionalchitinase/lysozyme\[O-NE-13]/142-304 | - | D88 | L71 | G54 | F13 | G55 | I91 | F8 | K7 | - |
| WZO85478.1/1-830putativebifunctionalchitinase/lysozyme\[O-NE-13]/564-737 | - | D96 | L79 | G62 | S27 | G63 | I99 | L22 | T21 | F2 |
| WZO76146.1/1-816bifunctionalchitinase/lysozyme\[40-NE-5]/142-304 | - | D88 | L71 | G54 | F13 | G55 | I91 | F8 | K7 | - |
| WZO76146.1/1-816bifunctionalchitinase/lysozyme\[40-NE-5]/549-723 | - | D97 | L80 | G63 | F25 | G64 | I100 | N20 | N19 | F3 |
| WZO76561.1/1-816putativebifunctionalchitinase/lysozyme\[41-NE-4]/142-304 | - | D88 | L71 | G54 | F13 | G55 | I91 | F8 | K7 | - |
| WZO76561.1/1-816putativebifunctionalchitinase/lysozyme\[41-NE-4]/549-723 | - | D97 | L80 | G63 | F25 | G64 | I100 | N20 | N19 | F3 |
| WZO78129.1/1-816chitinase\[NE-40ã??2-s]/142-304 | - | D88 | L71 | G54 | F13 | G55 | I91 | F8 | K7 | - |
| WZO78129.1/1-816chitinase\[NE-40ã??2-s]/549-723 | - | D97 | L80 | G63 | F25 | G64 | I100 | N20 | N19 | F3 |
| WZO79003.1/1-816putativebifunctionalchitinase/lysozyme\[NE-41-3-s]/142-304 | - | D88 | L71 | G54 | F13 | G55 | I91 | F8 | K7 | - |
| WZO79003.1/1-816putativebifunctionalchitinase/lysozyme\[NE-41-3-s]/549-723 | - | D97 | L80 | G63 | F25 | G64 | I100 | N20 | N19 | F3 |
| WZO80062.1/1-823putativebifunctionalchitinase/lysozyme\[N-NE-5]/142-304 | - | D88 | L71 | G54 | F13 | G55 | I91 | F8 | K7 | - |
| WZO80062.1/1-823putativebifunctionalchitinase/lysozyme\[N-NE-5]/549-723 | - | D97 | L80 | G63 | F25 | G64 | I100 | N20 | N19 | F3 |
| AGE58174.1/1-816putativebifunctionalchitinase/lysozyme\[ParameciumbursariaChlorellavirusNW665.2]/140-305 | - | D91 | L74 | G57 | F16 | G58 | I94 | F11 | K10 | - |
| AGE58174.1/1-816putativebifunctionalchitinase/lysozyme\[ParameciumbursariaChlorellavirusNW665.2]/550-723 | - | D96 | L79 | G62 | S27 | G63 | I99 | L22 | T21 | F2 |
| YP_001426411.1/1-805hypotheticalproteinFR483_N779R\[ParameciumbursariaChlorellavirusFR483]/139-304 | - | D91 | L74 | G57 | F16 | G58 | I94 | F11 | K10 | - |
| YP_001426411.1/1-805hypotheticalproteinFR483_N779R\[ParameciumbursariaChlorellavirusFR483]/539-712 | - | D96 | L79 | G62 | S27 | G63 | I99 | L22 | T21 | F2 |
| WZO86764.1/1-525chitinase\[O-NE-16]/142-304 | - | D88 | L71 | G54 | F13 | G55 | I91 | F8 | K7 | - |
| WZO93853.1/1-525chitinase\[OSyNE-4B-S1]/142-304 | - | D88 | L71 | G54 | F13 | G55 | I91 | F8 | K7 | - |
| WZO91298.1/1-445chitinase\[O-NE-28]/147-309 | - | D88 | L71 | G54 | F13 | G55 | I91 | F8 | K7 | - |
| WZO89577.1/1-445chitinase\[O-NE-24]/147-309 | - | D88 | L71 | G54 | F13 | G55 | I91 | F8 | K7 | - |
| WZO90501.1/1-525chitinase\[O-NE-26]/142-304 | - | D88 | L71 | G54 | F13 | G55 | I91 | F8 | K7 | - |
| WZO77210.1/1-525chitinase\[41-NE-6]/142-304 | - | D88 | L71 | G54 | F13 | G55 | I91 | F8 | K7 | - |
| WZO77693.1/1-525chitinase\[NE-40-1-m]/142-304 | - | D88 | L71 | G54 | F13 | G55 | I91 | F8 | K7 | - |
| WZO78681.1/1-525chitinase\[NE-41-2-m]/142-304 | - | D88 | L71 | G54 | F13 | G55 | I91 | F8 | K7 | - |
| WZO85099.1/1-530chitinase\[O-NE-12]/147-309 | - | D88 | L71 | G54 | F13 | G55 | I91 | F8 | K7 | - |
| YP_009325635.1/1-525Chitinase\[OnlySyngenNebraskavirus5]/142-304 | - | D88 | L71 | G54 | F13 | G55 | I91 | F8 | K7 | - |
| WZO88270.1/1-528chitinase\[O-NE-20]/147-309 | - | D88 | L71 | G54 | F13 | G55 | I91 | F8 | K7 | - |
| WZO92747.1/1-528chitinase\[NE-O-9-L]/147-309 | - | D88 | L71 | G54 | F13 | G55 | I91 | F8 | K7 | - |
| WZO93306.1/1-528chitinase\[OSyNE-4B-L2]/147-309 | - | D88 | L71 | G54 | F13 | G55 | I91 | F8 | K7 | - |
| WZO87136.1/1-528chitinase\[O-NE-17]/147-309 | - | D88 | L71 | G54 | F13 | G55 | I91 | F8 | K7 | - |
| WZO92491.1/1-528chitinase\[NE-O-8-L]/147-309 | - | D88 | L71 | G54 | F13 | G55 | I91 | F8 | K7 | - |
| WZO95440.1/1-528chitinase\[OSyNE-ZA]/147-309 | - | D88 | L71 | G54 | F13 | G55 | I91 | F8 | K7 | - |
| WZO78440.1/1-532chitinase\[NE-41-1-L]/147-309 | - | D88 | L71 | G54 | F13 | G55 | I91 | F8 | K7 | - |
| WZO94180.1/1-539chitinase\[OSyNE-5A-L1]/147-309 | - | D88 | L71 | G54 | F13 | G55 | I91 | F8 | K7 | - |
| WZO84329.1/1-525chitinase\[O-NE-10]/142-304 | - | D88 | L71 | G54 | F13 | G55 | I91 | F8 | K7 | - |
| WZO94943.1/1-525chitinase\[OSyNE-5B-S1]/142-304 | - | D88 | L71 | G54 | F13 | G55 | I91 | F8 | K7 | - |
| WZO93480.1/1-525chitinase\[OSyNE-4B-M2]/142-304 | - | D88 | L71 | G54 | F13 | G55 | I91 | F8 | K7 | - |
| WZO94540.1/1-525chitinase\[OSyNE-5B-M2]/142-304 | - | D88 | L71 | G54 | F13 | G55 | I91 | F8 | K7 | - |
| WZO79465.1/1-530chitinase\[N-NE-4]/147-309 | - | D88 | L71 | G54 | F13 | G55 | I91 | F8 | K7 | - |
| XYW60054.1/1-521chitinase\[NE-P-3-s]/139-304 | - | D91 | L74 | G57 | F16 | G58 | I94 | F11 | K10 | - |
| XYW60732.1/1-521chitinase\[NE-P-4-fs]/139-304 | - | D91 | L74 | G57 | F16 | G58 | I94 | F11 | K10 | - |
| XYW61232.1/1-521chitinase\[NE-P-8-f]/139-304 | - | D91 | L74 | G57 | F16 | G58 | I94 | F11 | K10 | - |
| XYW62772.1/1-521chitinase\[P-NE-12]/139-304 | - | D91 | L74 | G57 | F16 | G58 | I94 | F11 | K10 | - |
| WZO86398.1/1-532chitinase\[O-NE-15]/144-309 | - | D91 | L74 | G57 | F16 | G58 | I94 | F11 | K10 | - |
| XOK27477.1/1-582putativebifunctionalchitinase/lysozyme\[CL-S-1-m]/153-304 | - | D77 | L60 | G43 | F2 | G44 | I80 | - | - | - |
| AGE49332.1/1-595putativebifunctionalchitinase/lysozyme\[AcanthocystisturfaceaChlorellavirusBr0604L]/166-317 | - | D77 | L60 | G43 | F2 | G44 | I80 | - | - | - |
| AGE48989.1/1-520putativebifunctionalchitinase/lysozyme\[ParameciumbursariaChlorellavirusAP110A]/139-304 | - | D91 | L74 | G57 | F16 | G58 | I94 | F11 | K10 | - |
| AGE50008.1/1-520putativebifunctionalchitinase/lysozyme\[ParameciumbursariaChlorellavirusCan18-4]/139-304 | - | D91 | L74 | G57 | F16 | G58 | I94 | F11 | K10 | - |
| XYW62298.1/1-520chitinase\[P-NE-10]/139-304 | - | D91 | L74 | G57 | F16 | G58 | I94 | F11 | K10 | - |
| AGE52349.1/1-523putativebifunctionalchitinase/lysozyme\[ParameciumbursariaChlorellavirusCVR-1]/143-304 | - | D87 | L70 | G53 | F12 | G54 | I90 | F7 | K6 | - |
| YP_009702007.1/1-523putativebifunctionalchitinase/lysozyme\[ParameciumbursariaChlorellavirusCVA-1]/143-304 | - | D87 | L70 | G53 | F12 | G54 | I90 | F7 | K6 | - |
| ABT14345.1/1-520hypotheticalproteinMT325_M791R\[ParameciumbursariachlorellavirusMT325]/139-304 | - | D91 | L74 | G57 | F16 | G58 | I94 | F11 | K10 | - |

| | | | | | | | | | | |
|---|---|---|---|---|---|---|---|---|---|---|
| AGE51335.1/1-523putativebifunctionalchitinase/lysozyme\[ParameciumbursariaChlorellavirusCVG-1]/139-304 | - | D91 | L74 | G57 | F16 | G58 | I94 | F11 | K10 | - |
| XYW61920.1/1-523chitinase\[P-NE-9]/139-304 | - | D91 | L74 | G57 | F16 | G58 | I94 | F11 | K10 | - |
| XOK31597.1/1-580putativebifunctionalchitinase/lysozyme\[S-NE-11]/151-304 | - | D79 | L62 | G45 | F4 | G46 | I82 | - | - | - |
| XOK36306.1/1-580putativebifunctionalchitinase/lysozyme\[GNLD22]/152-304 | - | D78 | L61 | G44 | F3 | G45 | I81 | - | - | - |
| XOK36264.1/1-582putativebifunctionalchitinase/lysozyme\[LP-F3a-4a]/153-304 | - | D77 | L60 | G43 | F2 | G44 | I80 | - | - | - |
| XOK31528.1/1-582putativebifunctionalchitinase/lysozyme\[S-NE-10]/153-304 | - | D77 | L60 | G43 | F2 | G44 | I80 | - | - | - |
| XOK28546.1/1-582putativebifunctionalchitinase/lysozyme\[NES-4B-L1]/153-304 | - | D77 | L60 | G43 | F2 | G44 | I80 | - | - | - |
| YP_001427295.1/1-595hypotheticalproteinATCV1_Z814L\[Acanthocystisturfaceachlorellavirus1]/166-317 | - | D77 | L60 | G43 | F2 | G44 | I80 | - | - | - |
| AGE56132.1/1-601putativebifunctionalchitinase/lysozyme\[AcanthocystisturfaceaChlorellavirusMO0605SPH]/172-323 | - | D77 | L60 | G43 | F2 | G44 | I80 | - | - | - |
| XOK34753.1/1-421putativebifunctionalchitinase/lysozyme\[S-NE-19]/153-304 | - | D77 | L60 | G43 | F2 | G44 | I80 | - | - | - |
| AGE60252.1/1-601putativebifunctionalchitinase/lysozyme\[AcanthocystisturfaceaChlorellavirusWI0606]/172-323 | - | D77 | L60 | G43 | F2 | G44 | I80 | - | - | - |
| AGE57134.1/1-601putativebifunctionalchitinase/lysozyme\[AcanthocystisturfaceaChlorellavirusNE-JV-3]/172-323 | - | D77 | L60 | G43 | F2 | G44 | I80 | - | - | - |
| XOK29995.1/1-582putativebifunctionalchitinase/lysozyme\[NES-5A-S1]/153-304 | - | D77 | L60 | G43 | F2 | G44 | I80 | - | - | - |
| AGE49664.1/1-595putativebifunctionalchitinase/lysozyme\[AcanthocystisturfaceaChlorellavirusCan0610SP]/166-317 | - | D77 | L60 | G43 | F2 | G44 | I80 | - | - | - |
| AGE59935.1/1-595putativebifunctionalchitinase/lysozyme\[AcanthocystisturfaceaChlorellavirusTN603.4.2]/189-317 | - | D54 | L37 | G20 | - | G21 | I57 | - | - | - |
| AGE55823.1/1-596putativebifunctionalchitinase/lysozyme\[AcanthocystisturfaceaChlorellavirusMN0810.1]/185-317 | - | D58 | L41 | G24 | - | G25 | I61 | - | - | - |
| XOK35208.1/1-580putativebifunctionalchitinase/lysozyme\[S-NE-22]/149-304 | - | D81 | L64 | G47 | F6 | G48 | I84 | F1 | - | - |
| AGE50333.1/1-580putativebifunctionalchitinase/lysozyme\[AcanthocystisturfaceaChlorellavirusCanal-1]/149-304 | - | D81 | L64 | G47 | F6 | G48 | I84 | F1 | - | - |
| XOK28216.1/1-580putativebifunctionalchitinase/lysozyme\[NES-4A-S1]/149-304 | - | D81 | L64 | G47 | F6 | G48 | I84 | F1 | - | - |
| XOK30852.1/1-582putativebifunctionalchitinase/lysozyme\[S-NE-9]/153-304 | - | D77 | L60 | G43 | F2 | G44 | I80 | - | - | - |
| XOK34326.1/1-580putativebifunctionalchitinase/lysozyme\[S-NE-18]/152-304 | - | D78 | L61 | G44 | F3 | G45 | I81 | - | - | - |
| AGE56814.1/1-582putativebifunctionalchitinase/lysozyme\[AcanthocystisturfaceaChlorellavirusNE-JV-2]/153-304 | - | D77 | L60 | G43 | F2 | G44 | I80 | - | - | - |
| AGE56814.1/1-582putativebifunctionalchitinase/lysozyme\[AcanthocystisturfaceaChlorellavirusNE-JV-2]/153-304 | - | D77 | L60 | G43 | F2 | G44 | I80 | - | - | - |
| XOK32790.1/1-582putativebifunctionalchitinase/lysozyme\[S-NE-15]/153-304 | - | D77 | L60 | G43 | F2 | G44 | I80 | - | - | - |
| XOK35891.1/1-582putativebifunctionalchitinase/lysozyme\[S-NE-23]/153-304 | - | D77 | L60 | G43 | F2 | G44 | I80 | - | - | - |
| XOK33538.1/1-582putativebifunctionalchitinase/lysozyme\[S-NE-16]/153-304 | - | D77 | L60 | G43 | F2 | G44 | I80 | - | - | - |
| XOK33601.1/1-582putativebifunctionalchitinase/lysozyme\[S-NE-17]/153-304 | - | D77 | L60 | G43 | F2 | G44 | I80 | - | - | - |
| XOK32004.1/1-582putativebifunctionalchitinase/lysozyme\[S-NE-12]/153-304 | - | D77 | L60 | G43 | F2 | G44 | I80 | - | - | - |
| XOK35119.1/1-582putativebifunctionalchitinase/lysozyme\[S-NE-20]/175-304 | - | D55 | L38 | G21 | - | G22 | I58 | - | - | - |
| XOK27846.1/1-580putativebifunctionalchitinase/lysozyme\[NES-4A-M1]/152-304 | - | D78 | L61 | G44 | F3 | G45 | I81 | - | - | - |
| XOK30419.1/1-582chitinase\[S-NE-7]/153-304 | - | D77 | L60 | G43 | F2 | G44 | I80 | - | - | - |
| AGE59594.1/1-582putativebifunctionalchitinase/lysozyme\[AcanthocystisturfaceaChlorellavirusOR0704.3]/153-304 | - | D77 | L60 | G43 | F2 | G44 | I80 | - | - | - |
| XOK32390.1/1-589putativebifunctionalchitinase/lysozyme\[S-NE-13]/153-304 | - | D77 | L60 | G43 | F2 | G44 | I80 | - | - | - |
| AGE57849.1/1-601putativebifunctionalchitinase/lysozyme\[AcanthocystisturfaceaChlorellavirusNTS-1]/172-323 | - | D77 | L60 | G43 | F2 | G44 | I80 | - | - | - |

**Table S4:** Prevalence of coevolving residues on chlorovirus chitinases in set 2. Colored in green represents coevolved residues.

| Sequence | D560 | D901 | E563 | G534 | M895 | G1158 | G1175 | Y1169 | W1750 | G1741 | Y899 |
|---|---|---|---|---|---|---|---|---|---|---|---|
| WZO81342.1/1-840bifunctionalchitinase/lysozyme\[CA-4B]/142-304 | D90 | D159 | E92 | R86 | M156 | - | - | - | - | - | M158 |
| WZO81342.1/1-840bifunctionalchitinase/lysozyme\[CA-4B]/573-747 | D99 | D168 | E101 | N95 | M165 | - | - | - | - | - | M167 |
| WZO81006.1/1-840putativebifunctionalchitinase/lysozyme\[CA-4A]/142-304 | D90 | D159 | E92 | R86 | M156 | - | - | - | - | - | M158 |
| WZO81006.1/1-840putativebifunctionalchitinase/lysozyme\[CA-4A]/573-747 | D99 | D168 | E101 | N95 | M165 | - | - | - | - | - | M167 |
| WZO82266.1/1-840putativebifunctionalchitinase/lysozyme\[NY-2C]/142-304 | D90 | D159 | E92 | R86 | M156 | - | - | - | - | - | M158 |
| WZO82266.1/1-840putativebifunctionalchitinase/lysozyme\[NY-2C]/573-747 | D99 | D168 | E101 | N95 | M165 | - | - | - | - | - | M167 |
| WZO80816.1/1-830putativebifunctionalchitinase/lysozyme\[WNE-11A-L2]/142-304 | D90 | D159 | E92 | R86 | M156 | - | - | - | - | - | M158 |
| WZO80816.1/1-830putativebifunctionalchitinase/lysozyme\[WNE-11A-L2]/563-740 | D99 | D168 | E101 | N95 | M165 | - | - | - | - | - | M167 |
| AGE57241.1/1-815putativebifunctionalchitinase/lysozyme\[ParameciumbursariaChlorellavirusNE-JV-4]/142-304 | D90 | D159 | E92 | R86 | M156 | - | - | - | - | - | M158 |
| AGE57241.1/1-815putativebifunctionalchitinase/lysozyme\[ParameciumbursariaChlorellavirusNE-JV-4]/548-725 | D99 | D168 | E101 | N95 | M165 | - | - | - | - | - | M167 |
| WZO75107.1/1-840chitinase\[40-NE-3]/142-304 | D90 | D159 | E92 | R86 | M156 | - | - | - | - | - | M158 |
| WZO75107.1/1-840chitinase\[40-NE-3]/573-747 | D99 | D168 | E101 | N95 | M165 | - | - | - | - | - | M167 |
| WZO75726.1/1-840putativebifunctionalchitinase/lysozyme\[40-NE-4]/142-304 | D90 | D159 | E92 | R86 | M156 | - | - | - | - | - | M158 |
| WZO75726.1/1-840putativebifunctionalchitinase/lysozyme\[40-NE-4]/573-747 | D99 | D168 | E101 | N95 | M165 | - | - | - | - | - | M167 |
| WZO76741.1/1-840chitinase\[41-NE-5]/142-304 | D90 | D159 | E92 | R86 | M156 | - | - | - | - | - | M158 |
| WZO76741.1/1-840chitinase\[41-NE-5]/573-747 | D99 | D168 | E101 | N95 | M165 | - | - | - | - | - | M167 |
| WZO83158.1/1-830chitinase\[XZ-4C]/142-304 | D90 | D159 | E92 | R86 | M156 | - | - | - | - | - | M158 |
| WZO83158.1/1-830chitinase\[XZ-4C]/563-737 | D99 | D168 | E101 | N95 | M165 | - | - | - | - | - | M167 |
| AGE53811.1/1-835putativebifunctionalchitinase/lysozyme\[ParameciumbursariaChlorellavirusIL3A]/142-304 | D90 | D159 | E92 | R86 | M156 | - | - | - | - | - | M158 |
| AGE53811.1/1-835putativebifunctionalchitinase/lysozyme\[ParameciumbursariaChlorellavirusIL3A]/567-745 | D100 | D169 | E102 | N96 | M166 | - | - | - | - | - | M168 |
| AGE54506.1/1-817putativebifunctionalchitinase/lysozyme\[ParameciumbursariaChlorellavirusKS1B]/142-304 | D90 | D159 | E92 | R86 | M156 | - | - | - | - | - | M158 |
| AGE54506.1/1-817putativebifunctionalchitinase/lysozyme\[ParameciumbursariaChlorellavirusKS1B]/550-727 | D99 | D168 | E101 | N95 | M165 | - | - | - | - | - | M167 |
| AGE48399.1/1-835putativebifunctionalchitinase/lysozyme\[ParameciumbursariaChlorellavirusAN69C]/142-304 | D90 | D159 | E92 | R86 | M156 | - | - | - | - | - | M158 |
| AGE48399.1/1-835putativebifunctionalchitinase/lysozyme\[ParameciumbursariaChlorellavirusAN69C]/568-742 | D99 | D168 | E101 | N95 | M165 | - | - | - | - | - | M167 |
| WZO91574.1/1-809putativebifunctionalchitinase/lysozyme\[O-NE-29]/139-304 | D93 | D162 | E95 | R89 | M159 | - | - | - | - | - | M161 |
| WZO91574.1/1-809putativebifunctionalchitinase/lysozyme\[O-NE-29]/542-716 | D99 | D168 | E101 | H95 | M165 | - | - | - | - | - | M167 |

| | | | | | | | | | | | |
|---|---|---|---|---|---|---|---|---|---|---|---|
| WZO81686.1/1-840chitinase\[NC-1A]/142-304 | D90 | D159 | E92 | R86 | M156 | - | - | - | - | - | M158 |
| WZO81686.1/1-840chitinase\[NC-1A]/573-747 | D99 | D168 | E101 | N95 | M165 | - | - | - | - | - | M167 |
| NP_048529.2/1-830Chitinase\[ParameciumbursariaChlorellavirus1]/142-304 | D90 | D159 | E92 | R86 | M156 | - | - | - | - | - | M158 |
| NP_048529.2/1-830Chitinase\[ParameciumbursariaChlorellavirus1]/563-737 | D99 | D168 | E101 | N95 | M165 | - | - | - | - | - | M167 |
| WZO82806.1/1-830chitinase\[XZ-3A]/142-304 | D90 | D159 | E92 | R86 | M156 | - | - | - | - | - | M158 |
| WZO82806.1/1-830chitinase\[XZ-3A]/563-737 | D99 | D168 | E101 | N95 | M165 | - | - | - | - | - | M167 |
| WZO90762.1/1-809putativebifunctionalchitinase/lysozyme\[O-NE-27]/139-304 | D93 | D162 | E95 | R89 | M159 | - | - | - | - | - | M161 |
| WZO90762.1/1-809putativebifunctionalchitinase/lysozyme\[O-NE-27]/542-716 | D99 | D168 | E101 | H95 | M165 | - | - | - | - | - | M167 |
| WZO83510.1/1-820putativebifunctionalchitinase/lysozyme\[XZ-5C]/142-304 | D90 | D159 | E92 | R86 | M156 | - | - | - | - | - | M158 |
| WZO83510.1/1-820putativebifunctionalchitinase/lysozyme\[XZ-5C]/553-731 | D99 | D168 | E101 | N95 | M165 | - | - | - | - | - | M167 |
| WZO89952.1/1-822putativebifunctionalchitinase/lysozyme\[O-NE-25]/142-304 | D90 | D159 | E92 | R86 | M156 | - | - | - | - | - | M158 |
| WZO89952.1/1-822putativebifunctionalchitinase/lysozyme\[O-NE-25]/555-729 | D99 | D168 | E101 | H95 | M165 | - | - | - | - | - | M167 |
| AGE51440.1/1-836putativebifunctionalchitinase/lysozyme\[ParameciumbursariaChlorellavirusCviKI]/142-304 | D90 | D159 | E92 | R86 | M156 | - | - | - | - | - | M158 |
| AGE51440.1/1-836putativebifunctionalchitinase/lysozyme\[ParameciumbursariaChlorellavirusCviKI]/569-748 | D99 | D168 | E101 | N95 | M165 | - | - | - | - | - | M167 |
| AGE52455.1/1-826putativebifunctionalchitinase/lysozyme\[ParameciumbursariaChlorellavirusCvsA1]/142-304 | D90 | D159 | E92 | R86 | M156 | - | - | - | - | - | M158 |
| AGE52455.1/1-826putativebifunctionalchitinase/lysozyme\[ParameciumbursariaChlorellavirusCvsA1]/559-739 | D99 | D168 | E101 | N95 | M165 | - | - | - | - | - | M167 |
| WZO87987.1/1-821putativebifunctionalchitinase/lysozyme\[O-NE-19]/142-304 | D90 | D159 | E92 | H86 | M156 | - | - | - | - | - | M158 |
| WZO87987.1/1-821putativebifunctionalchitinase/lysozyme\[O-NE-19]/554-728 | D99 | D168 | E101 | H95 | M165 | - | - | - | - | - | M167 |
| WZO82436.1/1-820putativebifunctionalchitinase/lysozyme\[SH-6A]/142-304 | D90 | D159 | E92 | R86 | M156 | - | - | - | - | - | M158 |
| WZO82436.1/1-820putativebifunctionalchitinase/lysozyme\[SH-6A]/553-731 | D99 | D168 | E101 | N95 | M165 | - | - | - | - | - | M167 |
| WZO92132.1/1-827putativebifunctionalchitinase/lysozyme\[NE-O-7-s]/139-304 | D93 | D162 | E95 | R89 | M159 | - | - | - | - | - | M161 |
| WZO92132.1/1-827putativebifunctionalchitinase/lysozyme\[NE-O-7-s]/560-734 | D99 | D168 | E101 | H95 | M165 | - | - | - | - | - | M167 |
| WZO89127.1/1-809putativebifunctionalchitinase/lysozyme\[O-NE-23]/139-304 | D93 | D162 | E95 | R89 | M159 | - | - | - | - | - | M161 |
| WZO89127.1/1-809putativebifunctionalchitinase/lysozyme\[O-NE-23]/542-716 | D99 | D168 | E101 | H95 | M165 | - | - | - | - | - | M167 |
| WZO85876.1/1-819putativebifunctionalchitinase/lysozyme\[O-NE-14]/142-304 | D90 | D159 | E92 | R86 | M156 | - | - | - | - | - | M158 |
| WZO85876.1/1-819putativebifunctionalchitinase/lysozyme\[O-NE-14]/552-726 | D99 | D168 | E101 | N95 | M165 | - | - | - | - | - | M167 |
| WZO88848.1/1-822putativebifunctionalchitinase/lysozyme\[O-NE-22]/142-304 | D90 | D159 | E92 | H86 | M156 | - | - | - | - | - | M158 |
| WZO88848.1/1-822putativebifunctionalchitinase/lysozyme\[O-NE-22]/555-729 | D99 | D168 | E101 | H95 | M165 | - | - | - | - | - | M167 |
| WZO83897.1/1-820putativebifunctionalchitinase/lysozyme\[XZ-6E]/142-304 | D90 | D159 | E92 | R86 | M156 | - | - | - | - | - | M158 |
| WZO83897.1/1-820putativebifunctionalchitinase/lysozyme\[XZ-6E]/553-732 | D99 | D168 | E101 | N95 | M165 | - | - | - | - | - | M167 |
| YP_009665302.1/1-806putativebifunctionalchitinase/lysozyme\[ParameciumbursariaChlorellavirusNYs1]/142-304 | D90 | D159 | E92 | R86 | M156 | - | - | - | - | - | M158 |
| YP_009665302.1/1-806putativebifunctionalchitinase/lysozyme\[ParameciumbursariaChlorellavirusNYs1]/539-715 | D99 | D168 | E101 | Y95 | M165 | - | - | - | - | - | M167 |
| WZO87550.1/1-808putativebifunctionalchitinase/lysozyme\[O-NE-18]/143-303 | D88 | D157 | E90 | R84 | M154 | - | - | - | - | - | M156 |
| WZO87550.1/1-808putativebifunctionalchitinase/lysozyme\[O-NE-18]/541-715 | D99 | D168 | E101 | H95 | M165 | - | - | - | - | - | M167 |
| AGE54806.1/1-806putativebifunctionalchitinase/lysozyme\[ParameciumbursariaChlorellavirusMA1D]/142-304 | D90 | D159 | E92 | R86 | M156 | - | - | - | - | - | M158 |
| AGE54806.1/1-806putativebifunctionalchitinase/lysozyme\[ParameciumbursariaChlorellavirusMA1D]/539-715 | D99 | D168 | E101 | Y95 | M165 | - | - | - | - | - | M167 |
| WZO84706.1/1-831putativebifunctionalchitinase/lysozyme\[O-NE-11]/142-304 | D90 | D159 | E92 | R86 | M156 | - | - | - | - | - | M158 |
| WZO84706.1/1-831putativebifunctionalchitinase/lysozyme\[O-NE-11]/564-738 | D99 | D168 | E101 | N95 | M165 | - | - | - | - | - | M167 |
| WZO80449.1/1-806putativebifunctionalchitinase/lysozyme\[WNE-10B-S1]/142-304 | D90 | D159 | E92 | R86 | M156 | - | - | - | - | - | M158 |
| WZO80449.1/1-806putativebifunctionalchitinase/lysozyme\[WNE-10B-S1]/539-714 | D99 | D168 | E101 | Y95 | M165 | - | - | - | - | - | M167 |
| WZO85478.1/1-830putativebifunctionalchitinase/lysozyme\[O-NE-13]/142-304 | D90 | D159 | E92 | R86 | M156 | - | - | - | - | - | M158 |
| WZO85478.1/1-830putativebifunctionalchitinase/lysozyme\[O-NE-13]/564-737 | D98 | D167 | E100 | N94 | M164 | - | - | - | - | - | M166 |
| WZO76146.1/1-816bifunctionalchitinase/lysozyme\[40-NE-5]/142-304 | D90 | D159 | E92 | R86 | M156 | - | - | - | - | - | M158 |
| WZO76146.1/1-816bifunctionalchitinase/lysozyme\[40-NE-5]/549-723 | D99 | D168 | E101 | N95 | M165 | - | - | - | - | - | M167 |
| WZO76561.1/1-816putativebifunctionalchitinase/lysozyme\[41-NE-4]/142-304 | D90 | D159 | E92 | R86 | M156 | - | - | - | - | - | M158 |
| WZO76561.1/1-816putativebifunctionalchitinase/lysozyme\[41-NE-4]/549-723 | D99 | D168 | E101 | N95 | M165 | - | - | - | - | - | M167 |
| WZO78129.1/1-816chitinase\[NE-40â??2-s]/142-304 | D90 | D159 | E92 | R86 | M156 | - | - | - | - | - | M158 |
| WZO78129.1/1-816chitinase\[NE-40â??2-s]/549-723 | D99 | D168 | E101 | N95 | M165 | - | - | - | - | - | M167 |
| WZO79003.1/1-816putativebifunctionalchitinase/lysozyme\[NE-41-3-s]/142-304 | D90 | D159 | E92 | R86 | M156 | - | - | - | - | - | M158 |
| WZO79003.1/1-816putativebifunctionalchitinase/lysozyme\[NE-41-3-s]/549-723 | D99 | D168 | E101 | N95 | M165 | - | - | - | - | - | M167 |
| WZO80062.1/1-823putativebifunctionalchitinase/lysozyme\[N-NE-5]/142-304 | D90 | D159 | E92 | R86 | M156 | - | - | - | - | - | M158 |
| WZO80062.1/1-823putativebifunctionalchitinase/lysozyme\[N-NE-5]/549-723 | D99 | D168 | E101 | N95 | M165 | - | - | - | - | - | M167 |
| AGE58174.1/1-816putativebifunctionalchitinase/lysozyme\[ParameciumbursariaChlorellavirusNW665.2]/140-305 | D93 | D162 | E95 | R89 | M159 | - | - | - | - | - | M161 |
| AGE58174.1/1-816putativebifunctionalchitinase/lysozyme\[ParameciumbursariaChlorellavirusNW665.2]/550-723 | D98 | D167 | E100 | Y94 | M164 | - | - | - | - | - | M166 |
| YP_001426411.1/1-805hypotheticalproteinFR483_N779R\[ParameciumbursariaChlorellavirusFR483]/139-304 | D93 | D162 | E95 | R89 | M159 | - | - | - | - | - | M161 |
| YP_001426411.1/1-805hypotheticalproteinFR483_N779R\[ParameciumbursariaChlorellavirusFR483]/539-712 | D98 | D167 | E100 | Y94 | M164 | - | - | - | - | - | M166 |
| WZO86764.1/1-525chitinase\[O-NE-16]/142-304 | D90 | D159 | E92 | R86 | M156 | - | - | - | - | - | M158 |
| WZO93853.1/1-525chitinase\[OSyNE-4B-S1]/142-304 | D90 | D159 | E92 | R86 | M156 | - | - | - | - | - | M158 |
| WZO91298.1/1-445chitinase\[O-NE-28]/147-309 | D90 | D159 | E92 | R86 | M156 | - | - | - | - | - | M158 |
| WZO89577.1/1-445chitinase\[O-NE-24]/147-309 | D90 | D159 | E92 | R86 | M156 | - | - | - | - | - | M158 |
| WZO90501.1/1-525chitinase\[O-NE-26]/142-304 | D90 | D159 | E92 | R86 | M156 | - | - | - | - | - | M158 |
| WZO77210.1/1-525chitinase\[41-NE-6]/142-304 | D90 | D159 | E92 | R86 | M156 | - | - | - | - | - | M158 |
| WZO77693.1/1-525chitinase\[NE-40-1-m]/142-304 | D90 | D159 | E92 | R86 | M156 | - | - | - | - | - | M158 |
| WZO78681.1/1-525chitinase\[NE-41-2-m]/142-304 | D90 | D159 | E92 | R86 | M156 | - | - | - | - | - | M158 |
| WZO85099.1/1-530chitinase\[O-NE-12]/147-309 | D90 | D159 | E92 | R86 | M156 | - | - | - | - | - | M158 |
| YP_009325635.1/1-525Chitinase\[OnlySyngenNebraskavirus5]/142-304 | D90 | D159 | E92 | R86 | M156 | - | - | - | - | - | M158 |
| WZO88270.1/1-528chitinase\[O-NE-20]/147-309 | D90 | D159 | E92 | R86 | M156 | - | - | - | - | - | M158 |
| WZO92747.1/1-528chitinase\[NE-O-9-L]/147-309 | D90 | D159 | E92 | R86 | M156 | - | - | - | - | - | M158 |
| WZO93306.1/1-528chitinase\[OSyNE-4B-L2]/147-309 | D90 | D159 | E92 | R86 | M156 | - | - | - | - | - | M158 |
| WZO87136.1/1-528chitinase\[O-NE-17]/147-309 | D90 | D159 | E92 | R86 | M156 | - | - | - | - | - | M158 |
| WZO92491.1/1-528chitinase\[NE-O-8-L]/147-309 | D90 | D159 | E92 | R86 | M156 | - | - | - | - | - | M158 |

| | | | | | | | | | | | |
|---|---|---|---|---|---|---|---|---|---|---|---|
| WZO95440.1/1-528chitinase\[OSyNE-ZA]/147-309 | D90 | D159 | E92 | R86 | M156 | - | - | - | - | - | M158 |
| WZO78440.1/1-532chitinase\[NE-41-1-L]/147-309 | D90 | D159 | E92 | R86 | M156 | - | - | - | - | - | M158 |
| WZO94180.1/1-539chitinase\[OSyNE-5A-L1]/147-309 | D90 | D159 | E92 | R86 | M156 | - | - | - | - | - | M158 |
| WZO84329.1/1-525chitinase\[O-NE-10]/142-304 | D90 | D159 | E92 | R86 | M156 | - | - | - | - | - | M158 |
| WZO94943.1/1-525chitinase\[OSyNE-5B-S1]/142-304 | D90 | D159 | E92 | R86 | M156 | - | - | - | - | - | M158 |
| WZO93480.1/1-525chitinase\[OSyNE-4B-M2]/142-304 | D90 | D159 | E92 | R86 | M156 | - | - | - | - | - | M158 |
| WZO94540.1/1-525chitinase\[OSyNE-5B-M2]/142-304 | D90 | D159 | E92 | R86 | M156 | - | - | - | - | - | M158 |
| WZO79465.1/1-530chitinase\[N-NE-4]/147-309 | D90 | D159 | E92 | R86 | M156 | - | - | - | - | - | M158 |
| XYW60054.1/1-521chitinase\[NE-P-3-s]/139-304 | D93 | D162 | E95 | R89 | M159 | - | - | - | - | - | M161 |
| XYW60732.1/1-521chitinase\[NE-P-4-fs]/139-304 | D93 | D162 | E95 | R89 | M159 | - | - | - | - | - | M161 |
| XYW61232.1/1-521chitinase\[NE-P-8-f]/139-304 | D93 | D162 | E95 | R89 | M159 | - | - | - | - | - | M161 |
| XYW62772.1/1-521chitinase\[P-NE-12]/139-304 | D93 | D162 | E95 | R89 | M159 | - | - | - | - | - | M161 |
| WZO86398.1/1-532chitinase\[O-NE-15]/144-309 | D93 | D162 | E95 | R89 | M159 | - | - | - | - | - | M161 |
| XOK27477.1/1-582putativebifunctionalchitinase/lysozyme\[CL-S-1-m]/153-304 | D79 | D148 | E81 | R75 | M145 | - | - | - | - | - | M147 |
| AGE49332.1/1-595putativebifunctionalchitinase/lysozyme\[AcanthocystisturfaceaChlorellavirusBr0604L]/166-317 | D79 | D148 | E81 | R75 | M145 | - | - | - | - | - | M147 |
| AGE48989.1/1-520putativebifunctionalchitinase/lysozyme\[ParameciumbursariaChlorellavirusAP110A]/139-304 | D93 | D162 | E95 | R89 | M159 | - | - | - | - | - | M161 |
| AGE50008.1/1-520putativebifunctionalchitinase/lysozyme\[ParameciumbursariaChlorellavirusCan18-4]/139-304 | D93 | D162 | E95 | R89 | M159 | - | - | - | - | - | M161 |
| XYW62298.1/1-520chitinase\[P-NE-10]/139-304 | D93 | D162 | E95 | R89 | M159 | - | - | - | - | - | M161 |
| AGE52349.1/1-523putativebifunctionalchitinase/lysozyme\[ParameciumbursariaChlorellavirusCVR-1]/143-304 | D89 | D158 | E91 | R85 | M155 | - | - | - | - | - | M157 |
| YP_009702007.1/1-523putativebifunctionalchitinase/lysozyme\[ParameciumbursariaChlorellavirusCVA-1]/143-304 | D89 | D158 | E91 | R85 | M155 | - | - | - | - | - | M157 |
| ABT14345.1/1-520hypotheticalproteinMT325_M791R\[ParameciumbursariachlorellavirusMT325]/139-304 | D93 | D162 | E95 | R89 | M159 | - | - | - | - | - | M161 |
| AGE51335.1/1-523putativebifunctionalchitinase/lysozyme\[ParameciumbursariaChlorellavirusCVG-1]/139-304 | D93 | D162 | E95 | R89 | M159 | - | - | - | - | - | M161 |
| XYW61920.1/1-523chitinase\[P-NE-9]/139-304 | D93 | D162 | E95 | R89 | M159 | - | - | - | - | - | M161 |
| XOK31597.1/1-580putativebifunctionalchitinase/lysozyme\[S-NE-11]/151-304 | D81 | D150 | E83 | R77 | M147 | - | - | - | - | - | M149 |
| XOK36306.1/1-580putativebifunctionalchitinase/lysozyme\[GNLD22]/152-304 | D80 | D149 | E82 | R76 | M146 | - | - | - | - | - | M148 |
| XOK36264.1/1-582putativebifunctionalchitinase/lysozyme\[LP-F3a-4a]/153-304 | D79 | D148 | E81 | R75 | M145 | - | - | - | - | - | M147 |
| XOK31528.1/1-582putativebifunctionalchitinase/lysozyme\[S-NE-10]/153-304 | D79 | D148 | E81 | R75 | M145 | - | - | - | - | - | M147 |
| XOK28546.1/1-582putativebifunctionalchitinase/lysozyme\[NES-4B-L1]/153-304 | D79 | D148 | E81 | R75 | M145 | - | - | - | - | - | M147 |
| YP_001427295.1/1-595hypotheticalproteinATCV1_Z814L\[Acanthocystisturfaceachlorellavirus1]/166-317 | D79 | D148 | E81 | R75 | M145 | - | - | - | - | - | M147 |
| AGE56132.1/1-601putativebifunctionalchitinase/lysozyme\[AcanthocystisturfaceaChlorellavirusMO0605SPH]/172-323 | D79 | D148 | E81 | R75 | M145 | - | - | - | - | - | M147 |
| XOK34753.1/1-421putativebifunctionalchitinase/lysozyme\[S-NE-19]/153-304 | D79 | D148 | E81 | R75 | M145 | - | - | - | - | - | M147 |
| AGE60252.1/1-601putativebifunctionalchitinase/lysozyme\[AcanthocystisturfaceaChlorellavirusWI0606]/172-323 | D79 | D148 | E81 | R75 | M145 | - | - | - | - | - | M147 |
| AGE57134.1/1-601putativebifunctionalchitinase/lysozyme\[AcanthocystisturfaceaChlorellavirusNE-JV-3]/172-323 | D79 | D148 | E81 | R75 | M145 | - | - | - | - | - | M147 |
| XOK29995.1/1-582putativebifunctionalchitinase/lysozyme\[NES-5A-S1]/153-304 | D79 | D148 | E81 | R75 | M145 | - | - | - | - | - | M147 |
| AGE49664.1/1-595putativebifunctionalchitinase/lysozyme\[AcanthocystisturfaceaChlorellavirusCan0610SP]/166-317 | D79 | D148 | E81 | R75 | M145 | - | - | - | - | - | M147 |
| AGE59935.1/1-595putativebifunctionalchitinase/lysozyme\[AcanthocystisturfaceaChlorellavirusTN603.4.2]/189-317 | D56 | D125 | E58 | R52 | M122 | - | - | - | - | - | M124 |
| AGE55823.1/1-596putativebifunctionalchitinase/lysozyme\[AcanthocystisturfaceaChlorellavirusMN0810.1]/185-317 | D60 | D129 | E62 | R56 | M126 | - | - | - | - | - | M128 |
| XOK35208.1/1-580putativebifunctionalchitinase/lysozyme\[S-NE-22]/149-304 | D83 | D152 | E85 | R79 | M149 | - | - | - | - | - | M151 |
| AGE50333.1/1-580putativebifunctionalchitinase/lysozyme\[AcanthocystisturfaceaChlorellavirusCanal-1]/149-304 | D83 | D152 | E85 | R79 | M149 | - | - | - | - | - | M151 |
| XOK28216.1/1-580putativebifunctionalchitinase/lysozyme\[NES-4A-S1]/149-304 | D83 | D152 | E85 | R79 | M149 | - | - | - | - | - | M151 |
| XOK30852.1/1-582putativebifunctionalchitinase/lysozyme\[S-NE-9]/153-304 | D79 | D148 | E81 | R75 | M145 | - | - | - | - | - | M147 |
| XOK34326.1/1-580putativebifunctionalchitinase/lysozyme\[S-NE-18]/152-304 | D80 | D149 | E82 | R76 | M146 | - | - | - | - | - | M148 |
| AGE56814.1/1-582putativebifunctionalchitinase/lysozyme\[AcanthocystisturfaceaChlorellavirusNE-JV-2]/153-304 | D79 | D148 | E81 | R75 | M145 | - | - | - | - | - | M147 |
| AGE56814.1/1-582putativebifunctionalchitinase/lysozyme\[AcanthocystisturfaceaChlorellavirusNE-JV-2]/153-304 | D79 | D148 | E81 | R75 | M145 | - | - | - | - | - | M147 |
| XOK32790.1/1-582putativebifunctionalchitinase/lysozyme\[S-NE-15]/153-304 | D79 | D148 | E81 | R75 | M145 | - | - | - | - | - | M147 |
| XOK35891.1/1-582putativebifunctionalchitinase/lysozyme\[S-NE-23]/153-304 | D79 | D148 | E81 | R75 | M145 | - | - | - | - | - | M147 |
| XOK33538.1/1-582putativebifunctionalchitinase/lysozyme\[S-NE-16]/153-304 | D79 | D148 | E81 | R75 | M145 | - | - | - | - | - | M147 |
| XOK33601.1/1-582putativebifunctionalchitinase/lysozyme\[S-NE-17]/153-304 | D79 | D148 | E81 | R75 | M145 | - | - | - | - | - | M147 |
| XOK32004.1/1-582putativebifunctionalchitinase/lysozyme\[S-NE-12]/153-304 | D79 | D148 | E81 | R75 | M145 | - | - | - | - | - | M147 |
| XOK35119.1/1-582putativebifunctionalchitinase/lysozyme\[S-NE-20]/175-304 | D57 | D126 | E59 | R53 | M123 | - | - | - | - | - | M125 |
| XOK27846.1/1-580putativebifunctionalchitinase/lysozyme\[NES-4A-M1]/152-304 | D80 | D149 | E82 | R76 | M146 | - | - | - | - | - | M148 |
| XOK30419.1/1-582chitinase\[S-NE-7]/153-304 | D79 | D148 | E81 | R75 | M145 | - | - | - | - | - | M147 |
| AGE59594.1/1-582putativebifunctionalchitinase/lysozyme\[AcanthocystisturfaceaChlorellavirusOR0704.3]/153-304 | D79 | D148 | E81 | R75 | M145 | - | - | - | - | - | M147 |
| XOK32390.1/1-589putativebifunctionalchitinase/lysozyme\[S-NE-13]/153-304 | D79 | D148 | E81 | R75 | M145 | - | - | - | - | - | M147 |
| AGE57849.1/1-601putativebifunctionalchitinase/lysozyme\[AcanthocystisturfaceaChlorellavirusNTS-1]/172-323 | D79 | D148 | E81 | R75 | M145 | - | - | - | - | - | M147 |

**Table S5:** Prevalence of coevolving residues on chlorovirus chitinases in set 3.

| Sequence | C1406 | C71 | C4 |
|---|---|---|---|
| WZO81342.1/1-840bifunctionalchitinase/lysozyme\[CA-4B]/142-304 | - | L6 | - |
| WZO81342.1/1-840bifunctionalchitinase/lysozyme\[CA-4B]/573-747 | - | V21 | Q2 |
| WZO81006.1/1-840putativebifunctionalchitinase/lysozyme\[CA-4A]/142-304 | - | L6 | - |
| WZO81006.1/1-840putativebifunctionalchitinase/lysozyme\[CA-4A]/573-747 | - | V21 | Q2 |
| WZO82266.1/1-840putativebifunctionalchitinase/lysozyme\[NY-2C]/142-304 | - | L6 | - |
| WZO82266.1/1-840putativebifunctionalchitinase/lysozyme\[NY-2C]/573-747 | - | V21 | Q2 |
| WZO80816.1/1-830putativebifunctionalchitinase/lysozyme\[WNE-11A-L2]/142-304 | - | L6 | - |
| WZO80816.1/1-830putativebifunctionalchitinase/lysozyme\[WNE-11A-L2]/563-740 | - | V21 | Q2 |

| | | | |
|---|---|---|---|
| **AGE57241.1/1-815putativebifunctionalchitinase/lysozyme\[ParameciumbursariaChlorellavirusNE-JV-4]/142-304** | - | L6 | - |
| **AGE57241.1/1-815putativebifunctionalchitinase/lysozyme\[ParameciumbursariaChlorellavirusNE-JV-4]/548-725** | - | V21 | Q2 |
| **WZO75107.1/1-840chitinase\[40-NE-3]/142-304** | - | L6 | - |
| **WZO75107.1/1-840chitinase\[40-NE-3]/573-747** | - | V21 | Q2 |
| **WZO75726.1/1-840putativebifunctionalchitinase/lysozyme\[40-NE-4]/142-304** | - | L6 | - |
| **WZO75726.1/1-840putativebifunctionalchitinase/lysozyme\[40-NE-4]/573-747** | - | V21 | Q2 |
| **WZO76741.1/1-840chitinase\[41-NE-5]/142-304** | - | L6 | - |
| **WZO76741.1/1-840chitinase\[41-NE-5]/573-747** | - | V21 | Q2 |
| **WZO83158.1/1-830chitinase\[XZ-4C]/142-304** | - | L6 | - |
| **WZO83158.1/1-830chitinase\[XZ-4C]/563-737** | - | V21 | Q2 |
| **AGE53811.1/1-835putativebifunctionalchitinase/lysozyme\[ParameciumbursariaChlorellavirusIL3A]/142-304** | - | L6 | - |
| **AGE53811.1/1-835putativebifunctionalchitinase/lysozyme\[ParameciumbursariaChlorellavirusIL3A]/567-745** | - | V22 | Q3 |
| **AGE54506.1/1-817putativebifunctionalchitinase/lysozyme\[ParameciumbursariaChlorellavirusKS1B]/142-304** | - | L6 | - |
| **AGE54506.1/1-817putativebifunctionalchitinase/lysozyme\[ParameciumbursariaChlorellavirusKS1B]/550-727** | - | V21 | Q2 |
| **AGE48399.1/1-835putativebifunctionalchitinase/lysozyme\[ParameciumbursariaChlorellavirusAN69C]/142-304** | - | L6 | - |
| **AGE48399.1/1-835putativebifunctionalchitinase/lysozyme\[ParameciumbursariaChlorellavirusAN69C]/568-742** | - | V21 | Q2 |
| **WZO91574.1/1-809putativebifunctionalchitinase/lysozyme\[O-NE-29]/139-304** | - | L9 | - |
| **WZO91574.1/1-809putativebifunctionalchitinase/lysozyme\[O-NE-29]/542-716** | - | V21 | N2 |
| **WZO81686.1/1-840chitinase\[NC-1A]/142-304** | - | L6 | - |
| **WZO81686.1/1-840chitinase\[NC-1A]/573-747** | - | V21 | Q2 |
| **NP_048529.2/1-830Chitinase\[ParameciumbursariaChlorellavirus1]/142-304** | - | L6 | - |
| **NP_048529.2/1-830Chitinase\[ParameciumbursariaChlorellavirus1]/563-737** | - | V21 | Q2 |
| **WZO82806.1/1-830chitinase\[XZ-3A]/142-304** | - | L6 | - |
| **WZO82806.1/1-830chitinase\[XZ-3A]/563-737** | - | V21 | Q2 |
| **WZO90762.1/1-809putativebifunctionalchitinase/lysozyme\[O-NE-27]/139-304** | - | L9 | - |
| **WZO90762.1/1-809putativebifunctionalchitinase/lysozyme\[O-NE-27]/542-716** | - | V21 | N2 |
| **WZO83510.1/1-820putativebifunctionalchitinase/lysozyme\[XZ-5C]/142-304** | - | L6 | - |
| **WZO83510.1/1-820putativebifunctionalchitinase/lysozyme\[XZ-5C]/553-731** | - | V21 | Q2 |
| **WZO89952.1/1-822putativebifunctionalchitinase/lysozyme\[O-NE-25]/142-304** | - | L6 | - |
| **WZO89952.1/1-822putativebifunctionalchitinase/lysozyme\[O-NE-25]/555-729** | - | V21 | N2 |
| **AGE51440.1/1-836putativebifunctionalchitinase/lysozyme\[ParameciumbursariaChlorellavirusCviKI]/142-304** | - | L6 | - |
| **AGE51440.1/1-836putativebifunctionalchitinase/lysozyme\[ParameciumbursariaChlorellavirusCviKI]/569-748** | - | V21 | S2 |
| **AGE52455.1/1-826putativebifunctionalchitinase/lysozyme\[ParameciumbursariaChlorellavirusCvsA1]/142-304** | - | L6 | - |
| **AGE52455.1/1-826putativebifunctionalchitinase/lysozyme\[ParameciumbursariaChlorellavirusCvsA1]/559-739** | - | V21 | S2 |
| **WZO87987.1/1-821putativebifunctionalchitinase/lysozyme\[O-NE-19]/142-304** | - | L6 | - |
| **WZO87987.1/1-821putativebifunctionalchitinase/lysozyme\[O-NE-19]/554-728** | - | V21 | N2 |
| **WZO82436.1/1-820putativebifunctionalchitinase/lysozyme\[SH-6A]/142-304** | - | L6 | - |
| **WZO82436.1/1-820putativebifunctionalchitinase/lysozyme\[SH-6A]/553-731** | - | V21 | Q2 |
| **WZO92132.1/1-827putativebifunctionalchitinase/lysozyme\[NE-O-7-s]/139-304** | - | L9 | - |
| **WZO92132.1/1-827putativebifunctionalchitinase/lysozyme\[NE-O-7-s]/560-734** | - | V21 | N2 |
| **WZO89127.1/1-809putativebifunctionalchitinase/lysozyme\[O-NE-23]/139-304** | - | L9 | - |
| **WZO89127.1/1-809putativebifunctionalchitinase/lysozyme\[O-NE-23]/542-716** | - | V21 | N2 |
| **WZO85876.1/1-819putativebifunctionalchitinase/lysozyme\[O-NE-14]/142-304** | - | L6 | - |
| **WZO85876.1/1-819putativebifunctionalchitinase/lysozyme\[O-NE-14]/552-726** | - | V21 | K2 |
| **WZO88848.1/1-822putativebifunctionalchitinase/lysozyme\[O-NE-22]/142-304** | - | L6 | - |
| **WZO88848.1/1-822putativebifunctionalchitinase/lysozyme\[O-NE-22]/555-729** | - | V21 | N2 |
| **WZO83897.1/1-820putativebifunctionalchitinase/lysozyme\[XZ-6E]/142-304** | - | L6 | - |
| **WZO83897.1/1-820putativebifunctionalchitinase/lysozyme\[XZ-6E]/553-732** | - | V21 | Q2 |
| **YP_009665302.1/1-806putativebifunctionalchitinase/lysozyme\[ParameciumbursariaChlorellavirusNYs1]/142-304** | - | L6 | - |
| **YP_009665302.1/1-806putativebifunctionalchitinase/lysozyme\[ParameciumbursariaChlorellavirusNYs1]/539-715** | - | V21 | **C2** |
| **WZO87550.1/1-808putativebifunctionalchitinase/lysozyme\[O-NE-18]/143-303** | - | L4 | - |
| **WZO87550.1/1-808putativebifunctionalchitinase/lysozyme\[O-NE-18]/541-715** | - | V21 | N2 |
| **AGE54806.1/1-806putativebifunctionalchitinase/lysozyme\[ParameciumbursariaChlorellavirusMA1D]/142-304** | - | L6 | - |
| **AGE54806.1/1-806putativebifunctionalchitinase/lysozyme\[ParameciumbursariaChlorellavirusMA1D]/539-715** | - | V21 | **C2** |
| **WZO84706.1/1-831putativebifunctionalchitinase/lysozyme\[O-NE-11]/142-304** | - | L6 | - |
| **WZO84706.1/1-831putativebifunctionalchitinase/lysozyme\[O-NE-11]/564-738** | - | V21 | K2 |
| **WZO80449.1/1-806putativebifunctionalchitinase/lysozyme\[WNE-10B-S1]/142-304** | - | L6 | - |
| **WZO80449.1/1-806putativebifunctionalchitinase/lysozyme\[WNE-10B-S1]/539-714** | - | V21 | **C2** |
| **WZO85478.1/1-830putativebifunctionalchitinase/lysozyme\[O-NE-13]/142-304** | - | L6 | - |
| **WZO85478.1/1-830putativebifunctionalchitinase/lysozyme\[O-NE-13]/564-737** | - | V20 | S1 |
| **WZO76146.1/1-816bifunctionalchitinase/lysozyme\[40-NE-5]/142-304** | - | L6 | - |
| **WZO76146.1/1-816bifunctionalchitinase/lysozyme\[40-NE-5]/549-723** | - | S18 | G2 |
| **WZO76561.1/1-816putativebifunctionalchitinase/lysozyme\[41-NE-4]/142-304** | - | L6 | - |
| **WZO76561.1/1-816putativebifunctionalchitinase/lysozyme\[41-NE-4]/549-723** | - | S18 | G2 |
| **WZO78129.1/1-816chitinase\[NE-40ā??2-s]/142-304** | - | L6 | - |
| **WZO78129.1/1-816chitinase\[NE-40ā??2-s]/549-723** | - | S18 | G2 |
| **WZO79003.1/1-816putativebifunctionalchitinase/lysozyme\[NE-41-3-s]/142-304** | - | L6 | - |
| **WZO79003.1/1-816putativebifunctionalchitinase/lysozyme\[NE-41-3-s]/549-723** | - | S18 | G2 |
| **WZO80062.1/1-823putativebifunctionalchitinase/lysozyme\[N-NE-5]/142-304** | - | L6 | - |
| **WZO80062.1/1-823putativebifunctionalchitinase/lysozyme\[N-NE-5]/549-723** | - | S18 | G2 |
| **AGE58174.1/1-816putativebifunctionalchitinase/lysozyme\[ParameciumbursariaChlorellavirusNW665.2]/140-305** | - | L9 | - |

| | | | |
|---|---|---|---|
| AGE58174.1/1-816putativebifunctionalchitinase/lysozyme\[ParameciumbursariaChlorellavirusNW665.2]/550-723 | - | V20 | S1 |
| YP_001426411.1/1-805hypotheticalproteinFR483_N779R\[ParameciumbursariaChlorellavirusFR483]/139-304 | - | L9 | - |
| YP_001426411.1/1-805hypotheticalproteinFR483_N779R\[ParameciumbursariaChlorellavirusFR483]/539-712 | - | V20 | S1 |
| WZO86764.1/1-525chitinase\[O-NE-16]/142-304 | - | L6 | - |
| WZO93853.1/1-525chitinase\[OSyNE-4B-S1]/142-304 | - | L6 | - |
| WZO91298.1/1-445chitinase\[O-NE-28]/147-309 | - | L6 | - |
| WZO89577.1/1-445chitinase\[O-NE-24]/147-309 | - | L6 | - |
| WZO90501.1/1-525chitinase\[O-NE-26]/142-304 | - | L6 | - |
| WZO77210.1/1-525chitinase\[41-NE-6]/142-304 | - | L6 | - |
| WZO77693.1/1-525chitinase\[NE-40-1-m]/142-304 | - | L6 | - |
| WZO78681.1/1-525chitinase\[NE-41-2-m]/142-304 | - | L6 | - |
| WZO85099.1/1-530chitinase\[O-NE-12]/147-309 | - | L6 | - |
| YP_009325635.1/1-525Chitinase\[OnlySyngenNebraskavirus5]/142-304 | - | L6 | - |
| WZO88270.1/1-528chitinase\[O-NE-20]/147-309 | - | L6 | - |
| WZO92747.1/1-528chitinase\[NE-O-9-L]/147-309 | - | L6 | - |
| WZO93306.1/1-528chitinase\[OSyNE-4B-L2]/147-309 | - | L6 | - |
| WZO87136.1/1-528chitinase\[O-NE-17]/147-309 | - | L6 | - |
| WZO92491.1/1-528chitinase\[NE-O-8-L]/147-309 | - | L6 | - |
| WZO95440.1/1-528chitinase\[OSyNE-ZA]/147-309 | - | L6 | - |
| WZO78440.1/1-532chitinase\[NE-41-1-L]/147-309 | - | L6 | - |
| WZO94180.1/1-539chitinase\[OSyNE-5A-L1]/147-309 | - | L6 | - |
| WZO84329.1/1-525chitinase\[O-NE-10]/142-304 | - | L6 | - |
| WZO94943.1/1-525chitinase\[OSyNE-5B-S1]/142-304 | - | L6 | - |
| WZO93480.1/1-525chitinase\[OSyNE-4B-M2]/142-304 | - | L6 | - |
| WZO94540.1/1-525chitinase\[OSyNE-5B-M2]/142-304 | - | L6 | - |
| WZO79465.1/1-530chitinase\[N-NE-4]/147-309 | - | L6 | - |
| XYW60054.1/1-521chitinase\[NE-P-3-s]/139-304 | - | L9 | - |
| XYW60732.1/1-521chitinase\[NE-P-4-fs]/139-304 | - | L9 | - |
| XYW61232.1/1-521chitinase\[NE-P-8-f]/139-304 | - | L9 | - |
| XYW62772.1/1-521chitinase\[P-NE-12]/139-304 | - | L9 | - |
| WZO86398.1/1-532chitinase\[O-NE-15]/144-309 | - | L9 | - |
| XOK27477.1/1-582putativebifunctionalchitinase/lysozyme\[CL-S-1-m]/153-304 | - | - | - |
| AGE49332.1/1-595putativebifunctionalchitinase/lysozyme\[AcanthocystisturfaceaChlorellavirusBr0604L]/166-317 | - | - | - |
| AGE48989.1/1-520putativebifunctionalchitinase/lysozyme\[ParameciumbursariaChlorellavirusAP110A]/139-304 | - | L9 | - |
| AGE50008.1/1-520putativebifunctionalchitinase/lysozyme\[ParameciumbursariaChlorellavirusCan18-4]/139-304 | - | L9 | - |
| XYW62298.1/1-520chitinase\[P-NE-10]/139-304 | - | L9 | - |
| AGE52349.1/1-523putativebifunctionalchitinase/lysozyme\[ParameciumbursariaChlorellavirusCVR-1]/143-304 | - | L5 | - |
| YP_009702007.1/1-523putativebifunctionalchitinase/lysozyme\[ParameciumbursariaChlorellavirusCVA-1]/143-304 | - | L5 | - |
| ABT14345.1/1-520hypotheticalproteinMT325_M791R\[ParameciumbursariachlorellavirusMT325]/139-304 | - | L9 | - |
| AGE51335.1/1-523putativebifunctionalchitinase/lysozyme\[ParameciumbursariaChlorellavirusCVG-1]/139-304 | - | L9 | - |
| XYW61920.1/1-523chitinase\[P-NE-9]/139-304 | - | L9 | - |
| XOK31597.1/1-580putativebifunctionalchitinase/lysozyme\[S-NE-11]/151-304 | - | - | - |
| XOK36306.1/1-580putativebifunctionalchitinase/lysozyme\[GNLD22]/152-304 | - | - | - |
| XOK36264.1/1-582putativebifunctionalchitinase/lysozyme\[LP-F3a-4a]/153-304 | - | - | - |
| XOK31528.1/1-582putativebifunctionalchitinase/lysozyme\[S-NE-10]/153-304 | - | - | - |
| XOK28546.1/1-582putativebifunctionalchitinase/lysozyme\[NES-4B-L1]/153-304 | - | - | - |
| YP_001427295.1/1-595hypotheticalproteinATCV1_Z814L\[Acanthocystisturfaceachlorellavirus1]/166-317 | - | - | - |
| AGE56132.1/1-601putativebifunctionalchitinase/lysozyme\[AcanthocystisturfaceaChlorellavirusMO0605SPH]/172-323 | - | - | - |
| XOK34753.1/1-421putativebifunctionalchitinase/lysozyme\[S-NE-19]/153-304 | - | - | - |
| AGE60252.1/1-601putativebifunctionalchitinase/lysozyme\[AcanthocystisturfaceaChlorellavirusWI0606]/172-323 | - | - | - |
| AGE57134.1/1-601putativebifunctionalchitinase/lysozyme\[AcanthocystisturfaceaChlorellavirusNE-JV-3]/172-323 | - | - | - |
| XOK29995.1/1-582putativebifunctionalchitinase/lysozyme\[NES-5A-S1]/153-304 | - | - | - |
| AGE49664.1/1-595putativebifunctionalchitinase/lysozyme\[AcanthocystisturfaceaChlorellavirusCan0610SP]/166-317 | - | - | - |
| AGE59935.1/1-595putativebifunctionalchitinase/lysozyme\[AcanthocystisturfaceaChlorellavirusTN603.4.2]/189-317 | - | - | - |
| AGE55823.1/1-596putativebifunctionalchitinase/lysozyme\[AcanthocystisturfaceaChlorellavirusMN0810.1]/185-317 | - | - | - |
| XOK35208.1/1-580putativebifunctionalchitinase/lysozyme\[S-NE-22]/149-304 | - | - | - |
| AGE50333.1/1-580putativebifunctionalchitinase/lysozyme\[AcanthocystisturfaceaChlorellavirusCanal-1]/149-304 | - | - | - |
| XOK28216.1/1-580putativebifunctionalchitinase/lysozyme\[NES-4A-S1]/149-304 | - | - | - |
| XOK30852.1/1-582putativebifunctionalchitinase/lysozyme\[S-NE-9]/153-304 | - | - | - |
| XOK34326.1/1-580putativebifunctionalchitinase/lysozyme\[S-NE-18]/152-304 | - | - | - |
| AGE56814.1/1-582putativebifunctionalchitinase/lysozyme\[AcanthocystisturfaceaChlorellavirusNE-JV-2]/153-304 | - | - | - |
| AGE56814.1/1-582putativebifunctionalchitinase/lysozyme\[AcanthocystisturfaceaChlorellavirusNE-JV-2]/153-304 | - | - | - |
| XOK32790.1/1-582putativebifunctionalchitinase/lysozyme\[S-NE-15]/153-304 | - | - | - |
| XOK35891.1/1-582putativebifunctionalchitinase/lysozyme\[S-NE-23]/153-304 | - | - | - |
| XOK33538.1/1-582putativebifunctionalchitinase/lysozyme\[S-NE-16]/153-304 | - | - | - |
| XOK33601.1/1-582putativebifunctionalchitinase/lysozyme\[S-NE-17]/153-304 | - | - | - |
| XOK32004.1/1-582putativebifunctionalchitinase/lysozyme\[S-NE-12]/153-304 | - | - | - |
| XOK35119.1/1-582putativebifunctionalchitinase/lysozyme\[S-NE-20]/175-304 | - | - | - |
| XOK27846.1/1-580putativebifunctionalchitinase/lysozyme\[NES-4A-M1]/152-304 | - | - | - |
| XOK30419.1/1-582chitinase\[S-NE-7]/153-304 | - | - | - |
| AGE59594.1/1-582putativebifunctionalchitinase/lysozyme\[AcanthocystisturfaceaChlorellavirusOR0704.3]/153-304 | - | - | - |

| | | | |
|---|---|---|---|
| XOK32390.1/1-589putativebifunctionalchitinase/lysozyme\[S-NE-13]/153-304 | - | - | - |
| AGE57849.1/1-601putativebifunctionalchitinase/lysozyme\[AcanthocystisturfaceaChlorellavirusNTS-1]/172-323 | - | - | - |

**Table S6:** Prevalence of coevolving residues on chlorovirus chitinases in set 4.

| Sequence | G907 | W965 |
|---|---|---|
| WZO81342.1/1-840bifunctionalchitinase/lysozyme\[CA-4B]/142-304 | D162 | - |
| WZO81342.1/1-840bifunctionalchitinase/lysozyme\[CA-4B]/573-747 | T171 | E173 |
| WZO81006.1/1-840putativebifunctionalchitinase/lysozyme\[CA-4A]/142-304 | D162 | - |
| WZO81006.1/1-840putativebifunctionalchitinase/lysozyme\[CA-4A]/573-747 | T171 | E173 |
| WZO82266.1/1-840putativebifunctionalchitinase/lysozyme\[NY-2C]/142-304 | D162 | - |
| WZO82266.1/1-840putativebifunctionalchitinase/lysozyme\[NY-2C]/573-747 | T171 | E173 |
| WZO80816.1/1-830putativebifunctionalchitinase/lysozyme\[WNE-11A-L2]/142-304 | D162 | - |
| WZO80816.1/1-830putativebifunctionalchitinase/lysozyme\[WNE-11A-L2]/563-740 | T171 | E173 |
| AGE57241.1/1-815putativebifunctionalchitinase/lysozyme\[ParameciumbursariaChlorellavirusNE-JV-4]/142-304 | D162 | - |
| AGE57241.1/1-815putativebifunctionalchitinase/lysozyme\[ParameciumbursariaChlorellavirusNE-JV-4]/548-725 | T171 | E173 |
| WZO75107.1/1-840chitinase\[40-NE-3]/142-304 | D162 | - |
| WZO75107.1/1-840chitinase\[40-NE-3]/573-747 | T171 | E173 |
| WZO75726.1/1-840putativebifunctionalchitinase/lysozyme\[40-NE-4]/142-304 | D162 | - |
| WZO75726.1/1-840putativebifunctionalchitinase/lysozyme\[40-NE-4]/573-747 | T171 | E173 |
| WZO76741.1/1-840chitinase\[41-NE-5]/142-304 | D162 | - |
| WZO76741.1/1-840chitinase\[41-NE-5]/573-747 | T171 | E173 |
| WZO83158.1/1-830chitinase\[XZ-4C]/142-304 | D162 | - |
| WZO83158.1/1-830chitinase\[XZ-4C]/563-737 | T171 | E173 |
| AGE53811.1/1-835putativebifunctionalchitinase/lysozyme\[ParameciumbursariaChlorellavirusIL3A]/142-304 | D162 | - |
| AGE53811.1/1-835putativebifunctionalchitinase/lysozyme\[ParameciumbursariaChlorellavirusIL3A]/567-745 | T172 | E174 |
| AGE54506.1/1-817putativebifunctionalchitinase/lysozyme\[ParameciumbursariaChlorellavirusKS1B]/142-304 | D162 | - |
| AGE54506.1/1-817putativebifunctionalchitinase/lysozyme\[ParameciumbursariaChlorellavirusKS1B]/550-727 | T171 | E173 |
| AGE48399.1/1-835putativebifunctionalchitinase/lysozyme\[ParameciumbursariaChlorellavirusAN69C]/142-304 | D162 | - |
| AGE48399.1/1-835putativebifunctionalchitinase/lysozyme\[ParameciumbursariaChlorellavirusAN69C]/568-742 | T171 | E173 |
| WZO91574.1/1-809putativebifunctionalchitinase/lysozyme\[O-NE-29]/139-304 | D165 | - |
| WZO91574.1/1-809putativebifunctionalchitinase/lysozyme\[O-NE-29]/542-716 | N171 | E173 |
| WZO81686.1/1-840chitinase\[NC-1A]/142-304 | D162 | - |
| WZO81686.1/1-840chitinase\[NC-1A]/573-747 | T171 | E173 |
| NP_048529.2/1-830Chitinase\[ParameciumbursariaChlorellavirus1]/142-304 | D162 | - |
| NP_048529.2/1-830Chitinase\[ParameciumbursariaChlorellavirus1]/563-737 | T171 | E173 |
| WZO82806.1/1-830chitinase\[XZ-3A]/142-304 | D162 | - |
| WZO82806.1/1-830chitinase\[XZ-3A]/563-737 | T171 | E173 |
| WZO90762.1/1-809putativebifunctionalchitinase/lysozyme\[O-NE-27]/139-304 | D165 | - |
| WZO90762.1/1-809putativebifunctionalchitinase/lysozyme\[O-NE-27]/542-716 | T171 | E173 |
| WZO83510.1/1-820putativebifunctionalchitinase/lysozyme\[XZ-5C]/142-304 | D162 | - |
| WZO83510.1/1-820putativebifunctionalchitinase/lysozyme\[XZ-5C]/553-731 | T171 | E173 |
| WZO89952.1/1-822putativebifunctionalchitinase/lysozyme\[O-NE-25]/142-304 | D162 | - |
| WZO89952.1/1-822putativebifunctionalchitinase/lysozyme\[O-NE-25]/555-729 | N171 | E173 |
| AGE51440.1/1-836putativebifunctionalchitinase/lysozyme\[ParameciumbursariaChlorellavirusCviKI]/142-304 | D162 | - |
| AGE51440.1/1-836putativebifunctionalchitinase/lysozyme\[ParameciumbursariaChlorellavirusCviKI]/569-748 | N171 | E173 |
| AGE52455.1/1-826putativebifunctionalchitinase/lysozyme\[ParameciumbursariaChlorellavirusCvsA1]/142-304 | D162 | - |
| AGE52455.1/1-826putativebifunctionalchitinase/lysozyme\[ParameciumbursariaChlorellavirusCvsA1]/559-739 | T171 | E173 |
| WZO87987.1/1-821putativebifunctionalchitinase/lysozyme\[O-NE-19]/142-304 | D162 | - |
| WZO87987.1/1-821putativebifunctionalchitinase/lysozyme\[O-NE-19]/554-728 | N171 | E173 |
| WZO82436.1/1-820putativebifunctionalchitinase/lysozyme\[SH-6A]/142-304 | D162 | - |
| WZO82436.1/1-820putativebifunctionalchitinase/lysozyme\[SH-6A]/553-731 | T171 | E173 |
| WZO92132.1/1-827putativebifunctionalchitinase/lysozyme\[NE-O-7-s]/139-304 | D165 | - |
| WZO92132.1/1-827putativebifunctionalchitinase/lysozyme\[NE-O-7-s]/560-734 | T171 | E173 |
| WZO89127.1/1-809putativebifunctionalchitinase/lysozyme\[O-NE-23]/139-304 | D165 | - |
| WZO89127.1/1-809putativebifunctionalchitinase/lysozyme\[O-NE-23]/542-716 | N171 | E173 |
| WZO85876.1/1-819putativebifunctionalchitinase/lysozyme\[O-NE-14]/142-304 | D162 | - |
| WZO85876.1/1-819putativebifunctionalchitinase/lysozyme\[O-NE-14]/552-726 | T171 | E173 |
| WZO88848.1/1-822putativebifunctionalchitinase/lysozyme\[O-NE-22]/142-304 | D162 | - |
| WZO88848.1/1-822putativebifunctionalchitinase/lysozyme\[O-NE-22]/555-729 | N171 | E173 |
| WZO83897.1/1-820putativebifunctionalchitinase/lysozyme\[XZ-6E]/142-304 | D162 | - |
| WZO83897.1/1-820putativebifunctionalchitinase/lysozyme\[XZ-6E]/553-732 | T171 | E173 |
| YP_009665302.1/1-806putativebifunctionalchitinase/lysozyme\[ParameciumbursariaChlorellavirusNYs1]/142-304 | D162 | - |
| YP_009665302.1/1-806putativebifunctionalchitinase/lysozyme\[ParameciumbursariaChlorellavirusNYs1]/539-715 | T171 | E173 |
| WZO87550.1/1-808putativebifunctionalchitinase/lysozyme\[O-NE-18]/143-303 | D160 | - |
| WZO87550.1/1-808putativebifunctionalchitinase/lysozyme\[O-NE-18]/541-715 | N171 | E173 |
| AGE54806.1/1-806putativebifunctionalchitinase/lysozyme\[ParameciumbursariaChlorellavirusMA1D]/142-304 | D162 | - |

| | | |
|---|---|---|
| AGE54806.1/1-806putativebifunctionalchitinase/lysozyme\[ParameciumbursariaChlorellavirusMA1D]/539-715 | T171 | E173 |
| WZO84706.1/1-831putativebifunctionalchitinase/lysozyme\[O-NE-11]/142-304 | D162 | - |
| WZO84706.1/1-831putativebifunctionalchitinase/lysozyme\[O-NE-11]/564-738 | T171 | E173 |
| WZO80449.1/1-806putativebifunctionalchitinase/lysozyme\[WNE-10B-S1]/142-304 | D162 | - |
| WZO80449.1/1-806putativebifunctionalchitinase/lysozyme\[WNE-10B-S1]/539-714 | T171 | E173 |
| WZO85478.1/1-830putativebifunctionalchitinase/lysozyme\[O-NE-13]/142-304 | D162 | - |
| WZO85478.1/1-830putativebifunctionalchitinase/lysozyme\[O-NE-13]/564-737 | N170 | E172 |
| WZO76146.1/1-816bifunctionalchitinase/lysozyme\[40-NE-5]/142-304 | D162 | - |
| WZO76146.1/1-816bifunctionalchitinase/lysozyme\[40-NE-5]/549-723 | T171 | E173 |
| WZO76561.1/1-816putativebifunctionalchitinase/lysozyme\[41-NE-4]/142-304 | D162 | - |
| WZO76561.1/1-816putativebifunctionalchitinase/lysozyme\[41-NE-4]/549-723 | T171 | E173 |
| WZO78129.1/1-816chitinase\[NE-40ã??2-s]/142-304 | D162 | - |
| WZO78129.1/1-816chitinase\[NE-40ã??2-s]/549-723 | T171 | E173 |
| WZO79003.1/1-816putativebifunctionalchitinase/lysozyme\[NE-41-3-s]/142-304 | D162 | - |
| WZO79003.1/1-816putativebifunctionalchitinase/lysozyme\[NE-41-3-s]/549-723 | T171 | E173 |
| WZO80062.1/1-823putativebifunctionalchitinase/lysozyme\[N-NE-5]/142-304 | D162 | - |
| WZO80062.1/1-823putativebifunctionalchitinase/lysozyme\[N-NE-5]/549-723 | T171 | E173 |
| AGE58174.1/1-816putativebifunctionalchitinase/lysozyme\[ParameciumbursariaChlorellavirusNW665.2]/140-305 | D165 | - |
| AGE58174.1/1-816putativebifunctionalchitinase/lysozyme\[ParameciumbursariaChlorellavirusNW665.2]/550-723 | N170 | E172 |
| YP_001426411.1/1-805hypotheticalproteinFR483_N779R\[ParameciumbursariaChlorellavirusFR483]/139-304 | D165 | - |
| YP_001426411.1/1-805hypotheticalproteinFR483_N779R\[ParameciumbursariaChlorellavirusFR483]/539-712 | N170 | E172 |
| WZO86764.1/1-525chitinase\[O-NE-16]/142-304 | D162 | - |
| WZO93853.1/1-525chitinase\[OSyNE-4B-S1]/142-304 | D162 | - |
| WZO91298.1/1-445chitinase\[O-NE-28]/147-309 | D162 | - |
| WZO89577.1/1-445chitinase\[O-NE-24]/147-309 | D162 | - |
| WZO90501.1/1-525chitinase\[O-NE-26]/142-304 | D162 | - |
| WZO77210.1/1-525chitinase\[41-NE-6]/142-304 | D162 | - |
| WZO77693.1/1-525chitinase\[NE-40-1-m]/142-304 | D162 | - |
| WZO78681.1/1-525chitinase\[NE-41-2-m]/142-304 | D162 | - |
| WZO85099.1/1-530chitinase\[O-NE-12]/147-309 | D162 | - |
| YP_009325635.1/1-525Chitinase\[OnlySyngenNebraskavirus5]/142-304 | D162 | - |
| WZO88270.1/1-528chitinase\[O-NE-20]/147-309 | D162 | - |
| WZO92747.1/1-528chitinase\[NE-O-9-L]/147-309 | D162 | - |
| WZO93306.1/1-528chitinase\[OSyNE-4B-L2]/147-309 | D162 | - |
| WZO87136.1/1-528chitinase\[O-NE-17]/147-309 | D162 | - |
| WZO92491.1/1-528chitinase\[NE-O-8-L]/147-309 | D162 | - |
| WZO95440.1/1-528chitinase\[OSyNE-ZA]/147-309 | D162 | - |
| WZO78440.1/1-532chitinase\[NE-41-1-L]/147-309 | D162 | - |
| WZO94180.1/1-539chitinase\[OSyNE-5A-L1]/147-309 | D162 | - |
| WZO84329.1/1-525chitinase\[O-NE-10]/142-304 | D162 | - |
| WZO94943.1/1-525chitinase\[OSyNE-5B-S1]/142-304 | D162 | - |
| WZO93480.1/1-525chitinase\[OSyNE-4B-M2]/142-304 | D162 | - |
| WZO94540.1/1-525chitinase\[OSyNE-5B-M2]/142-304 | D162 | - |
| WZO79465.1/1-530chitinase\[N-NE-4]/147-309 | D162 | - |
| XYW60054.1/1-521chitinase\[NE-P-3-s]/139-304 | D165 | - |
| XYW60732.1/1-521chitinase\[NE-P-4-fs]/139-304 | D165 | - |
| XYW61232.1/1-521chitinase\[NE-P-8-f]/139-304 | D165 | - |
| XYW62772.1/1-521chitinase\[P-NE-12]/139-304 | D165 | - |
| WZO86398.1/1-532chitinase\[O-NE-15]/144-309 | D165 | - |
| XOK27477.1/1-582putativebifunctionalchitinase/lysozyme\[CL-S-1-m]/153-304 | D151 | - |
| AGE49332.1/1-595putativebifunctionalchitinase/lysozyme\[AcanthocystisturfaceaChlorellavirusBr0604L]/166-317 | D151 | - |
| AGE48989.1/1-520putativebifunctionalchitinase/lysozyme\[ParameciumbursariaChlorellavirusAP110A]/139-304 | D165 | - |
| AGE50008.1/1-520putativebifunctionalchitinase/lysozyme\[ParameciumbursariaChlorellavirusCan18-4]/139-304 | D165 | - |
| XYW62298.1/1-520chitinase\[P-NE-10]/139-304 | D165 | - |
| AGE52349.1/1-523putativebifunctionalchitinase/lysozyme\[ParameciumbursariaChlorellavirusCVR-1]/143-304 | D161 | - |
| YP_009702007.1/1-523putativebifunctionalchitinase/lysozyme\[ParameciumbursariaChlorellavirusCVA-1]/143-304 | D161 | - |
| ABT14345.1/1-520hypotheticalproteinMT325_M791R\[ParameciumbursariachlorellavirusMT325]/139-304 | D165 | - |
| AGE51335.1/1-523putativebifunctionalchitinase/lysozyme\[ParameciumbursariaChlorellavirusCVG-1]/139-304 | D165 | - |
| XYW61920.1/1-523chitinase\[P-NE-9]/139-304 | D165 | - |
| XOK31597.1/1-580putativebifunctionalchitinase/lysozyme\[S-NE-11]/151-304 | D153 | - |
| XOK36306.1/1-580putativebifunctionalchitinase/lysozyme\[GNLD22]/152-304 | D152 | - |
| XOK36264.1/1-582putativebifunctionalchitinase/lysozyme\[LP-F3a-4a]/153-304 | D151 | - |
| XOK31528.1/1-582putativebifunctionalchitinase/lysozyme\[S-NE-10]/153-304 | D151 | - |
| XOK28546.1/1-582putativebifunctionalchitinase/lysozyme\[NES-4B-L1]/153-304 | D151 | - |
| YP_001427295.1/1-595hypotheticalproteinATCV1_Z814L\[Acanthocystisturfaceachlorellavirus1]/166-317 | D151 | - |
| AGE56132.1/1-601putativebifunctionalchitinase/lysozyme\[AcanthocystisturfaceaChlorellavirusMO0605SPH]/172-323 | D151 | - |
| XOK34753.1/1-421putativebifunctionalchitinase/lysozyme\[S-NE-19]/153-304 | D151 | - |
| AGE60252.1/1-601putativebifunctionalchitinase/lysozyme\[AcanthocystisturfaceaChlorellavirusWI0606]/172-323 | D151 | - |
| AGE57134.1/1-601putativebifunctionalchitinase/lysozyme\[AcanthocystisturfaceaChlorellavirusNE-JV-3]/172-323 | D151 | - |
| XOK29995.1/1-582putativebifunctionalchitinase/lysozyme\[NES-5A-S1]/153-304 | D151 | - |
| AGE49664.1/1-595putativebifunctionalchitinase/lysozyme\[AcanthocystisturfaceaChlorellavirusCan0610SP]/166-317 | D151 | - |

| | | |
|---|---|---|
| AGE59935.1/1-595putativebifunctionalchitinase/lysozyme\[AcanthocystisturfaceaChlorellavirusTN603.4.2]/189-317 | D128 | - |
| AGE55823.1/1-596putativebifunctionalchitinase/lysozyme\[AcanthocystisturfaceaChlorellavirusMN0810.1]/185-317 | D132 | - |
| XOK35208.1/1-580putativebifunctionalchitinase/lysozyme\[S-NE-22]/149-304 | D155 | - |
| AGE50333.1/1-580putativebifunctionalchitinase/lysozyme\[AcanthocystisturfaceaChlorellavirusCanal-1]/149-304 | D155 | - |
| XOK28216.1/1-580putativebifunctionalchitinase/lysozyme\[NES-4A-S1]/149-304 | D155 | - |
| XOK30852.1/1-582putativebifunctionalchitinase/lysozyme\[S-NE-9]/153-304 | D151 | - |
| XOK34326.1/1-580putativebifunctionalchitinase/lysozyme\[S-NE-18]/152-304 | D152 | - |
| AGE56814.1/1-582putativebifunctionalchitinase/lysozyme\[AcanthocystisturfaceaChlorellavirusNE-JV-2]/153-304 | D151 | - |
| AGE56814.1/1-582putativebifunctionalchitinase/lysozyme\[AcanthocystisturfaceaChlorellavirusNE-JV-2]/153-304 | D151 | - |
| XOK32790.1/1-582putativebifunctionalchitinase/lysozyme\[S-NE-15]/153-304 | D151 | - |
| XOK35891.1/1-582putativebifunctionalchitinase/lysozyme\[S-NE-23]/153-304 | D151 | - |
| XOK33538.1/1-582putativebifunctionalchitinase/lysozyme\[S-NE-16]/153-304 | D151 | - |
| XOK33601.1/1-582putativebifunctionalchitinase/lysozyme\[S-NE-17]/153-304 | D151 | - |
| XOK32004.1/1-582putativebifunctionalchitinase/lysozyme\[S-NE-12]/153-304 | D151 | - |
| XOK35119.1/1-582putativebifunctionalchitinase/lysozyme\[S-NE-20]/175-304 | D129 | - |
| XOK27846.1/1-580putativebifunctionalchitinase/lysozyme\[NES-4A-M1]/152-304 | D152 | - |
| XOK30419.1/1-582chitinase\[S-NE-7]/153-304 | D151 | - |
| AGE59594.1/1-582putativebifunctionalchitinase/lysozyme\[AcanthocystisturfaceaChlorellavirusOR0704.3]/153-304 | D151 | - |
| XOK32390.1/1-589putativebifunctionalchitinase/lysozyme\[S-NE-13]/153-304 | D151 | - |
| AGE57849.1/1-601putativebifunctionalchitinase/lysozyme\[AcanthocystisturfaceaChlorellavirusNTS-1]/172-323 | D151 | - |

**Table S7:** Prevalence of conserved residues on chlorovirus chitinases in set 4. Colored in green represents coevolved residues.

| Sequence | G534 | D560 | E563 | D889 | M895 | Y899 | D901 | K1148 | G1158 | Y1169 | G1175 | K1699 | G1741 | W1750 |
|---|---|---|---|---|---|---|---|---|---|---|---|---|---|---|
| WZO81342.1/1-840bifunctionalchitinase/lysozyme\[CA-4B]/142-304 | R86 | D90 | E92 | H151 | M156 | M158 | D159 | - | - | - | - | - | - | - |
| WZO81342.1/1-840bifunctionalchitinase/lysozyme\[CA-4B]/573-747 | N95 | D99 | E101 | H160 | M165 | M167 | D168 | - | - | - | - | - | - | - |
| WZO81006.1/1-840putativebifunctionalchitinase/lysozyme\[CA-4A]/142-304 | R86 | D90 | E92 | H151 | M156 | M158 | D159 | - | - | - | - | - | - | - |
| WZO81006.1/1-840putativebifunctionalchitinase/lysozyme\[CA-4A]/573-747 | N95 | D99 | E101 | H160 | M165 | M167 | D168 | - | - | - | - | - | - | - |
| WZO82266.1/1-840putativebifunctionalchitinase/lysozyme\[NY-2C]/142-304 | R86 | D90 | E92 | H151 | M156 | M158 | D159 | - | - | - | - | - | - | - |
| WZO82266.1/1-840putativebifunctionalchitinase/lysozyme\[NY-2C]/573-747 | N95 | D99 | E101 | H160 | M165 | M167 | D168 | - | - | - | - | - | - | - |
| WZO80816.1/1-830putativebifunctionalchitinase/lysozyme\[WNE-11A-L2]/142-304 | R86 | D90 | E92 | H151 | M156 | M158 | D159 | - | - | - | - | - | - | - |
| WZO80816.1/1-830putativebifunctionalchitinase/lysozyme\[WNE-11A-L2]/563-740 | N95 | D99 | E101 | H160 | M165 | M167 | D168 | - | - | - | - | - | - | - |
| AGE57241.1/1-815putativebifunctionalchitinase/lysozyme\[ParameciumbursariaChlorellavirusNE-JV-4]/142-304 | R86 | D90 | E92 | H151 | M156 | M158 | D159 | - | - | - | - | - | - | - |
| AGE57241.1/1-815putativebifunctionalchitinase/lysozyme\[ParameciumbursariaChlorellavirusNE-JV-4]/548-725 | N95 | D99 | E101 | H160 | M165 | M167 | D168 | - | - | - | - | - | - | - |
| WZO75107.1/1-840chitinase\[40-NE-3]/142-304 | R86 | D90 | E92 | H151 | M156 | M158 | D159 | - | - | - | - | - | - | - |
| WZO75107.1/1-840chitinase\[40-NE-3]/573-747 | N95 | D99 | E101 | H160 | M165 | M167 | D168 | - | - | - | - | - | - | - |
| WZO75726.1/1-840putativebifunctionalchitinase/lysozyme\[40-NE-4]/142-304 | R86 | D90 | E92 | H151 | M156 | M158 | D159 | - | - | - | - | - | - | - |
| WZO75726.1/1-840putativebifunctionalchitinase/lysozyme\[40-NE-4]/573-747 | N95 | D99 | E101 | H160 | M165 | M167 | D168 | - | - | - | - | - | - | - |
| WZO76741.1/1-840chitinase\[41-NE-5]/142-304 | R86 | D90 | E92 | H151 | M156 | M158 | D159 | - | - | - | - | - | - | - |
| WZO76741.1/1-840chitinase\[41-NE-5]/573-747 | N95 | D99 | E101 | H160 | M165 | M167 | D168 | - | - | - | - | - | - | - |
| WZO83158.1/1-830chitinase\[XZ-4C]/142-304 | R86 | D90 | E92 | H151 | M156 | M158 | D159 | - | - | - | - | - | - | - |
| WZO83158.1/1-830chitinase\[XZ-4C]/563-737 | N95 | D99 | E101 | H160 | M165 | M167 | D168 | - | - | - | - | - | - | - |
| AGE53811.1/1-835putativebifunctionalchitinase/lysozyme\[ParameciumbursariaChlorellavirusIL3A]/142-304 | R86 | D90 | E92 | H151 | M156 | M158 | D159 | - | - | - | - | - | - | - |
| AGE53811.1/1-835putativebifunctionalchitinase/lysozyme\[ParameciumbursariaChlorellavirusIL3A]/567-745 | N96 | D100 | E102 | H161 | M166 | M168 | D169 | - | - | - | - | - | - | - |
| AGE54506.1/1-817putativebifunctionalchitinase/lysozyme\[ParameciumbursariaChlorellavirusKS1B]/142-304 | R86 | D90 | E92 | H151 | M156 | M158 | D159 | - | - | - | - | - | - | - |
| AGE54506.1/1-817putativebifunctionalchitinase/lysozyme\[ParameciumbursariaChlorellavirusKS1B]/550-727 | N95 | D99 | E101 | H160 | M165 | M167 | D168 | - | - | - | - | - | - | - |
| AGE48399.1/1-835putativebifunctionalchitinase/lysozyme\[ParameciumbursariaChlorellavirusAN69C]/142-304 | R86 | D90 | E92 | H151 | M156 | M158 | D159 | - | - | - | - | - | - | - |
| AGE48399.1/1-835putativebifunctionalchitinase/lysozyme\[ParameciumbursariaChlorellavirusAN69C]/568-742 | N95 | D99 | E101 | H160 | M165 | M167 | D168 | - | - | - | - | - | - | - |
| WZO91574.1/1-809putativebifunctionalchitinase/lysozyme\[O-NE-29]/139-304 | R89 | D93 | E95 | H154 | M159 | M161 | D162 | - | - | - | - | - | - | - |
| WZO91574.1/1-809putativebifunctionalchitinase/lysozyme\[O-NE-29]/542-716 | H95 | D99 | E101 | H160 | M165 | M167 | D168 | - | - | - | - | - | - | - |
| WZO81686.1/1-840chitinase\[NC-1A]/142-304 | R86 | D90 | E92 | H151 | M156 | M158 | D159 | - | - | - | - | - | - | - |
| WZO81686.1/1-840chitinase\[NC-1A]/573-747 | N95 | D99 | E101 | H160 | M165 | M167 | D168 | - | - | - | - | - | - | - |
| NP_048529.2/1-830Chitinase\[ParameciumbursariaChlorellavirus1]/142-304 | R86 | D90 | E92 | H151 | M156 | M158 | D159 | - | - | - | - | - | - | - |
| NP_048529.2/1-830Chitinase\[ParameciumbursariaChlorellavirus1]/563-737 | N95 | D99 | E101 | H160 | M165 | M167 | D168 | - | - | - | - | - | - | - |
| WZO82806.1/1-830chitinase\[XZ-3A]/142-304 | R86 | D90 | E92 | H151 | M156 | M158 | D159 | - | - | - | - | - | - | - |
| WZO82806.1/1-830chitinase\[XZ-3A]/563-737 | N95 | D99 | E101 | H160 | M165 | M167 | D168 | - | - | - | - | - | - | - |
| WZO90762.1/1-809putativebifunctionalchitinase/lysozyme\[O-NE-27]/139-304 | R89 | D93 | E95 | H154 | M159 | M161 | D162 | - | - | - | - | - | - | - |
| WZO90762.1/1-809putativebifunctionalchitinase/lysozyme\[O-NE-27]/542-716 | H95 | D99 | E101 | H160 | M165 | M167 | D168 | - | - | - | - | - | - | - |
| WZO83510.1/1-820putativebifunctionalchitinase/lysozyme\[XZ-5C]/142-304 | R86 | D90 | E92 | H151 | M156 | M158 | D159 | - | - | - | - | - | - | - |
| WZO83510.1/1-820putativebifunctionalchitinase/lysozyme\[XZ-5C]/553-731 | N95 | D99 | E101 | H160 | M165 | M167 | D168 | - | - | - | - | - | - | - |
| WZO89952.1/1-822putativebifunctionalchitinase/lysozyme\[O-NE-25]/142-304 | R86 | D90 | E92 | H151 | M156 | M158 | D159 | - | - | - | - | - | - | - |
| WZO89952.1/1-822putativebifunctionalchitinase/lysozyme\[O-NE-25]/555-729 | H95 | D99 | E101 | H160 | M165 | M167 | D168 | - | - | - | - | - | - | - |
| AGE51440.1/1-836putativebifunctionalchitinase/lysozyme\[ParameciumbursariaChlorellavirusCviKI]/142-304 | R86 | D90 | E92 | Y151 | M156 | M158 | D159 | - | - | - | - | - | - | - |
| AGE51440.1/1-836putativebifunctionalchitinase/lysozyme\[ParameciumbursariaChlorellavirusCviKI]/569-748 | N95 | D99 | E101 | H160 | M165 | M167 | D168 | - | - | - | - | - | - | - |
| AGE52455.1/1-826putativebifunctionalchitinase/lysozyme\[ParameciumbursariaChlorellavirusCvsA1]/142-304 | R86 | D90 | E92 | H151 | M156 | M158 | D159 | - | - | - | - | - | - | - |
| AGE52455.1/1-826putativebifunctionalchitinase/lysozyme\[ParameciumbursariaChlorellavirusCvsA1]/559-739 | N95 | D99 | E101 | H160 | M165 | M167 | D168 | - | - | - | - | - | - | - |

| | | | | | | | | | | | | | | |
|---|---|---|---|---|---|---|---|---|---|---|---|---|---|---|
| WZO87987.1/1-821putativebifunctionalchitinase/lysozyme\[O-NE-19]/142-304 | H86 | D90 | E92 | H151 | M156 | M158 | D159 | - | - | - | - | - | - | - |
| WZO87987.1/1-821putativebifunctionalchitinase/lysozyme\[O-NE-19]/554-728 | H95 | D99 | E101 | H160 | M165 | M167 | D168 | - | - | - | - | - | - | - |
| WZO82436.1/1-820putativebifunctionalchitinase/lysozyme\[SH-6A]/142-304 | R86 | D90 | E92 | H151 | M156 | M158 | D159 | - | - | - | - | - | - | - |
| WZO82436.1/1-820putativebifunctionalchitinase/lysozyme\[SH-6A]/553-731 | N95 | D99 | E101 | H160 | M165 | M167 | D168 | - | - | - | - | - | - | - |
| WZO92132.1/1-827putativebifunctionalchitinase/lysozyme\[NE-O-7-s]/139-304 | R89 | D93 | E95 | H154 | M159 | M161 | D162 | - | - | - | - | - | - | - |
| WZO92132.1/1-827putativebifunctionalchitinase/lysozyme\[NE-O-7-s]/560-734 | H95 | D99 | E101 | H160 | M165 | M167 | D168 | - | - | - | - | - | - | - |
| WZO89127.1/1-809putativebifunctionalchitinase/lysozyme\[O-NE-23]/139-304 | R89 | D93 | E95 | H154 | M159 | M161 | D162 | - | - | - | - | - | - | - |
| WZO89127.1/1-809putativebifunctionalchitinase/lysozyme\[O-NE-23]/542-716 | H95 | D99 | E101 | H160 | M165 | M167 | D168 | - | - | - | - | - | - | - |
| WZO85876.1/1-819putativebifunctionalchitinase/lysozyme\[O-NE-14]/142-304 | R86 | D90 | E92 | H151 | M156 | M158 | D159 | - | - | - | - | - | - | - |
| WZO85876.1/1-819putativebifunctionalchitinase/lysozyme\[O-NE-14]/552-726 | N95 | D99 | E101 | H160 | M165 | M167 | D168 | - | - | - | - | - | - | - |
| WZO88848.1/1-822putativebifunctionalchitinase/lysozyme\[O-NE-22]/142-304 | H86 | D90 | E92 | H151 | M156 | M158 | D159 | - | - | - | - | - | - | - |
| WZO88848.1/1-822putativebifunctionalchitinase/lysozyme\[O-NE-22]/555-729 | H95 | D99 | E101 | H160 | M165 | M167 | D168 | - | - | - | - | - | - | - |
| WZO83897.1/1-820putativebifunctionalchitinase/lysozyme\[XZ-6E]/142-304 | R86 | D90 | E92 | H151 | M156 | M158 | D159 | - | - | - | - | - | - | - |
| WZO83897.1/1-820putativebifunctionalchitinase/lysozyme\[XZ-6E]/553-732 | N95 | D99 | E101 | H160 | M165 | M167 | D168 | - | - | - | - | - | - | - |
| YP_009665302.1/1-806putativebifunctionalchitinase/lysozyme\[ParameciumbursariaChlorellavirusNYs1]/142-304 | R86 | D90 | E92 | H151 | M156 | M158 | D159 | - | - | - | - | - | - | - |
| YP_009665302.1/1-806putativebifunctionalchitinase/lysozyme\[ParameciumbursariaChlorellavirusNYs1]/539-715 | Y95 | D99 | E101 | H160 | M165 | M167 | D168 | - | - | - | - | - | - | - |
| WZO87550.1/1-808putativebifunctionalchitinase/lysozyme\[O-NE-18]/143-303 | R84 | D88 | E90 | H149 | M154 | M156 | D157 | - | - | - | - | - | - | - |
| WZO87550.1/1-808putativebifunctionalchitinase/lysozyme\[O-NE-18]/541-715 | H95 | D99 | E101 | H160 | M165 | M167 | D168 | - | - | - | - | - | - | - |
| AGE54806.1/1-806putativebifunctionalchitinase/lysozyme\[ParameciumbursariaChlorellavirusMA1D]/142-304 | R86 | D90 | E92 | H151 | M156 | M158 | D159 | - | - | - | - | - | - | - |
| AGE54806.1/1-806putativebifunctionalchitinase/lysozyme\[ParameciumbursariaChlorellavirusMA1D]/539-715 | Y95 | D99 | E101 | H160 | M165 | M167 | D168 | - | - | - | - | - | - | - |
| WZO84706.1/1-831putativebifunctionalchitinase/lysozyme\[O-NE-11]/142-304 | R86 | D90 | E92 | H151 | M156 | M158 | D159 | - | - | - | - | - | - | - |
| WZO84706.1/1-831putativebifunctionalchitinase/lysozyme\[O-NE-11]/564-738 | N95 | D99 | E101 | H160 | M165 | M167 | D168 | - | - | - | - | - | - | - |
| WZO80449.1/1-806putativebifunctionalchitinase/lysozyme\[WNE-10B-S1]/142-304 | R86 | D90 | E92 | H151 | M156 | M158 | D159 | - | - | - | - | - | - | - |
| WZO80449.1/1-806putativebifunctionalchitinase/lysozyme\[WNE-10B-S1]/539-714 | Y95 | D99 | E101 | H160 | M165 | M167 | D168 | - | - | - | - | - | - | - |
| WZO85478.1/1-830putativebifunctionalchitinase/lysozyme\[O-NE-13]/142-304 | R86 | D90 | E92 | Y151 | M156 | M158 | D159 | - | - | - | - | - | - | - |
| WZO85478.1/1-830putativebifunctionalchitinase/lysozyme\[O-NE-13]/564-737 | N94 | D98 | E100 | H159 | M164 | M166 | D167 | - | - | - | - | - | - | - |
| WZO76146.1/1-816bifunctionalchitinase/lysozyme\[40-NE-5]/142-304 | R86 | D90 | E92 | H151 | M156 | M158 | D159 | - | - | - | - | - | - | - |
| WZO76146.1/1-816bifunctionalchitinase/lysozyme\[40-NE-5]/549-723 | N95 | D99 | E101 | H160 | M165 | M167 | D168 | - | - | - | - | - | - | - |
| WZO76561.1/1-816putativebifunctionalchitinase/lysozyme\[41-NE-4]/142-304 | R86 | D90 | E92 | H151 | M156 | M158 | D159 | - | - | - | - | - | - | - |
| WZO76561.1/1-816putativebifunctionalchitinase/lysozyme\[41-NE-4]/549-723 | N95 | D99 | E101 | H160 | M165 | M167 | D168 | - | - | - | - | - | - | - |
| WZO78129.1/1-816chitinase\[NE-40â??2-s]/142-304 | R86 | D90 | E92 | H151 | M156 | M158 | D159 | - | - | - | - | - | - | - |
| WZO78129.1/1-816chitinase\[NE-40â??2-s]/549-723 | N95 | D99 | E101 | H160 | M165 | M167 | D168 | - | - | - | - | - | - | - |
| WZO79003.1/1-816putativebifunctionalchitinase/lysozyme\[NE-41-3-s]/142-304 | R86 | D90 | E92 | H151 | M156 | M158 | D159 | - | - | - | - | - | - | - |
| WZO79003.1/1-816putativebifunctionalchitinase/lysozyme\[NE-41-3-s]/549-723 | N95 | D99 | E101 | H160 | M165 | M167 | D168 | - | - | - | - | - | - | - |
| WZO80062.1/1-823putativebifunctionalchitinase/lysozyme\[N-NE-5]/142-304 | R86 | D90 | E92 | H151 | M156 | M158 | D159 | - | - | - | - | - | - | - |
| WZO80062.1/1-823putativebifunctionalchitinase/lysozyme\[N-NE-5]/549-723 | N95 | D99 | E101 | H160 | M165 | M167 | D168 | - | - | - | - | - | - | - |
| AGE58174.1/1-816putativebifunctionalchitinase/lysozyme\[ParameciumbursariaChlorellavirusNW665.2]/140-305 | R89 | D93 | E95 | E154 | M159 | M161 | D162 | - | - | - | - | - | - | - |
| AGE58174.1/1-816putativebifunctionalchitinase/lysozyme\[ParameciumbursariaChlorellavirusNW665.2]/550-723 | Y94 | D98 | E100 | H159 | M164 | M166 | D167 | - | - | - | - | - | - | - |
| YP_001426411.1/1-805hypotheticalproteinFR483_N779R\[ParameciumbursariaChlorellavirusFR483]/139-304 | R89 | D93 | E95 | E154 | M159 | M161 | D162 | - | - | - | - | - | - | - |
| YP_001426411.1/1-805hypotheticalproteinFR483_N779R\[ParameciumbursariaChlorellavirusFR483]/539-712 | Y94 | D98 | E100 | H159 | M164 | M166 | D167 | - | - | - | - | - | - | - |
| WZO86764.1/1-525chitinase\[O-NE-16]/142-304 | R86 | D90 | E92 | Y151 | M156 | M158 | D159 | - | - | - | - | - | - | - |
| WZO93853.1/1-525chitinase\[OSyNE-4B-S1]/142-304 | R86 | D90 | E92 | Y151 | M156 | M158 | D159 | - | - | - | - | - | - | - |
| WZO91298.1/1-445chitinase\[O-NE-28]/147-309 | R86 | D90 | E92 | H151 | M156 | M158 | D159 | - | - | - | - | - | - | - |
| WZO89577.1/1-445chitinase\[O-NE-24]/147-309 | R86 | D90 | E92 | H151 | M156 | M158 | D159 | - | - | - | - | - | - | - |
| WZO90501.1/1-525chitinase\[O-NE-26]/142-304 | R86 | D90 | E92 | H151 | M156 | M158 | D159 | - | - | - | - | - | - | - |
| WZO77210.1/1-525chitinase\[41-NE-6]/142-304 | R86 | D90 | E92 | H151 | M156 | M158 | D159 | - | - | - | - | - | - | - |
| WZO77693.1/1-525chitinase\[NE-40-1-m]/142-304 | R86 | D90 | E92 | H151 | M156 | M158 | D159 | - | - | - | - | - | - | - |
| WZO78681.1/1-525chitinase\[NE-41-2-m]/142-304 | R86 | D90 | E92 | H151 | M156 | M158 | D159 | - | - | - | - | - | - | - |
| WZO85099.1/1-530chitinase\[O-NE-12]/147-309 | R86 | D90 | E92 | H151 | M156 | M158 | D159 | - | - | - | - | - | - | - |
| YP_009325635.1/1-525Chitinase\[OnlySyngenNebraskavirus5]/142-304 | R86 | D90 | E92 | H151 | M156 | M158 | D159 | - | - | - | - | - | - | - |
| WZO88270.1/1-528chitinase\[O-NE-20]/147-309 | R86 | D90 | E92 | H151 | M156 | M158 | D159 | - | - | - | - | - | - | - |
| WZO92747.1/1-528chitinase\[NE-O-9-L]/147-309 | R86 | D90 | E92 | H151 | M156 | M158 | D159 | - | - | - | - | - | - | - |
| WZO93306.1/1-528chitinase\[OSyNE-4B-L2]/147-309 | R86 | D90 | E92 | H151 | M156 | M158 | D159 | - | - | - | - | - | - | - |
| WZO87136.1/1-528chitinase\[O-NE-17]/147-309 | R86 | D90 | E92 | H151 | M156 | M158 | D159 | - | - | - | - | - | - | - |
| WZO92491.1/1-528chitinase\[NE-O-8-L]/147-309 | R86 | D90 | E92 | H151 | M156 | M158 | D159 | - | - | - | - | - | - | - |
| WZO95440.1/1-528chitinase\[OSyNE-ZA]/147-309 | R86 | D90 | E92 | H151 | M156 | M158 | D159 | - | - | - | - | - | - | - |
| WZO78440.1/1-532chitinase\[NE-41-1-L]/147-309 | R86 | D90 | E92 | H151 | M156 | M158 | D159 | - | - | - | - | - | - | - |
| WZO94180.1/1-539chitinase\[OSyNE-5A-L1]/147-309 | R86 | D90 | E92 | H151 | M156 | M158 | D159 | - | - | - | - | - | - | - |
| WZO84329.1/1-525chitinase\[O-NE-10]/142-304 | R86 | D90 | E92 | H151 | M156 | M158 | D159 | - | - | - | - | - | - | - |
| WZO94943.1/1-525chitinase\[OSyNE-5B-S1]/142-304 | R86 | D90 | E92 | H151 | M156 | M158 | D159 | - | - | - | - | - | - | - |
| WZO93480.1/1-525chitinase\[OSyNE-4B-M2]/142-304 | R86 | D90 | E92 | H151 | M156 | M158 | D159 | - | - | - | - | - | - | - |
| WZO94540.1/1-525chitinase\[OSyNE-5B-M2]/142-304 | R86 | D90 | E92 | H151 | M156 | M158 | D159 | - | - | - | - | - | - | - |
| WZO79465.1/1-530chitinase\[N-NE-4]/147-309 | R86 | D90 | E92 | H151 | M156 | M158 | D159 | - | - | - | - | - | - | - |
| XYW60054.1/1-521chitinase\[NE-P-3-s]/139-304 | R89 | D93 | E95 | E154 | M159 | M161 | D162 | - | - | - | - | - | - | - |
| XYW60732.1/1-521chitinase\[NE-P-4-fs]/139-304 | R89 | D93 | E95 | E154 | M159 | M161 | D162 | - | - | - | - | - | - | - |
| XYW61232.1/1-521chitinase\[NE-P-8-f]/139-304 | R89 | D93 | E95 | E154 | M159 | M161 | D162 | - | - | - | - | - | - | - |
| XYW62772.1/1-521chitinase\[P-NE-12]/139-304 | R89 | D93 | E95 | E154 | M159 | M161 | D162 | - | - | - | - | - | - | - |
| WZO86398.1/1-532chitinase\[O-NE-15]/144-309 | R89 | D93 | E95 | H154 | M159 | M161 | D162 | - | - | - | - | - | - | - |
| XOK27477.1/1-582putativebifunctionalchitinase/lysozyme\[CL-S-1-m]/153-304 | R75 | D79 | E81 | E140 | M145 | M147 | D148 | - | - | - | - | - | - | - |
| AGE49332.1/1-595putativebifunctionalchitinase/lysozyme\[AcanthocystisturfaceaChlorellavirusBr0604L]/166-317 | R75 | D79 | E81 | E140 | M145 | M147 | D148 | - | - | - | - | - | - | - |
| AGE48989.1/1-520putativebifunctionalchitinase/lysozyme\[ParameciumbursariaChlorellavirusAP110A]/139-304 | R89 | D93 | E95 | E154 | M159 | M161 | D162 | - | - | - | - | - | - | - |

| | | | | | | | | | | | | | | |
|---|---|---|---|---|---|---|---|---|---|---|---|---|---|---|
| AGE50008.1/1-520putativebifunctionalchitinase/lysozyme\[ParameciumbursariaChlorellavirusCan18-4]/139-304 | R89 | D93 | E95 | E154 | M159 | M161 | D162 | - | - | - | - | - | - | - |
| XYW62298.1/1-520chitinase\[P-NE-10]/139-304 | R89 | D93 | E95 | E154 | M159 | M161 | D162 | - | - | - | - | - | - | - |
| AGE52349.1/1-523putativebifunctionalchitinase/lysozyme\[ParameciumbursariaChlorellavirusCVR-1]/143-304 | R85 | D89 | E91 | E150 | M155 | M157 | D158 | - | - | - | - | - | - | - |
| YP_009702007.1/1-523putativebifunctionalchitinase/lysozyme\[ParameciumbursariaChlorellavirusCVA-1]/143-304 | R85 | D89 | E91 | E150 | M155 | M157 | D158 | - | - | - | - | - | - | - |
| ABT14345.1/1-520hypotheticalproteinMT325_M791R\[ParameciumbursariachlorellavirusMT325]/139-304 | R89 | D93 | E95 | E154 | M159 | M161 | D162 | - | - | - | - | - | - | - |
| AGE51335.1/1-523putativebifunctionalchitinase/lysozyme\[ParameciumbursariaChlorellavirusCVG-1]/139-304 | R89 | D93 | E95 | E154 | M159 | M161 | D162 | - | - | - | - | - | - | - |
| XYW61920.1/1-523chitinase\[P-NE-9]/139-304 | R89 | D93 | E95 | E154 | M159 | M161 | D162 | - | - | - | - | - | - | - |
| XOK31597.1/1-580putativebifunctionalchitinase/lysozyme\[S-NE-11]/151-304 | R77 | D81 | E83 | E142 | M147 | M149 | D150 | - | - | - | - | - | - | - |
| XOK36306.1/1-580putativebifunctionalchitinase/lysozyme\[GNLD22]/152-304 | R76 | D80 | E82 | E141 | M146 | M148 | D149 | - | - | - | - | - | - | - |
| XOK36264.1/1-582putativebifunctionalchitinase/lysozyme\[LP-F3a-4a]/153-304 | R75 | D79 | E81 | E140 | M145 | M147 | D148 | - | - | - | - | - | - | - |
| XOK31528.1/1-582putativebifunctionalchitinase/lysozyme\[S-NE-10]/153-304 | R75 | D79 | E81 | E140 | M145 | M147 | D148 | - | - | - | - | - | - | - |
| XOK28546.1/1-582putativebifunctionalchitinase/lysozyme\[NES-4B-L1]/153-304 | R75 | D79 | E81 | E140 | M145 | M147 | D148 | - | - | - | - | - | - | - |
| YP_001427295.1/1-595hypotheticalproteinATCV1_Z814L\[Acanthocystisturfaceachlorellavirus1]/166-317 | R75 | D79 | E81 | E140 | M145 | M147 | D148 | - | - | - | - | - | - | - |
| AGE56132.1/1-601putativebifunctionalchitinase/lysozyme\[AcanthocystisturfaceaChlorellavirusMO0605SPH]/172-323 | R75 | D79 | E81 | E140 | M145 | M147 | D148 | - | - | - | - | - | - | - |
| XOK34753.1/1-421putativebifunctionalchitinase/lysozyme\[S-NE-19]/153-304 | R75 | D79 | E81 | E140 | M145 | M147 | D148 | - | - | - | - | - | - | - |
| AGE60252.1/1-601putativebifunctionalchitinase/lysozyme\[AcanthocystisturfaceaChlorellavirusWI0606]/172-323 | R75 | D79 | E81 | E140 | M145 | M147 | D148 | - | - | - | - | - | - | - |
| AGE57134.1/1-601putativebifunctionalchitinase/lysozyme\[AcanthocystisturfaceaChlorellavirusNE-JV-3]/172-323 | R75 | D79 | E81 | E140 | M145 | M147 | D148 | - | - | - | - | - | - | - |
| XOK29995.1/1-582putativebifunctionalchitinase/lysozyme\[NES-5A-S1]/153-304 | R75 | D79 | E81 | E140 | M145 | M147 | D148 | - | - | - | - | - | - | - |
| AGE49664.1/1-595putativebifunctionalchitinase/lysozyme\[AcanthocystisturfaceaChlorellavirusCan0610SP]/166-317 | R75 | D79 | E81 | E140 | M145 | M147 | D148 | - | - | - | - | - | - | - |
| AGE59935.1/1-595putativebifunctionalchitinase/lysozyme\[AcanthocystisturfaceaChlorellavirusTN603.4.2]/189-317 | R52 | D56 | E58 | E117 | M122 | M124 | D125 | - | - | - | - | - | - | - |
| AGE55823.1/1-596putativebifunctionalchitinase/lysozyme\[AcanthocystisturfaceaChlorellavirusMN0810.1]/185-317 | R56 | D60 | E62 | E121 | M126 | M128 | D129 | - | - | - | - | - | - | - |
| XOK35208.1/1-580putativebifunctionalchitinase/lysozyme\[S-NE-22]/149-304 | R79 | D83 | E85 | E144 | M149 | M151 | D152 | - | - | - | - | - | - | - |
| AGE50333.1/1-580putativebifunctionalchitinase/lysozyme\[AcanthocystisturfaceaChlorellavirusCanal-1]/149-304 | R79 | D83 | E85 | E144 | M149 | M151 | D152 | - | - | - | - | - | - | - |
| XOK28216.1/1-580putativebifunctionalchitinase/lysozyme\[NES-4A-S1]/149-304 | R79 | D83 | E85 | E144 | M149 | M151 | D152 | - | - | - | - | - | - | - |
| XOK30852.1/1-582putativebifunctionalchitinase/lysozyme\[S-NE-9]/153-304 | R75 | D79 | E81 | E140 | M145 | M147 | D148 | - | - | - | - | - | - | - |
| XOK34326.1/1-580putativebifunctionalchitinase/lysozyme\[S-NE-18]/152-304 | R76 | D80 | E82 | E141 | M146 | M148 | D149 | - | - | - | - | - | - | - |
| AGE56814.1/1-582putativebifunctionalchitinase/lysozyme\[AcanthocystisturfaceaChlorellavirusNE-JV-2]/153-304 | R75 | D79 | E81 | E140 | M145 | M147 | D148 | - | - | - | - | - | - | - |
| AGE56814.1/1-582putativebifunctionalchitinase/lysozyme\[AcanthocystisturfaceaChlorellavirusNE-JV-2]/153-304 | R75 | D79 | E81 | E140 | M145 | M147 | D148 | - | - | - | - | - | - | - |
| XOK32790.1/1-582putativebifunctionalchitinase/lysozyme\[S-NE-15]/153-304 | R75 | D79 | E81 | E140 | M145 | M147 | D148 | - | - | - | - | - | - | - |
| XOK35891.1/1-582putativebifunctionalchitinase/lysozyme\[S-NE-23]/153-304 | R75 | D79 | E81 | E140 | M145 | M147 | D148 | - | - | - | - | - | - | - |
| XOK33538.1/1-582putativebifunctionalchitinase/lysozyme\[S-NE-16]/153-304 | R75 | D79 | E81 | E140 | M145 | M147 | D148 | - | - | - | - | - | - | - |
| XOK33601.1/1-582putativebifunctionalchitinase/lysozyme\[S-NE-17]/153-304 | R75 | D79 | E81 | E140 | M145 | M147 | D148 | - | - | - | - | - | - | - |
| XOK32004.1/1-582putativebifunctionalchitinase/lysozyme\[S-NE-12]/153-304 | R75 | D79 | E81 | E140 | M145 | M147 | D148 | - | - | - | - | - | - | - |
| XOK35119.1/1-582putativebifunctionalchitinase/lysozyme\[S-NE-20]/175-304 | R53 | D57 | E59 | E118 | M123 | M125 | D126 | - | - | - | - | - | - | - |
| XOK27846.1/1-580putativebifunctionalchitinase/lysozyme\[NES-4A-M1]/152-304 | R76 | D80 | E82 | E141 | M146 | M148 | D149 | - | - | - | - | - | - | - |
| XOK30419.1/1-582chitinase\[S-NE-7]/153-304 | R75 | D79 | E81 | E140 | M145 | M147 | D148 | - | - | - | - | - | - | - |
| AGE59594.1/1-582putativebifunctionalchitinase/lysozyme\[AcanthocystisturfaceaChlorellavirusOR0704.3]/153-304 | R75 | D79 | E81 | E140 | M145 | M147 | D148 | - | - | - | - | - | - | - |
| XOK32390.1/1-589putativebifunctionalchitinase/lysozyme\[S-NE-13]/153-304 | R75 | D79 | E81 | E140 | M145 | M147 | D148 | - | - | - | - | - | - | - |
| AGE57849.1/1-601putativebifunctionalchitinase/lysozyme\[AcanthocystisturfaceaChlorellavirusNTS-1]/172-323 | R75 | D79 | E81 | E140 | M145 | M147 | D148 | - | - | - | - | - | - | - |

**Table S8:** Minimum inhibitory concentration (MIC) of GH18 domains *vs.* yeast and filamentous fungi at different pH.

| | pH 7-8 | | | pH 3–4 | | | |
|---|---|---|---|---|---|---|---|
| **Species** | **AMB** (µg/mL) | **FLC** (µg/mL) | **PBCV1 d2** (µg/mL) | **AMB** (µg/mL) | **FLC** (µg/mL) | **PBCV1 d2** (µg/mL) | **FR483 d2** (µg/mL) |
| *Candida albicans* | 2 | 4 | >1024 | 0,5 | 0,5 | >1024 | >1024 |
| *Candida parapsilosis* | 2 | 2 | >1024 | 0,5 | 4 | >1024 | >1024 |
| *Candida krusei* | 2 | 32 | >1024 | 0,5 | 32 | >1024 | >1024 |
| *Candida auris* | 2 | >64 | >1024 | 2 | 64 | >1024 | >1024 |
| *Cryptococcus neoformans* | 1 | 8 | >1024 | 0,5 | 16 | >1024 | >1024 |
| *Sporothrix brasiliensis* | 2 | >64 | >1024 | 2 | >64 | >1024 | >1024 |
| *Fusarium spp.* | 0,5 | >64 | >1024 | 1 | >64 | >1024 | >1024 |
| *Aspergillus fumigatus* | 1 | >64 | >1024 | 1 | >64 | >1024 | >1024 |

*AMB: Anfotericin B, FLC: Fluconazole
